# Sense-to-sense changes in bacterial genetic codes: Experimental validation by cell-free translation and native-host proteomics

**DOI:** 10.64898/2026.11.10.764470

**Authors:** Paul H. Kim, Megan A. McSweeney, Joshua T. Hong, Joanna P. Stecynska, Hannah Chan, Michael C. Jewett, Yooli K. Light, Kelly P. Williams, Joseph S. Schoeniger

## Abstract

In the sixty years since the publication of the complete genetic code, many variants of the classical codon table have been observed to occur naturally, including both stop codon and sense-to-sense reassignment, but in bacteria the only experimentally validated variants are stop-codon reassignments. Recently, computational analysis with Codetta predicted bacterial genetic codes containing sense-to-sense changes. Using an *Escherichia coli* cell-free protein synthesis (CFPS) system, we tested tRNAs predicted to reassign arginine codons to tryptophan. These candidate CGG-decoding tRNAs from bacteria phylogenetically distant from *E. coli* restored cyan fluorescence from an sfGFP Y66R reporter in a dose-dependent manner when added to CFPS extracts containing canonical tRNA^Arg^(CCG). Reactions lacking the CGG-to-Trp candidate tRNA remained near background. Reassignment was also observed with firefly luciferase and NanoLuc reporters, in which Trp incorporation restored activity and solubility. Intact-protein mass spectrometry yielded masses consistent with Arg-to-Trp substitution. Deep data-independent acquisition (DIA) proteomics of *Anaerococcus prevotii*, a species for which Codetta predicts CGG to code for Trp, revealed six peptides containing Trp encoded at native CGG codons and no peptides containing CGG-encoded Arg, consistent with native CGG decoding as Trp in this organism. Together, these results provide direct functional support for predicted bacterial sense-to-sense genetic codes and establish a cell-free framework for discovering natural tRNAs that expand translational coding capacity.

**GRAPHICAL ABSTRACT:** 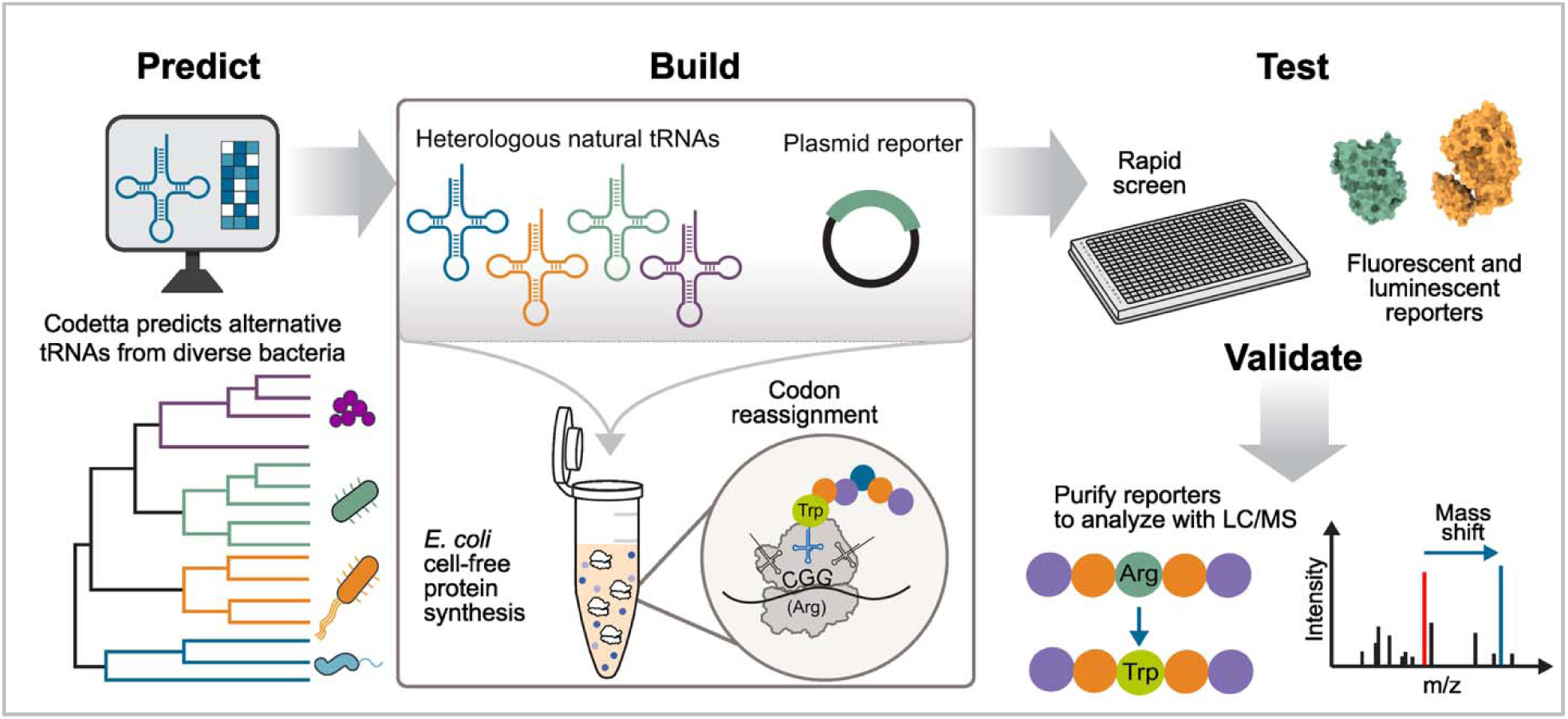

## INTRODUCTION

The classical genetic code [1] is not universal. Natural variants, in which codons are assigned different meanings [2], are of interest for at least two reasons: they show how organisms adapt the protein-coding sequences (CDSs) of an entire genome to a changed code, and they reveal the flexibility of the translational machinery, which synthetic biologists may exploit to expand the genetic code [1–3].

Currently accepted bacterial variants of the standard genetic code (translation table 11) are stop-to-sense changes (Table 1), including the specialized coding of selenocysteine and pyrrolysine [4]. The first variant bacterial code to be discovered, code 4, reassigns the UGA stop codon to Trp and is used by nearly all *Mycoplasmatales* [5, 6]. Code 25, which reassigns UGA to Gly, was subsequently identified throughout the order BD1-5 of the phylum *Patescibacteria* (see note to Table 1) [7]. Both groups have lost the gene for release factor 2 (RF2), which would otherwise compete with the UGA-decoding tRNAs. Some bacteriophages bring additional stop-to-sense changes (UAG-to-Ser or UAG-to-Gln) in their bacterial hosts, frequently carrying the decoding tRNA gene with the recoded stop codons concentrated in late genes [8, 9].

**Table 1.**
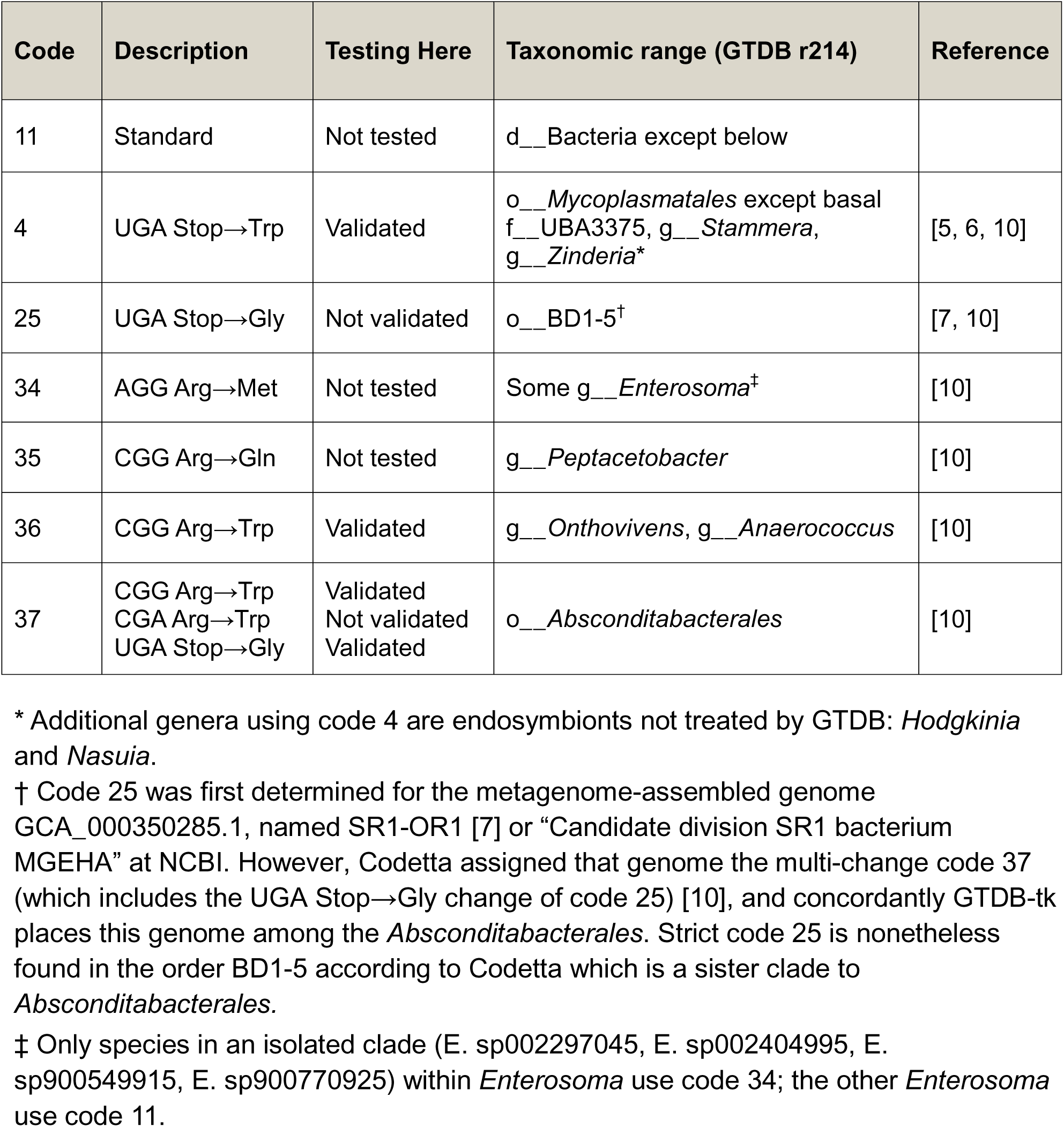
Bacterial genetic codes. Code numbers follow the NCBI list of genetic codes, except for the newly numbered codes 34–37. See Table S1 for a list of alternative tRNAs tested in this study.

Sense-to-sense changes are harder to detect than stop-to-sense changes because their effects on CDSs are more subtle. They are known from eukaryotic nuclear and mitochondrial genomes (Table S2), but had not been proposed for bacteria until Shulgina and Eddy applied their software, Codetta, to diverse bacterial genomes [10]. Codetta infers the most likely amino acid coded by each codon, i.e., the assignment that best preserves conserved amino acids in homologous proteins. It predicted four new bacterial genetic codes with sense-to-sense changes (Table 1), all at codons that normally encode Arg and all supported by analyses of the putative decoding tRNAs. These codes have not been added to the NCBI list because they lack experimental evidence.

Here, we test the predicted Arg-to-Trp assignments (codes 36 and 37) experimentally. If the natural tRNAs identified by Codetta decode CGG as Trp, they should do so in a heterologous translation system, provided that its tryptophanyl-tRNA synthetase charges them. Cell-free protein synthesis (CFPS) systems suit this purpose because candidate tRNAs can be added directly to the reaction [11, 12]. We therefore developed an *E. coli* CFPS platform in which *in vitro*-transcribed candidate tRNAs are added to an extract and screened for their ability to incorporate Trp at CGG codons. We engineered fluorescent and luminescent reporters whose function depends on Trp incorporation. We further verified the amino acid substitution by intact-protein mass spectrometry, measured competition with the endogenous *E. coli* tRNA^Arg^(CCG), and benchmarked the platform against stop-to-sense assignments. Finally, we used proteomics to investigate whether a native code 36 bacterium, *Anaerococcus prevotii*, carries Trp at its CGG codons. Our results show that natural tRNAs from the code 36 (*Anaerococcus* and *Onthovivens*) and code 37 (*Absconditabacterales*) lineages decode CGG as Trp *in vitro*, and that *A. prevotii* proteins contain Trp encoded at CGG codons.

## MATERIALS AND METHODS

### Selection of decoding tRNAs

From our collection of tRNA genes identified among 451, 630 bacterial genomes with taxonomy assigned according to GTDB r214, we focused on those from phylogenetic groups predicted to use nonstandard codes or to closely related outgroups according to GTDB taxonomy [10, 13–15]. For each group, tRNAs carrying the relevant anticodon were clustered by unique sequence, and the most frequently observed sequence was selected for testing. For code 37, an additional UGA-decoding tRNA containing an extended variable region (designated as 37w) was selected from a distinct subgroup.

Candidate tRNAs and their sequences are listed in Table S1. The tRNA nomenclature used in this study specifies the genetic code number, decoded codon, and predicted amino acid. For example, 36a_CGG_W denotes a tRNA from code 36 (*Anaerococcus*) that decodes CGG as tryptophan (W), whereas 36o denotes the corresponding tRNA from *Onthovivens*.

### Plasmid construction

Stop-to-sense decoding was evaluated with an elastin–sfGFP reporter that contains an in-frame UGA codon within the elastin sequence; UGA suppression yields full-length fluorescent protein (Figure S8A). Sense-to-sense decoding was evaluated with sfGFP Y66R reporters in which the chromophore-forming Tyr66 codon was replaced by CGG or CGA (Figure S8B). Incorporation of Arg at position 66 disrupts chromophore maturation and abolishes fluorescence, whereas incorporation of Trp yields cyan fluorescent protein (CFP) fluorescence. Unless stated otherwise, “sfGFP Y66R” denotes the single-CGG reporter sfGFP Y66R R217(CGC), in which the native R217 CGG codon was changed to CGC; the reporter that retains the R217 CGG codon is the two-site reporter (Figure S3). Firefly luciferase reporters were generated by replacing the W417 codon with CGG, and NanoLuc reporters by replacing the W12, W134 or W163 codon with CGG (Figure S8C). These Trp residues are important for correct folding, activity and solubility. Native CGG codons that were not intended as reassignment sites were synonymously changed to CGA, CGT or CGC, preserving the encoded Arg residues. Reporter sequences and target positions are summarized in Figure S8 and in the Supplementary Data.

All reporter constructs were generated with the pJL1-sfGFP plasmid (Addgene plasmid #102634) as the backbone expression vector. The vector was linearized with *Nde*I and *Sal*I high-fidelity restriction enzymes (NEB) and purified with the Monarch DNA Gel Extraction Kit (NEB) before assembly. The elastin–sfGFP coding sequence [17] with 20-bp homology arms for assembly into the vector backbone was synthesized by Twist Bioscience. Linearized vector and inserts were assembled with the NEBuilder HiFi DNA Assembly Kit (NEB) according to the manufacturer’s protocol. Assembled plasmids were transformed into chemically competent *E. coli* cells and plated on selective agar. Several colonies were picked and cultured in LB for plasmid miniprep, and the purified plasmids were verified by DNA sequencing. Firefly luciferase and NanoLuc reporter constructs were generated with the same cloning and assembly strategy. Reporter plasmids were diluted to 100 ng/µL and stored at −20 °C.

### Cell extract preparation

Cell extracts were prepared from *E. coli* BL21 Star (DE3) using a previously described procedure with minor modifications [12, 18]. Cells were streaked from a glycerol stock onto LB agar plates and incubated at 37 °C for 20 h. A single colony was inoculated into 50 mL LB and incubated overnight at 37 °C with shaking at 230 rpm. The next day, 1 L of 2x YTP (16 g/L tryptone, 10 g/L yeast extract, 5 g/L NaCl, 7 g/L dibasic potassium phosphate, and 3 g/L monobasic potassium phosphate) was inoculated with the overnight culture at OD_600_ = 0.05 and grown at 37 °C with shaking at 230 rpm. At OD_600_ = 0.6, IPTG was added to 1 mM to induce expression of T7 RNA polymerase, and cells were grown to OD_600_ = 2.8–3.0. Cells were pelleted by centrifugation at 5, 000 g for 15 min at 4 °C and washed three times with cold resuspension buffer (10 mM Tris-acetate, pH 8.2, 14 mM Mg acetate, 60 mM K acetate, 2 mM DTT). Cell pellets were flash-frozen in liquid nitrogen and stored at −80 °C. The reagents used are listed in Table S3.

Frozen cell pellets were thawed on ice and resuspended in 1 mL resuspension buffer per gram of wet cell mass. Cells were lysed by sonication or, for all ^14^C-leucine incorporation experiments, by high-pressure homogenization; all other CFPS experiments used sonicated extracts. For sonication, 0.7-mL aliquots in 1.7-mL microcentrifuge tubes were sonicated with a 3.175-mm probe on a Q125 Sonicator (Qsonica) at 20 kHz and 50% amplitude, with 5-s pulses separated by 10-s pauses, to a total energy input of 398 J. For homogenization, cells were lysed with a single pass at 20, 000 psi through an Avestin EmulsiFlex-B15 homogenizer. Lysates were clarified by centrifugation at 12, 000 g for 10 min at 4 °C; the supernatant was transferred to a fresh microcentrifuge tube, and the clarification step was repeated until no visible cell debris remained. Protein concentration was measured with the Qubit Protein Broad Range Assay Kit (Invitrogen) according to the manufacturer’s protocol. Extracts were aliquoted for single use, flash-frozen in liquid nitrogen and stored at −80 °C.

### Alternative tRNA *in vitro* transcription and purification

Alternative tRNAs and *E. coli* tRNA^Arg^(CCG) were produced with an RNase P-processing strategy adapted from a previously described protocol [16]. DNA templates encoding the tRNAs were obtained as linear gBlock fragments from Integrated DNA Technologies (IDT) (Table S1). Each construct contained a T7 promoter upstream of the tRNA sequence and constant flanking regions for PCR amplification, where [tRNA] represents the variable tRNA sequence (Table S1): 5’–tttcgccacctctgacttgagcgtcgatttttgtgatgctcgtcaggggggcggagcctatggaaaaacgccagc aacgcgatcccgcgaaattaatacgactcactatagggagaccacaacggtttccctctaga[tRNA] gtcgaccggctgctaacaaagcccgaaaggaagctgagttggctgctgccaccgctgagcaataactagcataacccct tggggcctctaaacgg–3’ Templates were amplified by PCR with Q5 High-Fidelity DNA Polymerase (NEB) according to the manufacturer’s instructions, using the forward primer 5′-TTTCGCCACCTCTGACTTGA-3′ and the reverse primer 5′-CCGTTTAGAGGCCCCAAGGG-3′ for 32 cycles at an annealing temperature of 65 °C. PCR products were purified with the Monarch PCR Cleanup kit (NEB) and verified by 1% agarose gel electrophoresis.

tRNAs were synthesized by *in vitro* transcription with the HiScribe Quick T7 High Yield RNA Synthesis Kit (NEB) according to the manufacturer’s protocol. Reactions were incubated at 37 °C for 20 h and purified with the Monarch RNA Spin Cleanup kit (NEB). tRNA concentrations were measured with the Qubit RNA Broad Range kit (Invitrogen), and tRNAs were stored at −80 °C.

### Cell-free protein synthesis reactions

Cell-free protein synthesis (CFPS) reactions were adapted from an *E. coli* TX-TL system [17]. Reactions (15 µL) were assembled and incubated at 25 °C for 20–24 h unless otherwise specified.

Each reaction contained crude cell-free extract, an energy solution, amino acids, salts, cofactors and plasmid DNA encoding the target protein [18, 19]. The final reaction mixture contained 57 mM HEPES-KOH (pH 7.2), 1.2 mM ATP, 0.85 mM each of GTP, UTP and CTP, 34 µg/mL folinic acid, 170 µg/mL *E. coli* tRNA mixture, 2 mM each of the 20 canonical amino acids, 0.33 mM NAD, 0.27 mM coenzyme A, 1.5 mM spermidine, 1 mM putrescine, 4 mM sodium oxalate, 33 mM phosphoenolpyruvate (PEP), 130 mM potassium glutamate, 10 mM ammonium glutamate and 10 mM magnesium glutamate. For reactions testing alternative tRNAs, 1.65 µM purified tRNA was added to each 15-µL reaction. For tRNAs that depend on RNase P processing, reactions were incubated at room temperature for 2 h before the reporter plasmid was added, to allow processing and maturation of the precursor tRNAs. Reporter plasmid DNA was then added to a final concentration of 6.67 ng/µL (100 ng per 15-µL reaction), and reactions were incubated as indicated.

### Fluorescence and luminescence measurements

CFP and GFP fluorescence were measured with a Tecan Spark microplate reader. After incubation, CFPS reactions were briefly centrifuged to clarify the lysate, and 2 µL of each reaction was diluted into 48 µL nuclease-free water in a clear-bottom black 384-well plate. Fluorescence was read from the bottom of the plate with excitation/emission wavelengths of 430/473 nm (15 nm bandwidth) for CFP and 485/528 nm (10 nm bandwidth) for GFP (sfGFP and elastin–sfGFP). Gain and *z*-position settings were optimized with CFP and sfGFP positive controls and held constant within each experiment. Representative emission spectra are shown in Figure S4.

Firefly luciferase and NanoLuc activities were quantified with the Nano-Glo Dual-Luciferase Reporter Assay System (Promega) according to the manufacturer’s protocol. CFPS reactions expressing luciferase reporters, with or without alternative tRNAs, were incubated at 25 °C for 20 h before analysis and then diluted 1, 000- to 100, 000-fold with distilled water, depending on luminescence intensity. Diluted samples were transferred to a white 384-well flat-bottom microplate and mixed with an equal volume of freshly prepared Nano-Glo assay reagent. Plates were briefly mixed and incubated at room temperature for 1 min to allow the signal to stabilize. Luminescence was measured every 2 min for 30 min with a Tecan Spark microplate reader (0.5-s integration time per well; orbital shaking every 15 s at 3 mm amplitude). Background luminescence from no-DNA control reactions was subtracted from all measurements before analysis.

### Quantification of cell-free protein yields by ^14^C-leucine incorporation

Radioactive incorporation assays were carried out as previously described [18]. CFPS reactions were assembled as above, with the addition of 10 µM ^14^C-leucine, 50 µCi (Revvity) and 15 µL reactions were incubated at 30 °C overnight in 2 mL tubes.

Following overnight incubation, 6 µL of the reaction was mixed with 6 µL of 0.5 M potassium hydroxide (KOH) and incubated for 20 min at 37 °C in PCR tubes to measure total protein. The remainder of the reaction was centrifuged at 14, 000 g for 10 min at 4 °C to pellet the insoluble fraction, and 6 µL of the supernatant (taking care not to disturb the pellet) was mixed with 6 µL of 0.5 M KOH and incubated for 20 min at 37 °C to measure soluble protein. Aliquots (4 µL) of the total-protein and soluble-protein mixtures were spotted onto two filtermats (Revvity) and dried under a heat lamp for at least 20 min. One filtermat was washed three times with 5% (w/v) trichloroacetic acid (Fisher Scientific) for 15 min at 4 °C; the other was not washed. The washed filtermat was dried under the heat lamp for at least 2 h. Both filtermats were placed on a hot plate at 70 °C, and scintillation wax (Revvity) was melted onto each. Radioactivity was counted with a MicroBeta2 (Revvity). Counts from a reaction assembled without template DNA were subtracted from all reactions as background. The concentration of incorporated leucine was calculated by multiplying the fraction of incorporated radioactivity (washed counts/unwashed counts) by the total leucine concentration in the reaction. Protein yield was then calculated by dividing the incorporated leucine concentration by the number of leucine residues in the expressed protein and multiplying by the protein’s molecular weight.

### SDS-PAGE and protein quantification

Protein samples were analyzed on NuPAGE 4–12% Bis-Tris mini protein gels (Thermo Fisher Scientific). After CFPS, samples were briefly centrifuged to pellet the insoluble fraction. A 2-µL aliquot of the clarified lysate was mixed with 4× LDS sample buffer (Invitrogen), TCEP (tris(2-carboxyethyl)phosphine) and water to a total volume of 20 µL, and denatured at 70 °C for 10 min before electrophoresis.

Ten microliters of each prepared sample were loaded and separated in NuPAGE MOPS running buffer (Invitrogen) at 150 V for 45 min. Gels were stained with SimplyBlue SafeStain (Thermo Fisher Scientific) for 1 h at room temperature with shaking, washed three times for 15 min with distilled water, and imaged with an Azure C200 gel imaging workstation. Band intensities were quantified with ImageJ (Figures S1, S2 and S6) [20].

### Protein production and intact-protein mass spectrometry

Control reporter plasmids were transformed into *E. coli* BL21 Star (DE3) via heat shock. Transformed cells were grown overnight at 37 °C on LB agar with appropriate antibiotics. Single colonies were picked and inoculated into 4 mL of LB medium and grown overnight at 37 °C. Subsequently, 50 mL of LB medium was inoculated with the overnight starter culture at an OD_600_ = 0.05. Once the OD_600_ = 0.6, cultures were induced with 1 mM IPTG and grown at 37 °C until the OD_600_ reached 2.0. Cultures were then pelleted at 5, 000 g for 15 min at 4 °C and washed with resuspension buffer (10 mM Tris-acetate (pH 8.2), 14 mM magnesium acetate, 60 mM potassium glutamate, 1 mM TCEP, in Milli-Q water) once. Pellets were weighed, flash-frozen in liquid nitrogen, and stored at −80 °C.

Frozen pellets were thawed on ice and resuspended with resuspension buffer at 1 g / 1 mL, cell pellet mass to buffer ratio. Suspensions were sonicated with a 3.175 mm diameter probe at 20 kHz frequency in 0.7 mL aliquots using 5-sec on / 10-sec off cycles at 50% amplitude. Lysed cells were centrifuged for 20 min at 18, 000 g at 4 °C. Supernatants were collected and filtered through 0.22 µm PVDF membrane (Sigma Aldrich). Filtered lysates were then purified using BIO-RAD NGC Discover Pro FPLC system. Reporter proteins with Strep-tags were captured in the FPLC with Strep-Tactin HP Sepharose 1 mL column. Column was washed with 5 column volumes of binding buffer (100 mM Tris-HCl, 150 mM NaCl, 1 mM EDTA, pH 8.0) and then eluted with 5 column volumes of elution buffer (2.5 mM desthiobiotin, 100 mM Tris-HCl, 150 mM NaCl, 1 mM EDTA, pH 8.0). Elution fractions were collected via Bio-Rad NGC Fraction Collector.

Reporter samples were produced from 200 µL CFPS reactions and purified with Strep-Tactin®XT 4Flow® high-capacity spin column kit (IBA Lifesciences) following manufacturer’s protocol. Purified proteins were stored at −20 °C until use. Elution fractions were verified by SDS-PAGE gel (Figure S9). Intact protein masses were measured using a Waters ACQUITY UPLC system coupled to a Waters SYNAPT G2-Si mass spectrometer operated in positive electrospray ionization mode. Purified proteins were diluted to 0.2 mg/mL in water/acetonitrile (90:10, v/v) containing 0.1% formic acid.

Chromatographic separation was performed using an ACQUITY UPLC Protein BEH C4 column (300 Å pore size, 1.7 µm particle size, 2.1 × 50 mm; Waters) maintained at 70 °C. Mobile phase A consisted of water containing 0.1% formic acid, and mobile phase B consisted of acetonitrile containing 0.1% formic acid. Proteins were eluted using a linear gradient from 20% to 90% mobile phase B over 10 min at a flow rate of 0.06 mL/min. The sample injection volume was 1 µL.

The SYNAPT G2-Si was calibrated in positive-ion mode using sodium iodide over an m/z range of 100–3000. Data were acquired in continuum format and sensitivity mode over m/z 500–2500 using a scan time of 1.0 sec. Representative source settings were as follows: capillary voltage, 3.0 kV; sampling-cone voltage, 40 V; source temperature, 120 °C; desolvation temperature, 350 °C; cone-gas flow, 50 L/h; and desolvation-gas flow, 600 L/h. When lock-mass correction was applied, leucine enkephalin was infused at 10 µL/min, and spectra were corrected using the reference ion at m/z 556.2771.

Raw spectra were acquired and processed using MassLynx version 4.2. Spectra collected across the apex of each protein elution peak were combined, and an adjacent retention-time region was used for background subtraction (Figure S10). Multiply charged ion envelopes were deconvoluted using the MaxEnt1 algorithm (Figures 3C and 4C, D; Figure S11). Deconvolution was performed over mass ranges of 15, 000– 35, 000 Da for NanoLuc and sfGFP and 50, 000–70, 000 Da for firefly luciferase, using an output resolution of 0.5 Da/channel, a uniform Gaussian model with a peak width at half-height of 0.4 Da, and 30 iterations. No additional spectral smoothing was applied. Reported molecular masses correspond to the most abundant neutral deconvoluted species and are presented as average molecular masses (Figures 3C and 4C, D).

**Figure 1.**
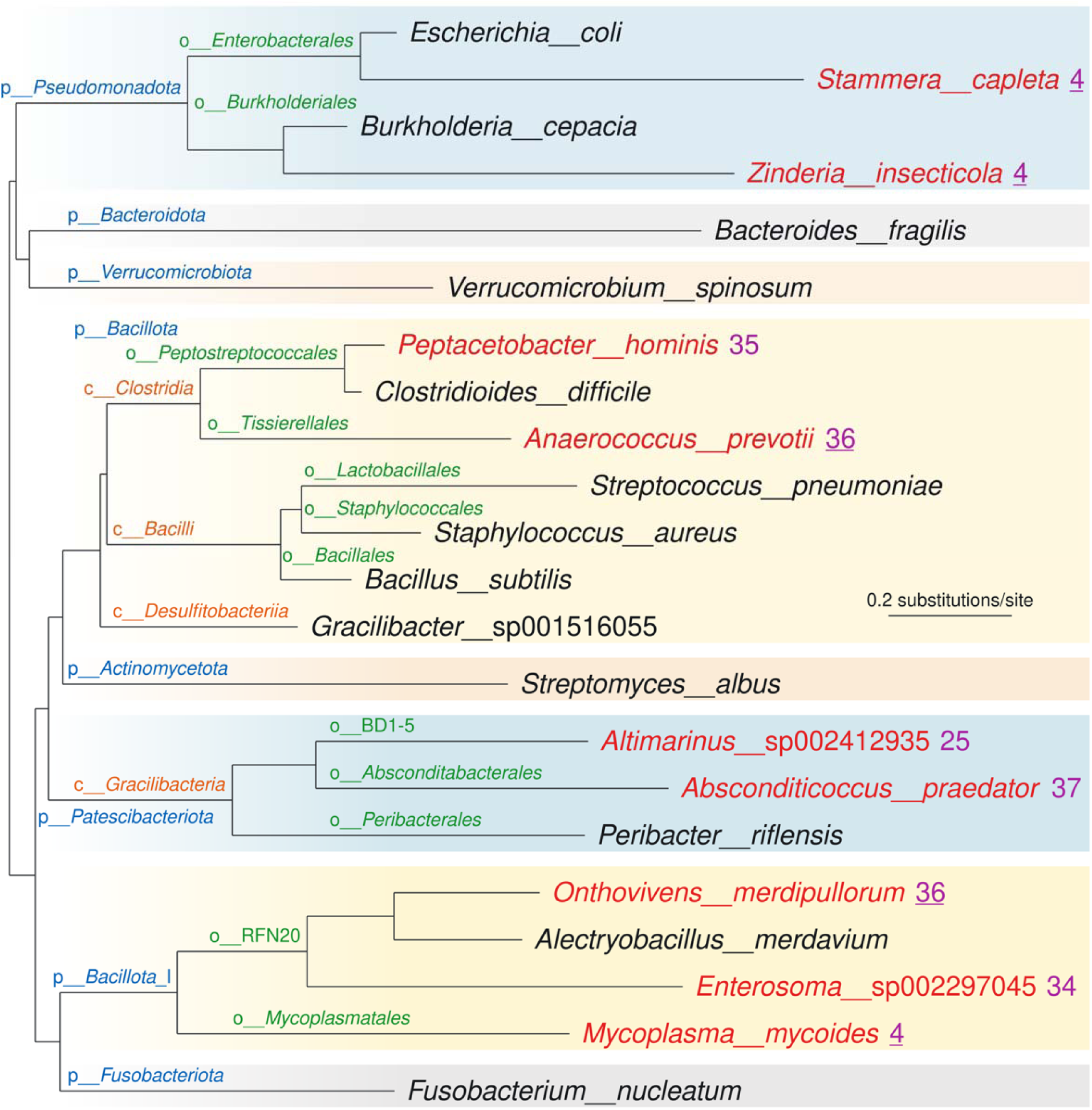
Tree of genetic codes. The bacterial species tree from GTDB release 232 was pruned to a select set of species with PareTree(v.1.0.2). Species are shown in black when they use the standard code 11 and in red when they use a variant code, given to the right of the species name. Underlined code numbers mark codes that may have arisen more than once or through horizontal transfer (codes 36 and 4). Code 4 is also used by the endosymbiont genera Hodgkinia and Nasuia, which were not placed on this tree. Phyla, classes and orders are labeled in blue, orange and green, respectively. Scale bar, 0.2 substitutions per site.

**Figure 2.**
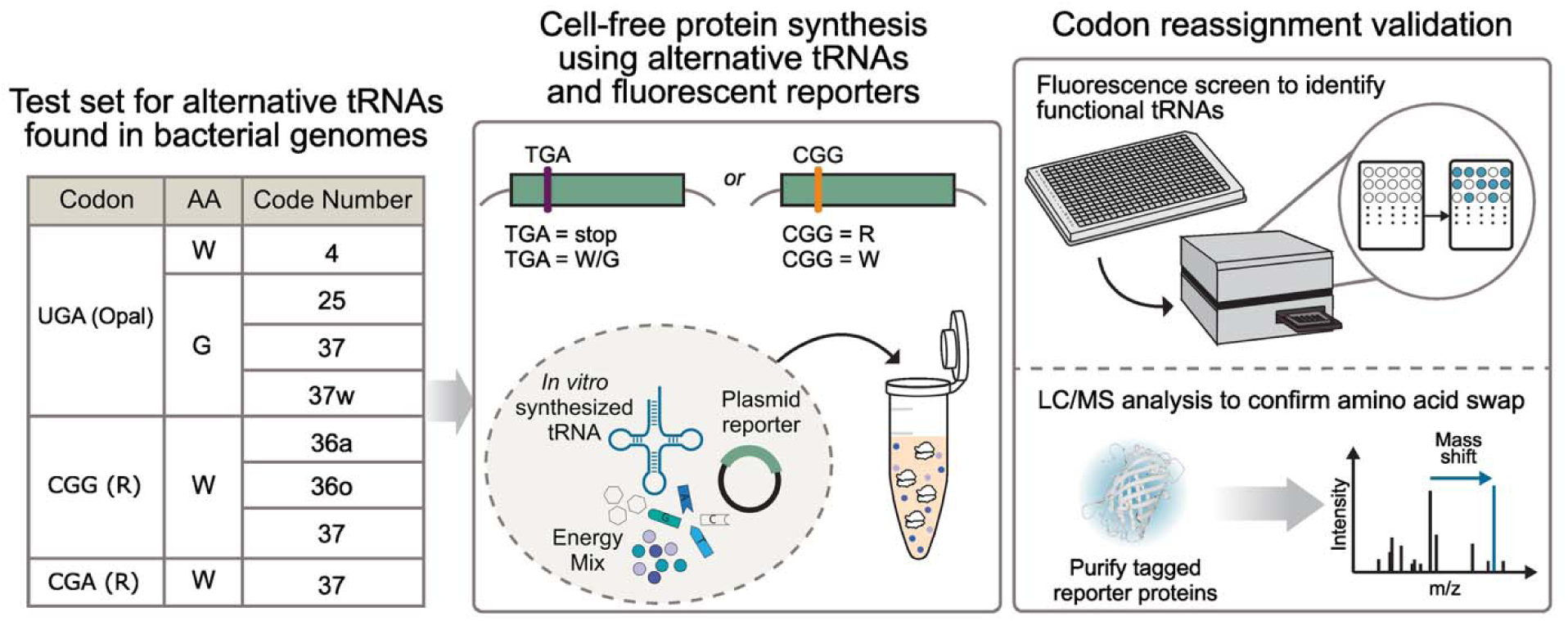
Experimental workflow for identifying and validating tRNA-mediated codon reassignment. Candidate tRNAs predicted by Codetta (left; codon, predicted amino acid and code number) were synthesized *in vitro* and tested in *E. coli* CFPS reactions with two fluorescence reporters: (i) a stop-codon suppression reporter (elastin–sfGFP with an in-frame UGA codon) and (ii) an arginine-to-tryptophan (R to W) substitution reporter based on sfGFP Y66R, in which Trp incorporation at position 66 restores cyan fluorescence. Fluorescence was quantified with a plate reader (top right). Reporter proteins were then purified and analyzed by mass spectrometry (bottom right) to confirm the expected amino acid substitution.

**Figure 3.**
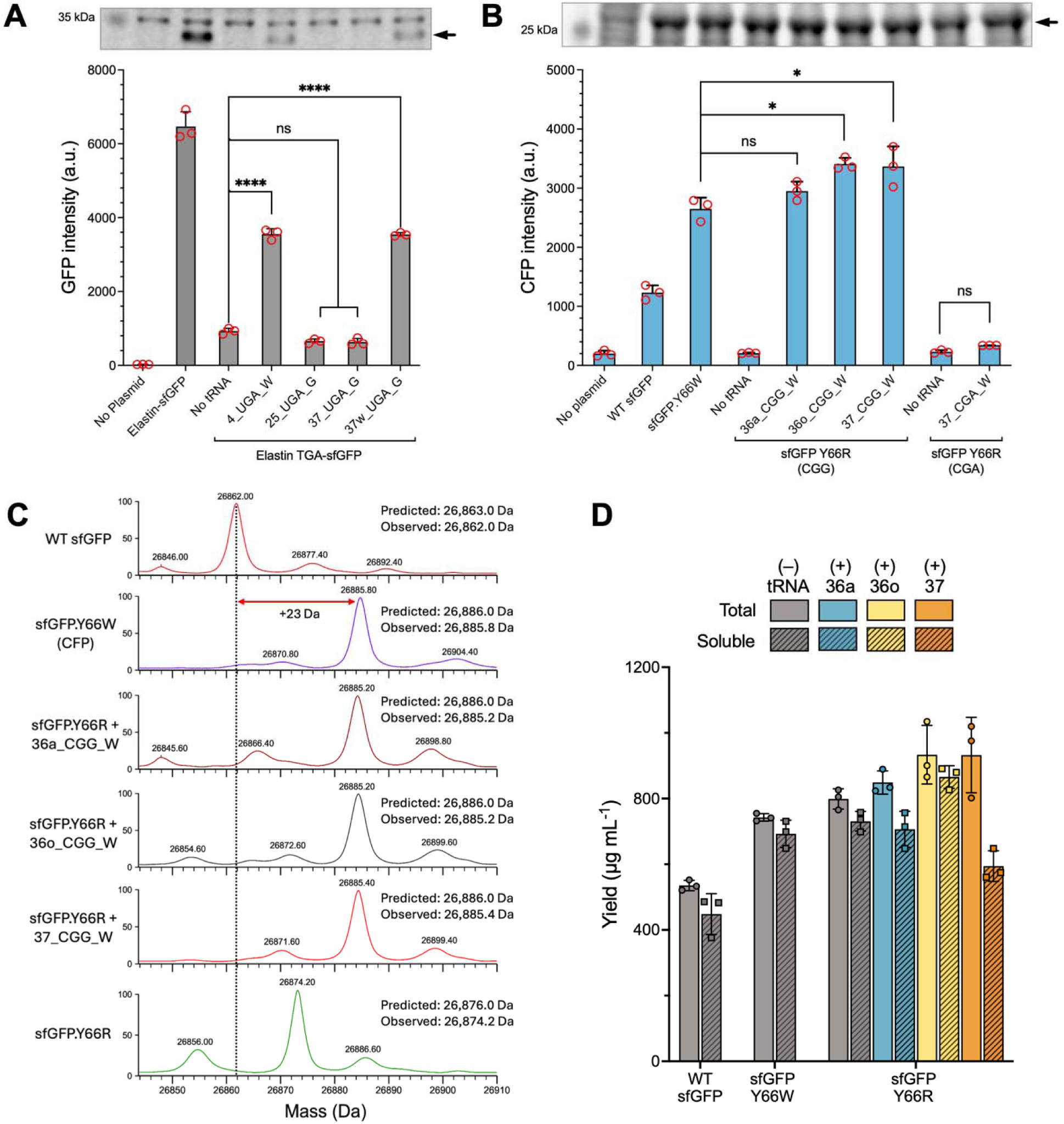
Functional and intact-mass analysis of alternative tRNA-mediated decoding with elastin–sfGFP and sfGFP reporters. (**A**) GFP fluorescence (excitation/emission 485/528 nm) from WT elastin–sfGFP and the elastin–sfGFP UGA reporter with the indicated UGA-decoding tRNAs. The gel above shows CFPS products; the arrow marks full-length elastin–sfGFP (~29.4 kDa). (**B**) CFP fluorescence (430/473 nm) from sfGFP Y66R reporters containing CGG or CGA at position 66, with the indicated alternative tRNAs; the sfGFP Y66W positive control is included. The gel above shows reporter production; the arrow marks sfGFP (~27 kDa). Full gels for A and B are shown in Figure S1. (**C**) Deconvoluted intact-mass spectra of WT sfGFP, sfGFP Y66W and sfGFP Y66R expressed with or without 36a_CGG_W, 36o_CGG_W or 37_CGG_W. Predicted and observed average masses are indicated in the panel. Predicted masses account for removal of the initiator Met and for a loss of ~20 Da for chromophore maturation, which occurs in WT, Y66W and Trp-containing reporters but not in Y66R. The dashed line marks the mass of WT sfGFP; red arrows give the mass difference to WT sfGFP. The y-axis indicates relative intensity (%). (**D**) Total and soluble sfGFP reporter yields quantified by ^14^C-leucine incorporation. Solid bars show total yield and hatched bars show soluble yield. Gray indicates no added alternative tRNA; blue, yellow and orange indicate 36a_CGG_W, 36o_CGG_W and 37_CGG_W tRNAs, respectively. In A, B and D, points show individual measurements and bars show mean ± SD for n = 3 independent CFPS reactions (technical replicates). For the fluorescence comparisons in A and B, statistical significance was determined by one-way ANOVA followed by Dunnett’s multiple comparisons test against the no-tRNA control (A) or Šidák’s multiple comparisons test for the selected pairs shown (B): * P < 0.05, ** P < 0.01, *** P < 0.001, **** P < 0.0001; ns, not significant.

**Figure 4.**
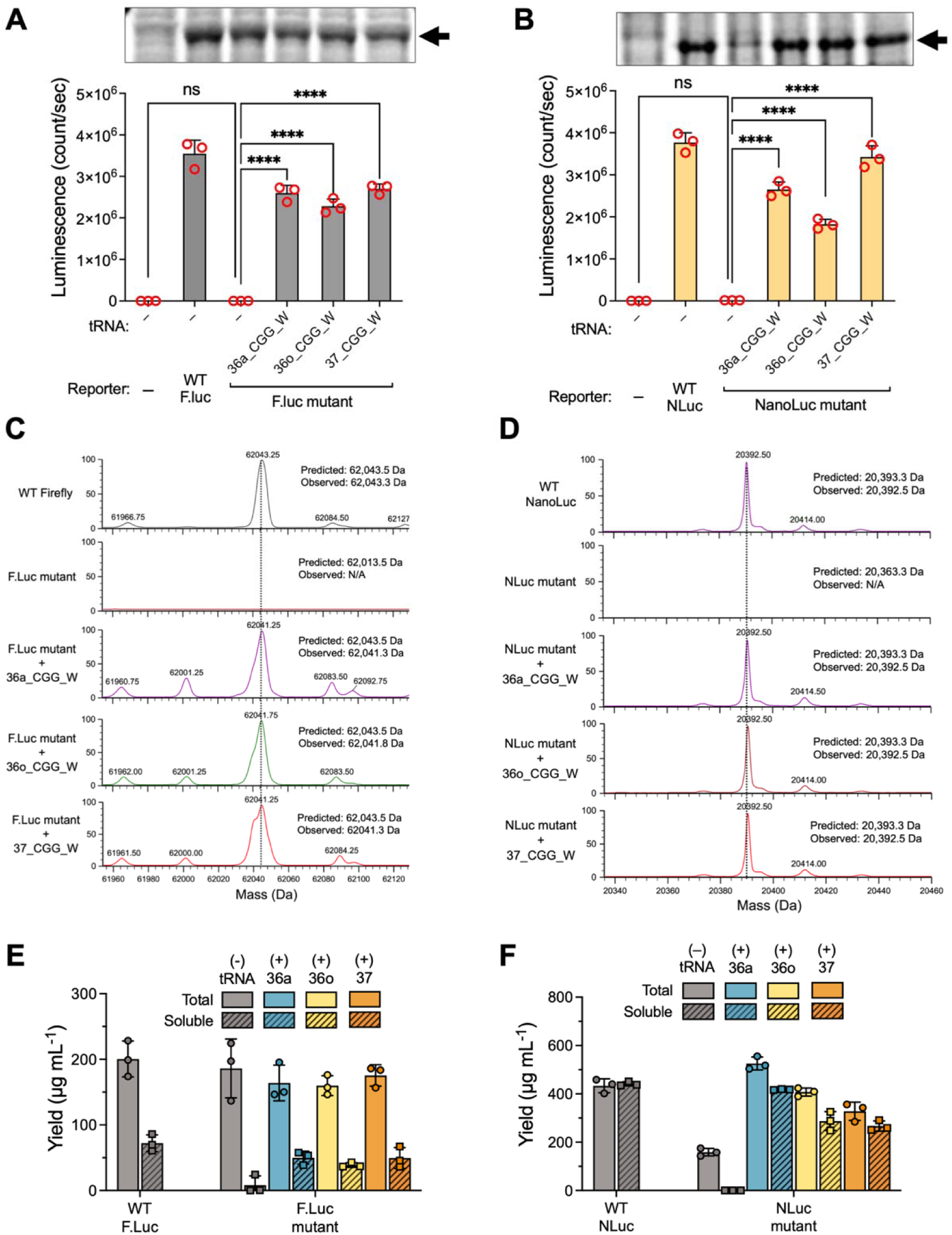
Functional, intact-mass and protein-yield analyses of luciferase reporters with alternative tRNAs. (**A**) Firefly luciferase W417R and (**B**) NanoLuc W12R luminescence after CFPS with the indicated CGG-decoding tRNAs. WT reporters serve as positive controls; mutant reporters without alternative tRNA and no-reporter reactions serve as negative controls. Gels above the plots show reporter production. Arrows mark firefly luciferase (~62 kDa) and NanoLuc (~20.3 kDa); full gels are shown in Figure S2. (**C**) Deconvoluted intact-mass spectra of WT firefly luciferase and W417R expressed without or with 36a_CGG_W, 36o_CGG_W or 37_CGG_W. WT firefly luciferase has a predicted mass of 62, 043.5 Da and an observed mass of 62, 043.3 Da. The W417R mutant has a predicted mass of 62, 013.5 Da, but no intact-protein peak was assigned in the mutant spectrum without tRNA. The tRNA-supplemented samples have observed masses of 62, 041.3, 62, 041.8 and 62, 041.3 Da, respectively. (**D**) Deconvoluted intact-mass spectra of WT NanoLuc and NanoLuc W12R expressed with or without the same alternative tRNAs. Dashed lines in C and D mark the WT mass positions. The y-axis indicates relative intensity (%). (**E**) Firefly luciferase and (**F**) NanoLuc total and soluble protein yields quantified by ^14^C-leucine incorporation. Solid bars show total yield and hatched bars show soluble yield. Gray indicates no added alternative tRNA; blue, yellow and orange indicate 36a, 36o and 37 tRNA, respectively. In A, B, E and F, points show individual measurements and bars show mean ± SD for n = 3 independent CFPS reactions (technical replicates). For luminescence comparisons in A and B, statistical significance was determined by one-way ANOVA followed by Dunnett’s multiple comparisons test against the mutant reporter without alternative tRNA: * P < 0.05, ** P < 0.01, *** P < 0.001, **** P < 0.0001; ns, not significant.

### Anaerococcus prevotii proteomics

The type strain of *A. prevotii* (strain PC 1; ATCC 9321 = DSM 20548) was purchased from ATCC and revived through liquid growth in Chopped Meat Glucose Broth, CMG, (Anaerobe systems, AS-813) and plated on Brucella Blood agar plates (Anaerobe systems, AS-111). The strain was then grown in its standard culture medium, Peptone Yeast Extract Broth with Glucose (Anaerobe systems, AS-822), which is equivalent to DSMZ Medium 104 recommended for its standard growth. The strain was cultured in liquid for ~72 h. All culture steps were done at 37 °C, in an anaerobic chamber (Don Whitley Scientific, A35, gas mixture =5% CO2, 5% H2, 90% N2). The cells were harvested by centrifugation (8, 000 g, 10 min, 4 °C); the pellets were washed thrice in PBS, then flash-frozen in liquid nitrogen. Next, cell pellets were analyzed by Novogene (project X202SC26086554-Z01; sample A_prevotii_1; n = 1) for protein extraction and quantification, tryptic digestion, peptide desalting and data-independent acquisition (DIA) proteomics.

Peptides were analyzed on an Orbitrap Astral mass spectrometer (Thermo Fisher Scientific) coupled to a Vanquish Neo UHPLC system in trap-and-elute mode (0.5 cm × 300 µm trap column; 15 cm × 150 µm analytical column; 60-min method, 24 samples per day). Peptides were separated with a gradient of buffer B (80% acetonitrile, 0.1% formic acid) in buffer A (0.1% formic acid in water) from about 8% to 22.5% B over 42 min and to 35% B at 52.8 min, followed by a wash to 99% B, at 0.5 µL/min. The ion source was operated at 2.0 kV with the ion-transfer tube at 290 °C. MS1 spectra were acquired in the Orbitrap at a resolution of 240, 000 (m/z 380–980; maximum injection time 5 ms; normalized AGC target 500%). DIA MS2 spectra were acquired in the Astral analyzer with 300 non-overlapping 2-Th isolation windows covering m/z 380–980, acquired as three interleaved sets of 100 windows (one MS1 scan per set; cycle time 0.55 s, so that each window was sampled about every 1.6 s), HCD at a normalized collision energy of 25%, a scan range of m/z 150–2, 000, a maximum injection time of 3 ms and an AGC target of 50, 000. Instrument settings were read from the raw-file method header (Table S6).

The *Anaerococcus prevotii* DSM 20548 chromosome (GenBank CP001708.1) contains 1, 702 annotated CDSs corresponding to 1, 697 unique protein sequences. Four transposase loci (Apre_0251, Apre_0729, Apre_1194, and Apre_1360) encode proteins identical to that encoded by Apre_0040, while the elongation factor Tu encoded by Apre_1140 is identical to that encoded by Apre_0460. The canonical search database retained one representative of each unique protein sequence, excluding these five redundant accessions without losing unique sequence information. Addition of 37 CGG-to-Trp variant sequences yielded a combined search database containing 1, 734 entries. For the 37 proteins with CGG codons, both the CGG→Arg and CGG→Trp variant forms were included in the database. The 37 alternative proteins represented 39 predicted CGG→Trp sites; 35 proteins contained one site, while ACV28313.1 and ACV29475.1 each contained 2 sites.

Raw data were processed with DIA-NN v1.8.1 with trypsin specificity, up to two missed cleavages, carbamidomethylation of cysteine as a fixed modification, N-terminal methionine excision as a variable modification and fragment-ion quantification, at a precursor false-discovery rate of 1% (Q.Value and Global.Q.Value ≤ 0.01) [21]. The precursor and protein-group matrices (report.pr_matrix.tsv and report.pg_matrix.tsv) were used for the analyses below.

Peptides were assigned to CGG sites by exact matching to the reference and variant sequences. For each site we recorded (i) whether a Trp-containing peptide spanning the site was identified and (ii) whether an Arg-form partner peptide was identified. A partner peptide is a fully cleaved tryptic peptide of 7–30 residues that ends at, or begins immediately after, the residue at the site and therefore exists only if trypsin cleaves after that residue (that is, if it is Arg); Trp-form peptides were defined analogously. Peptides outside 7–30 residues were considered invisible, as the identified peptides span 7–30 residues (Figure S24). The detection probability of a visible peptide in a given protein was estimated as the fraction of its fully cleaved 7–30-residue tryptic peptides that were identified, excluding site-adjacent peptides. The expected number of detected peptides under each hypothesis is the sum of these probabilities, and the probability of the observed count follows the Poisson-binomial distribution (independence assumed). The expected number of false-positive Trp-form identifications was estimated as 1% of the identified precursors multiplied by the fraction of *in silico* library precursors (7–30 residues, ≤2 missed cleavages, charge 2–4) that are Trp-form variant peptides (151 of 125, 492 peptides). Codon usage was calculated from the 1, 702 annotated CDSs (555, 757 codons); tRNA genes were taken from the GenBank annotation and aligned by global pairwise alignment. Analyses used custom Python scripts (PRIDE dataset identifier PXD086557).

*Raw-data inspection.* The Thermo raw file was read with ThermoRawFileParser (v2.0.0-dev, macOS arm64 build) and analyzed with custom Python scripts [22].

Because retention times from the DIA-NN report were not available, the elution of each Trp-form peptide was located directly from the MS2 scans. For every MS2 scan whose 2-Th isolation window contained the precursor, the b and y ions of the peptide (charge 1+ and 2+, m/z ≥ 150; b1 and y1 excluded) were matched to the nearest peak within 5 ppm, and every window of five consecutive such scans (about 8 s) was scored as the sum, over scans, of the number of matched ions above three. The same score was calculated for 198 decoy fragment lists made by adding 3 to 200 times 1.000495 Da (divided by the charge) to the fragment masses, which preserves the mass defect. The family-wise probability is (k + 1)/(198 + 1), where k is the number of decoy lists whose highest window score anywhere in the 60-min run was at least as high as the observed score. Extracted-ion chromatograms of the matched ions, the MS1 intensity at the monoisotopic m/z and its M+1 and M+2 isotope peaks (±10 ppm; sum of five MS1 scans around the MS1 maximum within 6 s of the cluster center), and the highest-scoring MS2 spectrum were then inspected (Figures S26 and S27; Table S8).

### Analysis of tRNA competition data

A simple kinetic model of protein yield as representative of the average rate of production of protein gives the relationship *S_W_* / *S_R_* = (*k_W_ tRNA_W_)*/(*k_R_ tRNA_R_*) where *S* is the instantaneous ribosomal CGG site occupancy by one or the other tRNA (which under our average rate assumption is proportional to the total protein of each type produced), *k* is the respective effective on rate, and *tRNA* is the concentration.

Assuming that the total number of ribosomal CGG sites *S* is limiting, that the mRNA and tRNAs are always in excess, and that *K* = *k_W_* / *k_R_* is a dimensionless ratio of the effective on rates (*K* can be viewed loosely as a relative efficiency of the two tRNAs). We assume for this model that the *tRNA_W_* charges exclusively with tryptophan.

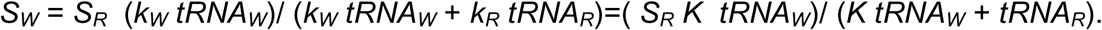

We note that *tRNA_R_* is actually the sum of basal levels in CFE *tRNA_Rbase_* present in the CFE plus added *tRNA_Radd_*.

For purposes of fitting the data, we note that the signal measured is proportional to *S_W_* but depends on various assay conditions that turn it into a measured fluorescence intensity, and may include a background. Lacking the constraints necessary to turn this into molar detectivity, we lump the parameters *S_R_ K* as *F*, and add a background term *B* to get an expression that can be directly fitted to data:

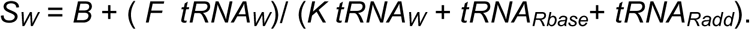

Nonlinear regression curve fits were performed using the Mathematica (version 14.3) function NonlinearModelFit, with the four parameters *B, F, K* and *tRNA_Rbase_*fitted to the above hyperbolic expression for *S_W_* : *B* = 426, *F* = 1950, *K* = 0.27, and *tRNA_Rbase_* = 0.37.

### Statistical analysis

Statistical analyses were performed as specified in the figure legends. Unless otherwise indicated, data are presented as mean ± SD of at least three independent CFPS reactions prepared from the same extract batch (technical replicates). Groups were compared by one-way ANOVA followed by Dunnett’s multiple comparisons test when each group was compared with a single control group (Figures 3A and 4A, B) or by Šidák’s multiple comparisons test when selected pairs were compared (Figures 3B and 5); P values were adjusted for multiple comparisons. Analyses and graphs were prepared with Prism 10 (GraphPad Software); Excel version 16 (Microsoft) was used for data organization.

**Figure 5.**
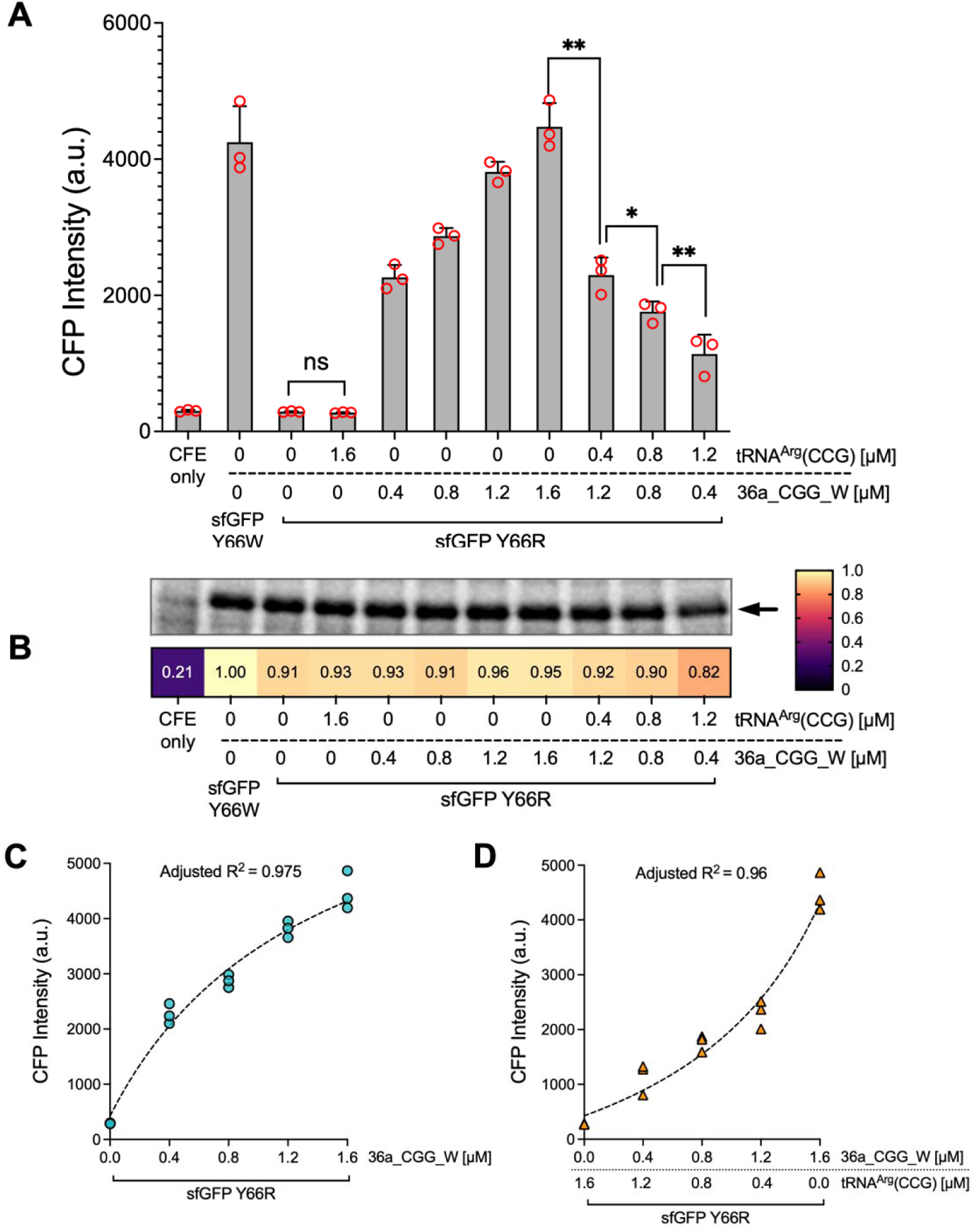
Competition between the alternative tRNA 36a_CGG_W and endogenous ***E. coli* tRNA^Arg^(CCG) at the CGG codon.** (**A**) CFPS reactions expressing the sfGFP Y66R reporter were supplemented with 36a_CGG_W and *E. coli* tRNA^Arg^(CCG) (labeled as *E. coli*_R) at the indicated concentrations. Mixed-tRNA conditions contained a constant total of 1.6 µM added tRNA; 36a_CGG_W-only conditions were included at the corresponding alternative-tRNA doses. Controls include extract only (CFE only), sfGFP Y66W (positive control), no added tRNA, and tRNA^Arg^(CCG) only. CFP fluorescence decreased as the proportion of canonical tRNA increased. Red circles show individual measurements, and bars show mean ± SD for n = 3 independent CFPS reactions (technical replicates). Brackets indicate comparisons of selected pairs, including mixed-tRNA conditions with the same total tRNA, by one-way ANOVA followed by Šidák’s multiple comparisons test: * P < 0.05, ** P < 0.01; ns, not significant. (**B**) Coomassie-stained SDS–PAGE analysis of sfGFP produced in CFPS reactions under the conditions shown in (A). The arrow indicates the sfGFP band (~27 kDa). The heat map shows densitometric quantification of sfGFP. Band intensities were normalized to the total protein signal in each lane and expressed relative to the sfGFP Y66W control (set to 1). (**C**) The figure shows the nonlinear regression curve fit plotted for the data in panel A, where there was no added tRNA^Arg^(CCG). The adjusted R^2^ was 0.975. (**D**) The graph shows the fit for the data when the total amount of added tRNA^Arg^(CCG) + 36a_CGG_W tRNA was maintained at 1.6 µM. The adjusted R^2^ was 0.96, showing acceptable goodness of fit.

### Phylogenetic analysis

Protein sequences were aligned with MAFFT v7.526 using default parameters [23]. Maximum-likelihood trees were inferred with RAxML-NG [24] using the LG+G8+F model [25], 10 parsimony starting trees and 200 bootstrap replicates. Ingroup and outgroup genomes were defined by their GTDB taxonomy and Codetta assignment.

## RESULTS

### Phylogenetic analysis of Codetta predicted sense-to-sense genetic code changes in bacteria

Codetta identified several candidate bacterial sense-to-sense reassignments; although it would have been able to detect these at any sense codon, all involving affected codons canonically assigned to arginine (Figure 1; Table 1)[10, 26]. The alternative assignments improved the conservation of amino acids at homologous protein positions and were supported by candidate decoding tRNAs. Because not all genomes in our expanded collection were included in the original Codetta analysis, we also analyzed closely related genomes that had not yet been analyzed. This analysis supported assignment of code 36 to s JALEKO01 sp022769085, whose branch subtends the *Anaerococcus* group in the species tree.

Inspection of the tRNA complements identified candidates with anticodons complementary to the reassigned codons and with sequence features consistent with altered aminoacylation identity. Representative tRNAs from the predicted code groups and from closely related variants were selected for functional testing (Table S1).

### An *E. coli* cell-free platform detects stop-to-sense decoding by natural tRNAs

We established an *E. coli* CFPS platform for rapid functional testing of Codetta-predicted tRNAs (Figure 2). In this platform, candidate tRNAs are transcribed *in vitro*, added to an extract-based CFPS reaction, and tested with reporters whose output depends on decoding of an engineered codon. Reaction temperature, incubation time, Mg^2+^ concentration, reporter plasmid concentration and supplemented tRNA concentration were optimized before screening (Figures S5–S7), and the selected conditions were used in subsequent experiments unless otherwise specified.

We first tested stop-to-sense decoding with an elastin–sfGFP reporter, which contains an in-frame UGA codon within the elastin sequence (Figure S8A) [27]. With canonical decoding, translation terminates at the engineered stop codon, and sfGFP fluorescence is lost. Decoding of UGA by the Trp-incorporating suppressor tRNA (4_UGA_W) or by the Gly-incorporating suppressor tRNA (37w_UGA_G) restores full-length reporter production and sfGFP fluorescence. Production of the full-length reporter was also detected by Coomassie-stained SDS-PAGE (Figure 3A; Figure S1A).

Reactions lacking a supplemented suppressor tRNA retained, on average, 15.2% of the fluorescence of the WT elastin–sfGFP control. This background is consistent with low-level UGA readthrough in *E. coli*, which can result from near-cognate decoding and incomplete termination [2]. In CFPS reactions, competition between endogenous tRNAs and release factors may allow a small fraction of ribosomes to read through the engineered stop codon, producing detectable full-length reporter even without a suppressor tRNA [28, 29].

Among the tested candidates, the 4_UGA_W and 37w_UGA_G tRNAs restored GFP fluorescence substantially above the no-tRNA control, reaching on average about 55% of WT elastin–sfGFP, whereas the 25_UGA_G and 37_UGA_G tRNAs produced fluorescence comparable to the no-tRNA control (Figure 3A). Suppression efficiency therefore varied substantially among the Codetta-predicted UGA-decoding tRNAs. The inactivity of the 25_UGA_G tRNA in this assay is not evidence against code 25, which was established in the native context and by co-expression of the SR1 glycyl-tRNA synthetase with its cognate tRNA in *E. coli* [7]; it may instead reflect incompatibility with the *E. coli* glycyl-tRNA synthetase or competition with release factors in the extract. The 37w_UGA_G tRNA is a rarer variant among the proposed UGA-decoding tRNAs of the *Absconditabacterales*; it bears a significantly extended variable loop with a stem–loop structure that may improve decoding [10]. Collectively, these assays show that several natural tRNAs remain functional in a heterologous bacterial translation system and can alter decoding outcomes at defined codon positions.

### Natural CGG-decoding tRNAs from codes 36 and 37 restore Trp-dependent fluorescence

To test the predicted Arg-to-Trp reassignments directly, we developed a fluorescence-based reporter assay with engineered sfGFP variants. The assay exploits the essential role of residue 66 in chromophore formation: substitution of Tyr66 with Trp converts sfGFP into a cyan fluorescent protein (CFP) with a blue-shifted emission spectrum (Figure S4) [30–32]. We replaced the native Tyr66 codon with an Arg codon, either CGG or CGA to test CGG- and CGA-decoding tRNAs, respectively, so that canonical incorporation of Arg at position 66 prevents chromophore formation and abolishes fluorescence, whereas successful Arg-to-Trp reassignment restores CFP fluorescence (Figure 3B).

Under canonical decoding conditions, the sfGFP Y66R reporters gave only background fluorescence (Figure 3B). Supplementation with the candidate CGG-decoding tRNAs from codes 36a, 36o and 37 restored CFP fluorescence to 111.4 ± 6%, 128.7 ± 3.9% and 127.1 ± 12.9%, respectively, compared to the Y66W CFP positive control (Figure 3B). The signal generated with the 36a_CGG_W tRNA was not significantly different from that of the Y66W positive control, whereas the 36o_CGG_W and 37_CGG_W tRNAs produced significantly higher CFP signals. Because fluorescence also depends on protein yield, folding and chromophore maturation, values above 100% of the Y66W control do not imply more-than-complete decoding (see Discussion). In contrast, the candidate CGA-decoding tRNA from code 37 (37_CGA_W) failed to restore CFP fluorescence above background. Together, these results indicate that at least three distinct natural tRNAs predicted by Codetta decode CGG as Trp in the *E. coli* CFPS system.

### Intact-protein mass spectrometry is consistent with Arg-to-Trp substitution

To confirm that CFP recovery reflects Arg-to-Trp substitution rather than indirect effects on reporter expression or chromophore maturation, we analyzed Strep-tag-purified reporter proteins by intact-protein mass spectrometry (Figure 3C). The sequence-derived mass of Strep-tagged sfGFP is 26, 883 Da, but sfGFP variants that can form the chromophore (T65–Y66–G67 in the WT) undergo post-translational cyclization, dehydration and oxidation, with a net mass loss of approximately 20 Da [33]; the predicted mass of mature WT sfGFP–Strep is therefore 26, 863 Da. As expected, WT sfGFP (observed 26, 862.0 Da) and the sfGFP Y66R reporter, which cannot form the chromophore (observed 26, 874.2 Da), had masses close to their predicted values of 26, 863 and 26, 876 Da, respectively. The sfGFP Y66W positive control showed the expected increase of 23 Da relative to WT sfGFP (observed 26, 885.8 Da; predicted 26, 886 Da), consistent with replacement of Tyr by Trp at position 66.

The sfGFP Y66R reporter carries an engineered CGG codon at position 66, the site predicted to undergo reassignment (Figure S8B). Substitution of Arg by Trp adds 30 Da to the polypeptide, but Trp, unlike Arg, permits chromophore maturation, which removes ~20 Da. The expected net shift of the mature, Trp-containing protein relative to the Y66R reporter is therefore +10 Da (predicted masses 26, 886 and 26, 876 Da).

Reporter proteins synthesized with the 36a, 36o or 37 tRNA showed a dominant species at ~26, 886 Da (36a, 26, 885.2 Da; 36o, 26, 885.2 Da; 37, 26, 885.4 Da), about 11 Da above the Y66R reporter and indistinguishable from the Y66W control (26, 886 Da), whereas the reporter synthesized without added tRNA retained the mass expected for the Arg-containing protein (26, 874.2 Da; Figure 3C).

### Reassignment generalizes to firefly luciferase and NanoLuc reporters

To test whether reassignment generalizes across protein scaffolds and sequence contexts, we developed two enzymatic reporters based on firefly luciferase and NanoLuc. Compared with fluorescence reporters, luciferase reporters offer a broad dynamic range, low background and sensitive quantification of partially restored protein activity [34, 35].

We selected conserved Trp residues that contribute to substrate binding, structural stability or catalysis as reassignment targets: W417 in firefly luciferase and W12, W134 and W163 in NanoLuc (Figure 4A, B; Figures S8C and S12) [36, 37].

Replacing these Trp codons with CGG (Arg) was expected to impair activity substantially, providing a sensitive readout of Trp incorporation through restored function [34, 38]. Among the individual NanoLuc CGG substitutions, W12R caused the strongest reduction in luminescence and in soluble-protein yield (Figure S12), and was therefore used for subsequent reassignment assays. To confine reassignment to the intended sites, endogenous CGG codons in the firefly luciferase and NanoLuc coding sequences were synonymously mutated to CGA or CGT, which preserves the native protein sequence while eliminating potential off-target decoding (Figure S8C).

Under canonical decoding conditions, Arg codons at these critical Trp positions strongly reduced luciferase activity: both mutant reporters gave near-background luminescence without alternative tRNA (Figure 4A, B). Supplementation with any of the three candidate CGG-decoding Trp tRNAs significantly increased reporter activity (P < 0.0001). The 36a_CGG_W, 36o_CGG_W, and 37_CGG_W tRNAs restored firefly luciferase activity to approximately 73%, 64% and 76% of the WT level, and NanoLuc W12R activity to approximately 70%, 48% and 91% of the WT level, respectively.

Rescue efficiency varied between reporters and tRNAs (Figure 3B, 4A, B). Differences in recovery between reporters and codon positions suggest that local sequence context, translational dynamics, and kinetic competition between endogenous Arg tRNAs and the alternative tRNAs may influence reassignment efficiency.

Next, intact-protein mass spectrometry of the tRNA-supplemented CFPS reactions was performed and showed the mass of the WT protein (Figure 4C, D); no intact-protein peak could be assigned for the mutant reporters made without alternative tRNA, consistent with their absence from the soluble fraction (Figure 4E, F). These results show that the alternative tRNAs mediate site-specific reassignment in multiple independent reporters, arguing against a reporter-specific artifact.

To determine whether changes in reporter activity reflect differences in protein yield or solubility, we measured the total and soluble yields of the sfGFP, NanoLuc and firefly luciferase reporters, first qualitatively by Coomassie-stained SDS-PAGE (Figures S1 and S2) and then quantitatively by ^14^C-leucine incorporation (Figures 3D and 4E, F). For sfGFP, the Y66R reporter showed total yields comparable to the control constructs, and the alternative tRNAs did not reduce overall protein production (Figure 3D). Soluble yields were generally comparable to that of the Y66W control.

The effect of codon substitution on solubility depended on the reporter and the substituted site. The W12R substitution in NanoLuc reduced total protein production relative to WT and, more strikingly, yielded almost no soluble protein without alternative tRNA. Addition of the 36a_CGG_W, 36o_CGG_W or 37_CGG_W tRNA restored the soluble fraction of NanoLuc W12R (Figure 4F), consistent with Trp incorporation by all three tRNAs. For firefly luciferase W417R, each of the three tRNAs partially restored the soluble fraction (Figure 4E). Taken together, the restoration of solubility provides an additional, activity-independent readout of Trp incorporation for the variants whose solubility requires Trp.

### The supplemented alternative tRNA competes with endogenous tRNA^Arg^(CCG)

To determine to what extent supplemented alternative tRNAs compete with endogenous *E. coli* tRNAs during translation, we performed titration experiments with the sfGFP Y66R CFP reporter. Because the cell-free extract retains canonical tRNA^Arg^(CCG) that can decode CGG codons, alternative tRNAs must compete for accommodation at the ribosomal A site to mediate reassignment. The endogenous tRNA^Arg^(CCG) is a low-abundance *E. coli* Arg isoacceptor: absolute measurements place it at approximately 1.5–2.6 µM, corresponding to ~0.6–1.2% of the total tRNA pool depending on growth condition [39], and more recent RNA-seq-based estimates report ~2% of the total tRNA signal [40].

Increasing concentration of the 36a_CGG_W tRNA produced a dose-dependent increase in fluorescence from the sfGFP Y66R reporter, consistent with progressive redirection of CGG decoding toward Trp (Figure S7A). Of the concentrations tested (0, 0.16, 0.33, 0.83, 1.65, 2.15 and 3.3 µM), 1.65 µM gave the maximum CFP signal, which did not increase at higher concentrations. These results are consistent with competitive decoding, in which the relative abundance of competing tRNA species influences codon recognition and decoding outcomes [41, 42].

To evaluate competition directly, we supplemented purified endogenous tRNA^Arg^(CCG) into reactions containing the reassignment-capable 36a_CGG_W tRNA, while keeping the total added tRNA constant at 1.6 µM per 15-µL reaction to avoid effects of tRNA crowding. Endogenous tRNA^Arg^(CCG) alone yielded only background fluorescence. Increasing the competitor concentration to 0.4, 0.8 and 1.2 µM, while decreasing 36a_CGG_W to 1.2, 0.8 and 0.4 µM, respectively, reduced CFP intensity to approximately 51.4 ± 5.8%, 39.4 ± 3.4% and 25.5 ± 6.4% of the signal obtained with 1.6 µM of 36a_CGG_W alone (mean ± SD, n = 3; Figure 5A). These mixtures also produced lower fluorescence than reactions containing the corresponding concentrations of 36a_CGG_W without added tRNA^Arg^(CCG), supporting competition between the endogenous and alternative tRNAs for CGG decoding. Analysis of these data (Figure S28) yielded a value of 0.37 ± 0.16 µM for the tRNA^Arg^(CCG) pre-existing in the CFE and a ratio of on-rates at the ribosomal A site for 36a_CGG_W to tRNA^Arg^(CCG) of 0.27 ± 0.12, perhaps due to lower levels of tryptophan charging.

To assess whether the fluorescence changes were accompanied by changes in reporter accumulation, we analyzed the reaction products by Coomassie-stained SDS– PAGE. A prominent band near the expected molecular mass of sfGFP (~27 kDa) was observed in all reporter-containing reactions, including sfGFP Y66R reactions that yielded only background CFP fluorescence (Figure 5B). Thus, low fluorescence did not necessarily indicate poor reporter production. Densitometric analysis showed broadly similar sfGFP band intensities across the Y66R reactions, ranging from 0.82 to 0.96 relative to the sfGFP Y66W control, which was normalized to 1.00 (Figure 5B). In the competition mixtures containing 0.4, 0.8 and 1.2 µM endogenous Arg tRNA, relative band intensities were 0.92, 0.90 and 0.82, respectively, compared with 0.95 for the reaction supplemented with 1.6 µM 36a_CGG_W alone. These modest differences in band intensity contrasted with the substantially larger reductions in CFP fluorescence. Together, these observations support the interpretation that competition between the endogenous and alternative tRNAs primarily alters the fraction of fluorescent, Trp-containing sfGFP, while overall reporter accumulation remains broadly similar.

Plotting CFP intensity against the concentration of supplemented 36a_CGG_W in the absence of added endogenous Arg tRNA revealed a nonlinear, concentration-dependent increase in reporter fluorescence (Figure 5C). The fitted curve closely described the measured response (R² = 0.975), with progressively smaller fluorescence gains at higher tRNA concentrations. This pattern is consistent with the approach toward saturation observed in the broader titration series (Figure S7A).

In the competition series, where the total added tRNA concentration remained fixed at 1.6 µM, CFP fluorescence increased as the mixture shifted from endogenous tRNA^Arg^(CCG) toward 36a_CGG_W (Figure 5D). The fitted curve captured this relationship (R² = 0.96), with the largest fluorescence increase occurring as 36a_CGG_W rose from 1.2 to 1.6 µM and added tRNA^Arg^(CCG) decreased from 0.4 µM to zero. Together with the lower signals in the competition mixtures than at matched concentrations of 36a_CGG_W alone, these results indicate that reporter fluorescence depends on both the concentration of the alternative tRNA and the abundance of its endogenous competitor. The fitted parameters show that the 36a_CGG_W tRNA is around 4 times less efficient than the endogenous tRNA^Arg^(CCG), which may indicate that the charging by the TrpRS is inefficient or incomplete.

### A single alternative tRNA supports simultaneous CGG reassignment in two reporter proteins

To determine whether a single alternative tRNA could mediate reassignment in two different proteins simultaneously, we co-expressed sfGFP Y66R and RFP R95R in the same *E. coli* CFPS reaction. Arg95 was selected because it is essential for RFP maturation and fluorescence [43]. Its codon was synonymously changed to CGG, preserving R95 under canonical decoding while allowing CGG-to-Trp reassignment to generate the nonfluorescent R95W variant. This loss-of-fluorescence readout complements the restoration of CFP fluorescence upon Trp incorporation at position 66 of sfGFP Y66R (Figure 6A). To verify whether reporter output scaled with template abundance during co-expression, we varied the CFP and RFP reporter DNA ratio while maintaining a total DNA concentration of 10 nM. Fluorescence from each reporter increased approximately in proportion to its respective template concentration (Figure 6B), supporting an approximately linear relationship between reporter DNA abundance and fluorescence output over the tested range.

**Figure 6.**
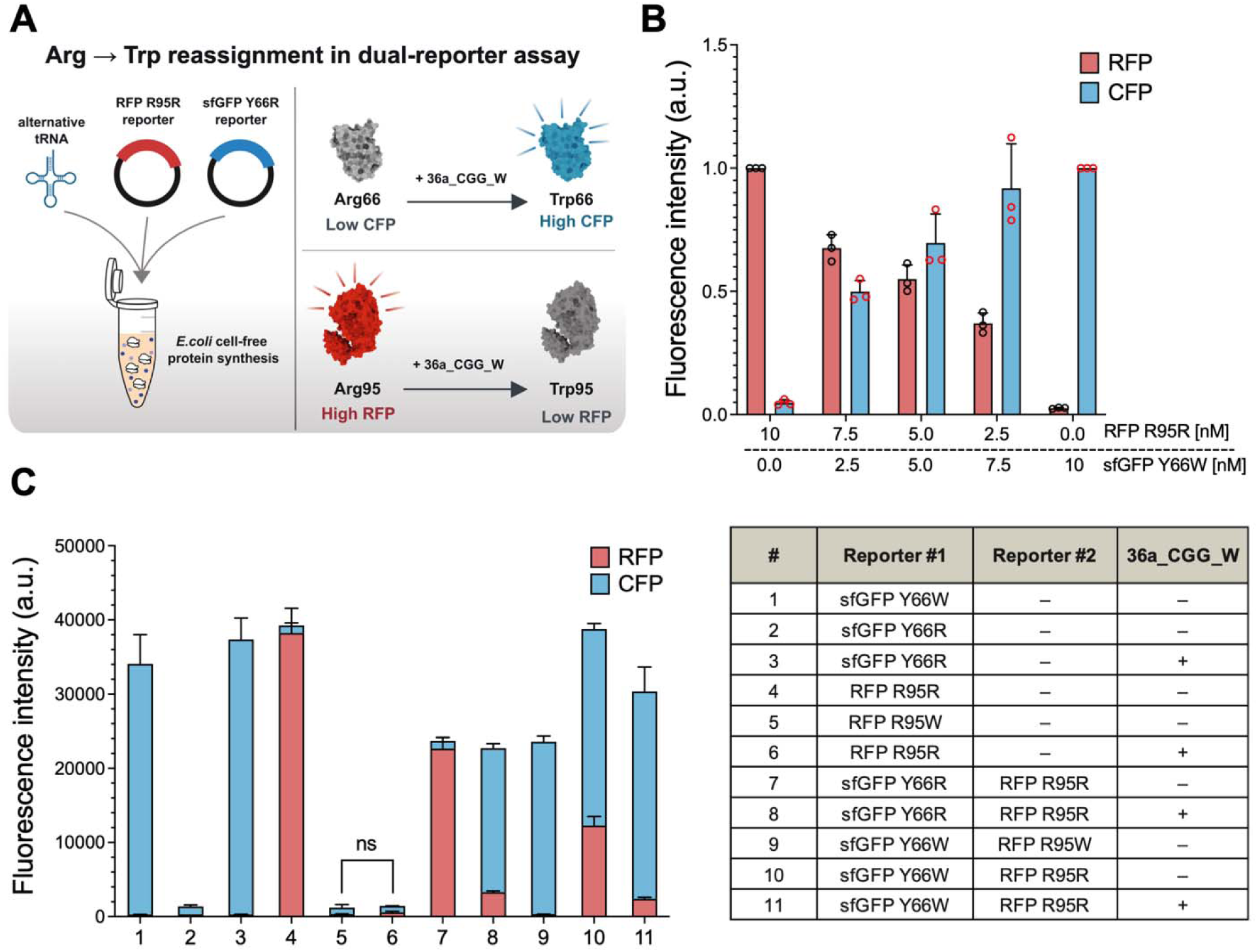
Dual-reporter assay for Arg-to-Trp reassignment. (**A**) Schematic of the assay. Supplementation with 36a_CGG_W tRNA promotes Trp incorporation at CGG codons, increasing CFP fluorescence from sfGFP Y66R and reducing RFP fluorescence through the Arg95-to-Trp substitution. (**B**) Normalized RFP and CFP fluorescence at varying reporter DNA ratios, with total reporter DNA maintained at 10 nM. (**C**) CFP and RFP fluorescence from individual or combined reporters in the presence (+) or absence (−) of 36a_CGG_W, as indicated in the table. sfGFP Y66W and RFP R95W serve as positive and negative controls, respectively.

When expressed individually, sfGFP Y66R produced background CFP fluorescence without supplemented tRNA, whereas addition of 36a_CGG_W restored fluorescence comparable to the sfGFP Y66W positive control (Figure 6C, conditions 1– 3). Conversely, supplementation with the same tRNA reduced RFP R95R fluorescence to near background, resembling the nonfluorescent RFP R95W control (conditions 4–6). When both reporters were co-expressed, adding 36a_CGG_W increased CFP fluorescence and substantially reduced RFP fluorescence within the same reaction (conditions 7–8). Reactions containing sfGFP Y66W and RFP R95R likewise showed reduced RFP fluorescence while retaining substantial CFP fluorescence after tRNA supplementation (conditions 10–11). These complementary responses support the ability of a single alternative tRNA to mediate CGG-to-Trp reassignment in two distinct protein targets simultaneously in *E. coli* CFPS.

### *E. coli* aminoacyl-tRNA synthetases are sufficient for decoding

Besides the tRNA itself, a second potential determinant of an alternative genetic code is the aminoacyl-tRNA synthetase (AARS) that recognizes and charges the specialized tRNA. Our results suggest that a specialized AARS is not strictly required for decoding by the candidate tRNAs, because the endogenous *E. coli* AARSs were sufficient to support their activity in CFPS. Phylogenetic signatures of AARS specialization could nonetheless exist, for example if the AARSs of the ingroup (species using the new code) formed a single clade on an extended branch relative to the AARSs of closely related outgroup species.

To examine this possibility, we generated ingroup–outgroup phylogenies for the relevant AARS groups (Figures S13–S18). For GlyRS in codes 25 and 37, MetRS in code 34, GlnRS in code 35 and TrpRS in code 37 (Figures S13, S14, S17 and S18, respectively), the ingroup formed a single clade but not on a particularly long branch. For both lineages using code 36 (Figures S15 and S16), TrpRS occurs in two distinct forms, W1 and W2, whose distribution suggests horizontal transfer: the ingroup includes genomes with the W1 form and genomes with the W2 form (no genome encodes both), and the same mixing occurs in the outgroup. These patterns make it unlikely that TrpRS is a primary determinant of CGG-to-Trp reassignment in code 36 and focus attention on the tRNAs, which are more clearly distinguished between ingroup and outgroup.

Consistent with charging by the host TrpRS, all four tRNAs predicted to decode Arg codons as Trp (36a_CGG_W, 36o_CGG_W, 37_CGG_W and 37_CGA_W (though 37_CGA_W was inactive)) and the 4_UGA_W carry guanine at the discriminator position (position 73), a major identity determinant of *E. coli* tRNA^Trp^, whereas the Gly-decoding UGA tRNAs (25_UGA_G, 37_UGA_G, 37w_UGA_G) carry uracil at this position (Table S1). Aminoacylation assays with purified *E. coli* TrpRS would test this directly [44, 45].

### A native code 36 bacterium decodes CGG codons as Trp

The experiments above show that natural code 36 and code 37 tRNAs can decode CGG as Trp in a heterologous system. We therefore asked whether this occurs in a native bacterium. Most of the genomes with predicted sense-to-sense codon changes are metagenomic, from uncultured organisms. However, we were able to obtain a genome-sequenced isolate *Anaerococcus prevotii* ATCC 9321 assigned to code 36 (Figure 1). The chromosome of this strain encodes 1, 702 annotated proteins, among which CGG is the rarest codon: 39 of 555, 757 sense codons, in 37 proteins (Figure S22). Its only CCG-anticodon tRNA is 78% and 84% identical to the code 36 tRNAs 36a and 36o that decoded CGG as Trp in CFPS (Figure S23; Table S7).

We analyzed an *A. prevotii* culture by data-independent acquisition (DIA) proteomics and searched the data against its annotated (CGG-to-Arg) proteins together with variants in which every CGG is translated as Trp (see Methods). Trp, unlike Arg, is not cleaved by trypsin, so a CGG-to-Trp codon merges the short Arg-form tryptic pieces on either side of the site into one longer peptide that should improve detection (Figure 7A; Figure S20). At 1% precursor FDR, 16, 277 precursors (13, 800 peptides ranging in length from 7 to 30 residues) were detected. These peptides provided evidence of expression for 1, 347 (79.1%) of the full list of 1, 702 proteins. The CGG-codon proteins had a similar expression rate, 27 (73.0%) of the 37 proteins (Figure 7B; Figure S19). The genes for these 27 expressed proteins contained 29 CGG codons. Six of the CGG sites were covered by an identified Trp-containing peptide (Figure 7C; Table S5): A<u>W</u>EGSLR, YG<u>W</u>EGLTFPGGK, EI<u>W</u>ENNTPK, IS<u>W</u>EISK, VIPM<u>W</u>DGNSK and VIEG<u>W</u>AR. Detection of only six (20.7%) of the 29 expressed CGG codons is roughly compatible with rates of coverage of protein residues by the identified peptides. The residue coverage rate for all expressed proteins was 31.4%, and for the 27 expressed CGG-codon proteins it was 20.8% (4, 071 detected of 19, 608 total residues). None of the six CGG-to-Trp confirming peptides occurs in the canonical proteome. No Arg-form peptide was detected at any CGG site, whereas about 6.6 of the 22 Arg-form peptides that were detectable in principle would be expected if CGG encoded Arg (P = 1.5 × 10^−4^); the six Trp-form peptides found are close to the 8.3 expected if CGG encodes Trp (Figure S21).

**Figure 7.**
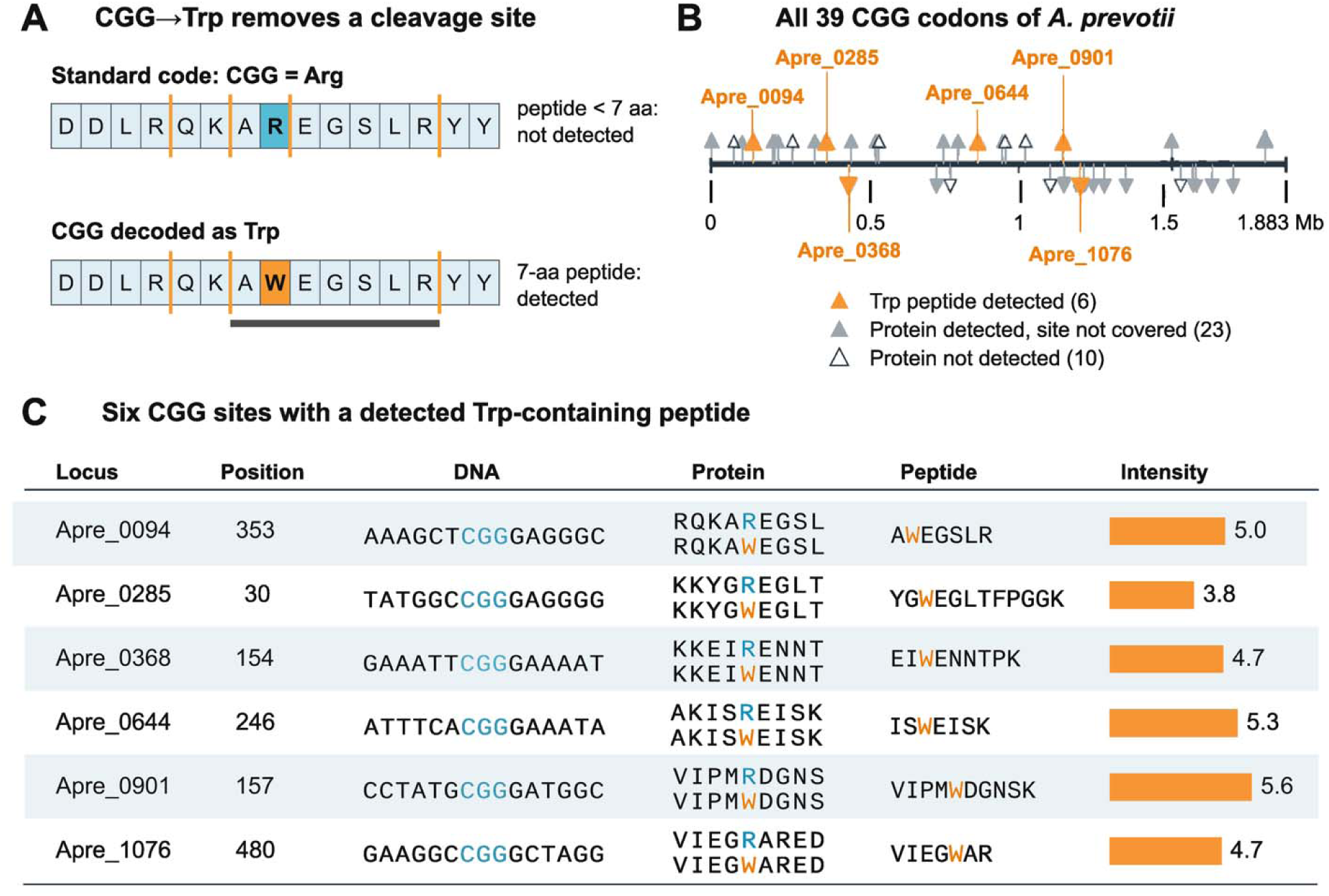
Native-host proteomics detects Trp at CGG codons in *A. prevotii*. (**A**) Decoding CGG as Trp instead of Arg removes a trypsin cleavage site. At position 353 of Apre_0094, cleavage of the Arg-containing sequence produces QK, AR and EGSLR, each shorter than the 7-residue detection threshold. The Trp-containing sequence produces the detectable 7-residue peptide AWEGSLR. Orange lines indicate trypsin cleavage sites. (**B**) Locations of all 39 in-frame CGG codons on chromosome CP001708.1. Triangles above and below the axis indicate the forward and reverse strands, respectively. Orange indicates a detected Trp-containing peptide; gray indicates a detected protein without peptide coverage of the CGG site; open triangles indicate proteins not detected. (**C**) The six CGG sites with detected Trp-containing peptides. Each row shows the locus, amino acid position, surrounding coding-strand DNA, reference Arg-containing and predicted Trp-containing protein sequences, detected peptide, and peptide precursor signal intensity (log10, arbitrary units). Data are from one culture analyzed in a DIA run. Details are provided in Methods and Figures S19–S27.

Misassignment is unlikely to explain all six peptides: the expected number of false-positive Trp-form peptide identifications at 1% FDR is about 0.2 (Figure S21), and the raw DIA scans support the identifications, most clearly for ISWEISK and VIPMWDGNSK, whose Trp-containing fragment ions are among the most intense and co-elute with the precursor signal, and less clearly for the other four peptides (Figures S26 and S27; Table S8). Taken together, these results are consistent with translation of CGG as Trp in a native code 36 bacterium; however, several limits apply: 1) The data come from one culture. 2) At five of the six detected sites the Arg-form pieces are too short to detect, so a minority of Arg cannot be excluded there; the absence of Arg-form peptides is informative only at the 19 expressed CGG sites that have detectable partner peptides. 3) The identifications rest on DIA-NN scoring. 4) The cultured strain is the type strain whose genome was sequenced (CP001708.1), but we did not sequence our own stock, so CGG-to-TGG variants in the stock cannot be excluded.

## DISCUSSION

We provide the first experimental test of the sense-to-sense genetic codes predicted for bacteria by Codetta [26]. In an *E. coli* cell-free translation system, natural tRNAs from the *Anaerococcus* and *Onthovivens* lineages (code 36) and from the *Absconditabacterales* (code 37) decoded CGG as Trp, supporting code 36 in both lineages and the Arg CGG-to-Trp change of code 37 (Figure 3B). The 37w_UGA_G tRNA also decoded UGA as Gly, supporting a second change proposed for code 37, whereas the Arg CGA-to-Trp assignment of code 37 was not supported in our assay because the candidate 37_CGA_W tRNA did not restore reporter fluorescence (Figure 3B). Proteomics of the native code 36 bacterium *A. prevotii* detected Trp-containing peptides at six CGG codons and no Arg-form peptide at any CGG site, consistent with CGG decoding as Trp in its natural host (Figure 7). Sense codon reassignment has been engineered previously in *E. coli*, but typically to non-canonical amino acids and with engineered orthogonal tRNA/synthetase pairs; here, natural tRNAs from distant bacteria executed predicted reassignments of canonical amino acids.

We suggest that code 36 and the experimentally supported assignments of code 37 (Arg CGG-to-Trp and Stop UGA-to-Gly) be added to official lists of NCBI genetic code variants, under the names “*Anaerococcus* and *Onthovivens* code” and “*Absconditabacterales* code”, respectively. We also note that the name applied to code 25 (“Candidate Division SR1 and *Gracilibacteria* code”) is misleading: the genome-based taxonomy GTDB places code 25 (order BD1-5) and code 37 (order *Absconditabacterales*) as sister clades within the class *Gracilibacteria*. To avoid confusion, we propose renaming code 25 the “BD1-5 code”.

Genomes with CGG-to-Trp reassignment clustered within the low-GC content range, suggesting an association between genomic nucleotide composition and alternative CGG decoding (Figure S28). The relatively low abundance of CGG in these genomes may facilitate reassignment by limiting the number of coding sites affected, although this association alone does not establish a causal role for low GC content.

Several candidate tRNAs produced reporter signals comparable to or greater than the corresponding Trp-encoded positive controls. For example, 36o_CGG_W and 37_CGG_W gave mean CFP signals of 128.7% and 127.1% of the Y66W control, respectively. Reporter intensity is not a direct measure of the fraction of codons decoded as Trp, because it also depends on protein yield, folding and chromophore maturation; indeed, the Y66R reporters accumulated somewhat more total protein than the Y66W control (Figure 3D). Nevertheless, the competition experiments show that output changes predictably with the ratio of alternative to canonical CGG-decoding tRNAs (Figure 5). The ability of phylogenetically distant tRNAs to be aminoacylated and used by the *E. coli* translation system suggests that naturally occurring tRNAs may provide useful starting points for engineered translation systems [16, 27, 46].

Reassignment at multiple engineered sites across different proteins supports authentic decoding rather than a reporter-specific artifact. The sfGFP Y66R reporter provided a sensitive readout because Trp incorporation at the chromophore-forming position restores cyan fluorescence. Reassignment activity was also detected in firefly luciferase and NanoLuc reporters (Figure 4), as well as in sfGFP and NanoLuc constructs containing multiple CGG substitution sites (Figures S3 and S12).

Furthermore, the dual-reporter assay supported simultaneous CGG-to-Trp reassignment in two distinct protein targets by a single alternative tRNA within the same CFPS reaction (Figure 6). Together, these findings demonstrate activity across several reporter contexts, an important consideration for genetic code engineering, where reassignment must accommodate diverse coding sequences, protein folds and local sequence contexts.

We observed variable performance of individual tRNAs across reporters and target sites. This indicates that the apparent reassignment output depends on several interacting factors. In the *E. coli* CFPS system, these include competition with endogenous tRNAs, aminoacylation efficiency, nearby codons, local mRNA structure, ribosome kinetics and the structural importance of the substituted residue [41, 42, 47]; changes in any of these variables can influence the apparent output of each reporter. A protein with several reassigned codons also gives canonical Arg tRNAs repeated opportunities to introduce a disruptive substitution. Peptide-level tandem mass spectrometry of the reporters, which would measure the fraction of Trp and Arg incorporated at each CGG site independently of reporter activity, is therefore the logical next step.

Several limitations should be noted. First, the tRNAs were tested with *E. coli* extracts and translation machinery rather than with extracts of their native hosts (which are often unculturable). The tRNA recognition rules of *E. coli* AARSs may bias the results if they differ from those of the native hosts, and the failure of a tRNA in our assay (code 25, code 37 UGA, 37_CGA_W) is therefore not evidence against the predicted code; for 37_CGA_W, competition from the native tRNA^Arg^(ICG), which decodes CGA, may also contribute [45, 50]. Second, the extract contains endogenous tRNAs and release factors that compete with the supplemented tRNAs, so the fractional output reflects competition as well as intrinsic decoding capacity; reconstituted translation systems [51, 52] or oligonucleotide-mediated depletion of endogenous tRNAs [49] could remove this competition. Third, the fluorescence and luminescence assays used different extract preparations from those used for the ¹ C-leucine incorporation assays, which were performed in a collaborator’s laboratory. Although this limits direct quantitative comparisons between assays, the complementary findings from the two laboratories support our overall conclusions.

By showing that Codetta-indicated tRNAs from distant bacteria operate in an *E. coli* CFPS system and mediate Arg-to-Trp reassignment in several reporter contexts, and that a native code 36 bacterium carries Trp at CGG codons, this study establishes a route for testing genetic code hypotheses outside and inside their native biological settings. As genomic surveys continue to uncover unusual coding systems, and new genomically recoded organisms are built and need tRNA charging systems [48, 53–56], combining reporter-based CFPS screens with mass spectrometry confirmation should provide an efficient way to screen and confirm natural tRNAs that expand the known genetic codes and create new opportunities for genetic code engineering.

## SUPPLEMENTARY DATA

Supplementary data are available at NAR online.

## ACKNOWLEDGEMENTS

The authors thank Michael Daly (Waters Corporation) for assistance with mass spectrometry and data analysis, and Tong-Un Chae (Stanford University) for assistance with cell-free extract optimization and helpful discussions regarding data interpretation. Novogene performed sample processing and DIA-NN proteomics of *A. prevotii*.

## Author contributions

Paul H. Kim (Conceptualization [equal], Formal analysis [equal], Investigation [lead], Methodology [equal], Validation [equal], Visualization [lead], Writing—original draft [lead], Writing—review & editing [lead]), Megan A. McSweeney (Investigation [supporting], Validation [supporting], Writing—review & editing [supporting]), Joshua T. Hong (Investigation [supporting], Validation [supporting]), Joanna P. Stecynska (Investigation [supporting]), Hannah Chan (Investigation [supporting]), Michael C. Jewett (Methodology [equal], Resources [equal], Writing— review & editing [equal]), Yooli K. Light (Conceptualization [equal], Supervision [lead], Methodology [equal], Writing—review & editing [equal]), Kelly P. Williams (Conceptualization [equal], Supervision [equal], Writing—review & editing [equal]), and Joseph S. Schoeniger (Conceptualization [equal], Supervision [equal], Writing—review & editing [equal]).

## FUNDING

This work was supported by the Laboratory Directed Research and Development program (project 236996) at Sandia National Laboratories, a multimission laboratory managed and operated by National Technology and Engineering Solutions of Sandia LLC, a wholly owned subsidiary of Honeywell International Inc. for the U.S. Department of Energy’s National Nuclear Security Administration under contract DE-NA0003525.

## COMPETING INTERESTS

M.C.J. has a financial interest in Pearl Bio, Inc., Ridge Bio, Synolo Therapeutics, Inc., and Gauntlet Bio. M.C.J.’s interests are reviewed and managed by Northwestern University and Stanford University in accordance with their competing interest policies. All other authors declare no competing interests.

## DATA AVAILABILITY

The data underlying this article are available in the article and in its online supplementary material. The mass spectrometry proteomics data (raw file, DIA-NN output, search database) have been deposited in the ProteomeXchange Consortium via the PRIDE partner repository with the dataset identifier PXD086557 and project DOI: 10.6019/PXD086557. The per-site analysis tables are provided as Supplementary Data. Reporter plasmid sequences are available in the Supplementary Data.

---

**Supplementary Figure S1.**
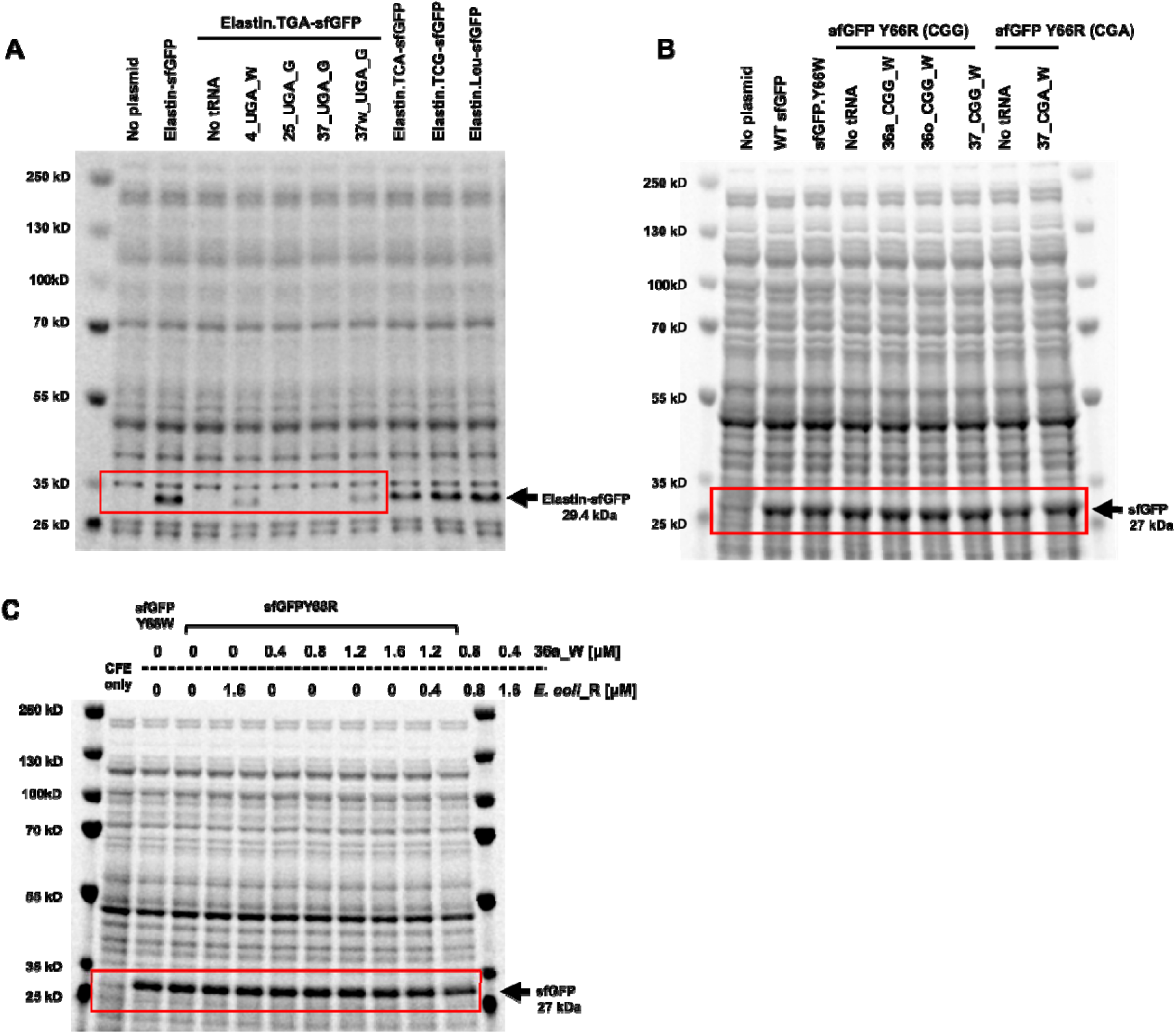
Full Coomassie-stained SDS–PAGE gels of CFPS reporter production. Gels show proteins produced in CFPS reactions with or without alternative tRNAs. Red boxes indicate the regions presented as cropped gel images in the corresponding main figure panels. Arrows mark the expected positions of the full-length reporter proteins. (**A**) CFPS reactions expressing an Elastin-sfGFP reporter containing an in-frame UGA stop codon were supplemented with alternative tRNAs predicted to recognize UGA and promote amino acid incorporation. Wild-type Elastin-sfGFP variants containing Tyr, Trp, or Leu substitutions at the target position were included as positive controls. (**B**) CFPS reactions expressing sfGFP reporters containing a codon substitution at position Y66. Alternative tRNAs predicted to decode CGG or CGA as Trp were added as indicated. (**C**) CFPS reactions expressing sfGFP Y66R with the indicated concentrations of 36a_CGG_W and endogenous *E. coli* tRNA^Arg^(CCG). sfGFP Y66W and extract-only (CFE only) reactions were included as controls.

**Supplementary Figure S2.**
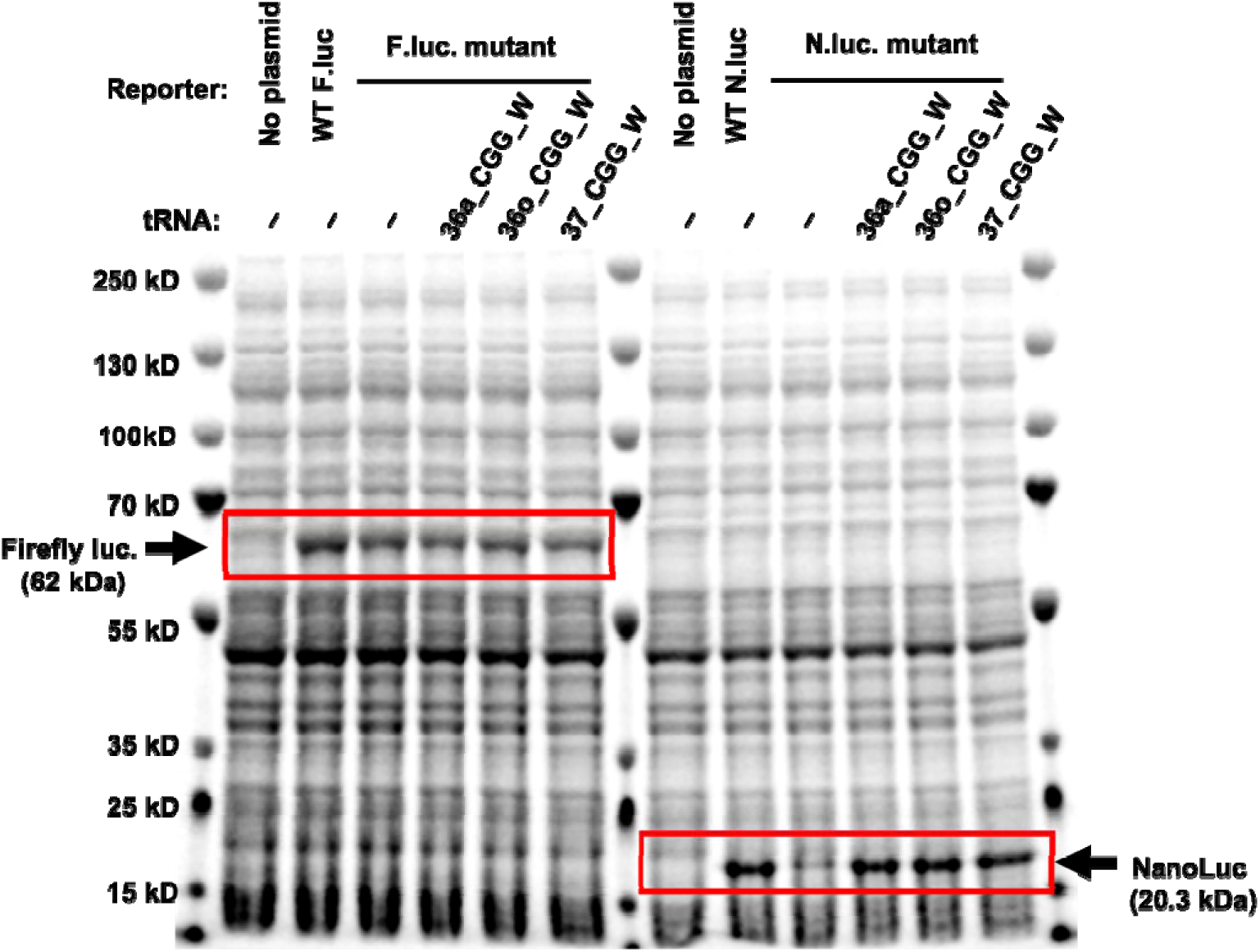
SDS-PAGE analysis of firefly luciferase and NanoLuc reporter production in CFPS reactions. CFPS reactions expressing wild-type or mutant luciferase reporters were analyzed by SDS-PAGE to assess reporter protein production under the same conditions used for luminescence assays. Firefly luciferase samples included no-plasmid control, wild-type firefly luciferase, and mutant firefly luciferase reactions lacking or supplemented with the indicated CGG-decoding alternative tRNAs. NanoLuc samples included no-plasmid control, wild-type NanoLuc, and mutant NanoLuc reactions lacking or supplemented with the same alternative tRNAs. Red boxes highlight bands at the expected molecular weights for firefly luciferase and NanoLuc. Arrows indicate firefly luciferase at 62 kDa and NanoLuc at 20.3 kDa.

**Supplementary Figure S3.**
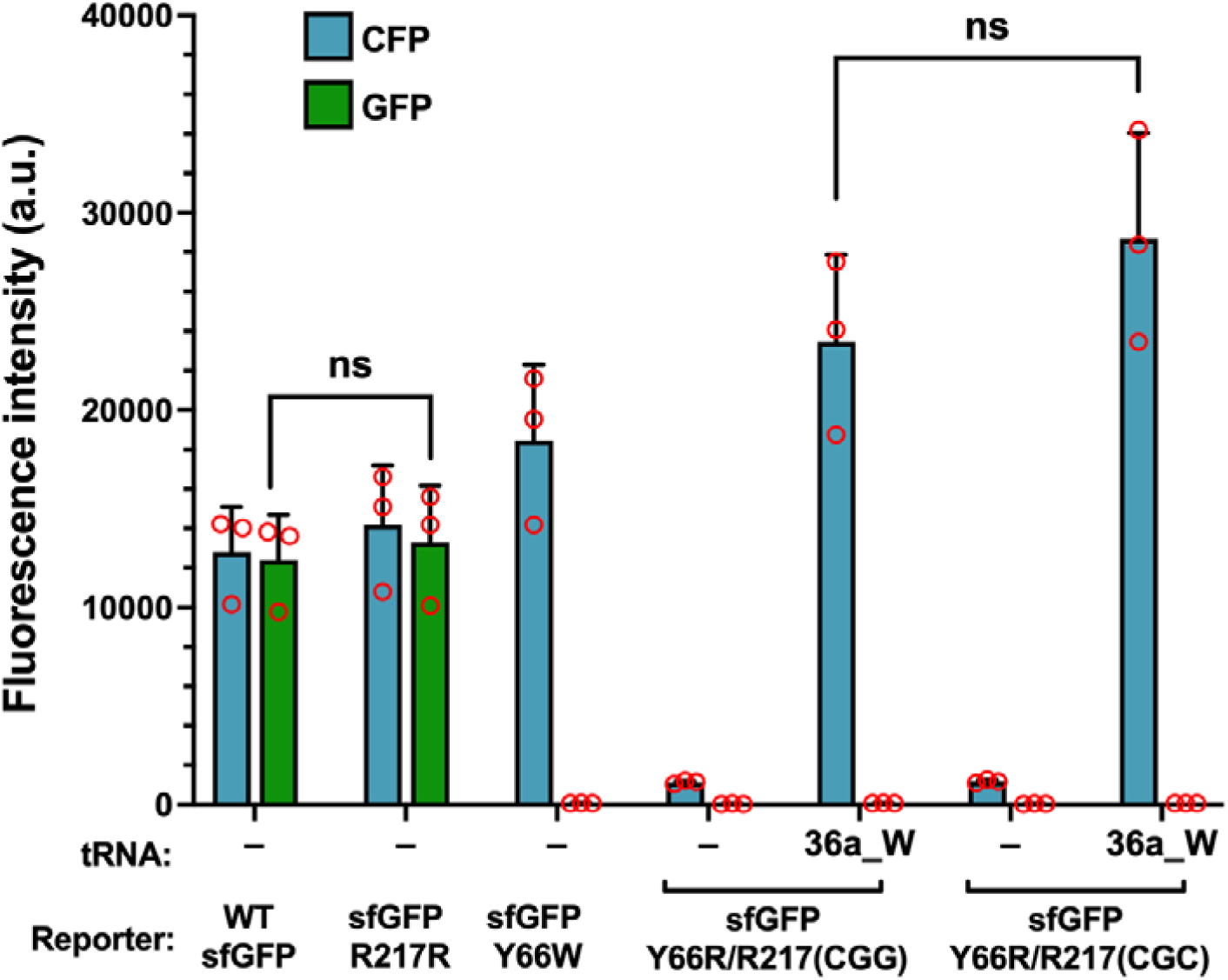
Comparison of single-site, sfGFP Y66R/R217(CGC) and two-site, sfGFP Y66R/R217(CGG), CGG reporters for Arg-to-Trp reassignment in sfGFP. CFP fluorescence measurements from CFPS reactions expressing reporter constructs containing either one or two target codons (CGG) for alternative tRNA-mediated amino acid substitution. Reactions were supplemented with the code 36a_CGG_W alternative tRNA, and fluorescence intensity was measured following 20-hour incubation. The two-site reporter produced slightly lower CFP fluorescence than the single-site reporter, but the difference was not statistically significant (ns). Blue bars show the CFP channel and green bars the GFP channel; WT sfGFP and sfGFP R217(CGC) are shown as controls. Arbitrary units differ from those in main Figure 3 because plate-reader settings were optimized for each experiment (Materials and methods). Reactions lacking alternative tRNAs showed background-level fluorescence. Error bars represent standard deviation from at least three independent CFPS reactions. An unpaired two-tailed Student’s t-test was used (*P* values, * *P* < 0.05, \*\**P* < 0.01 indicate significant differences between samples; ‘ns’ represents not significant).

**Supplementary Figure S4.**
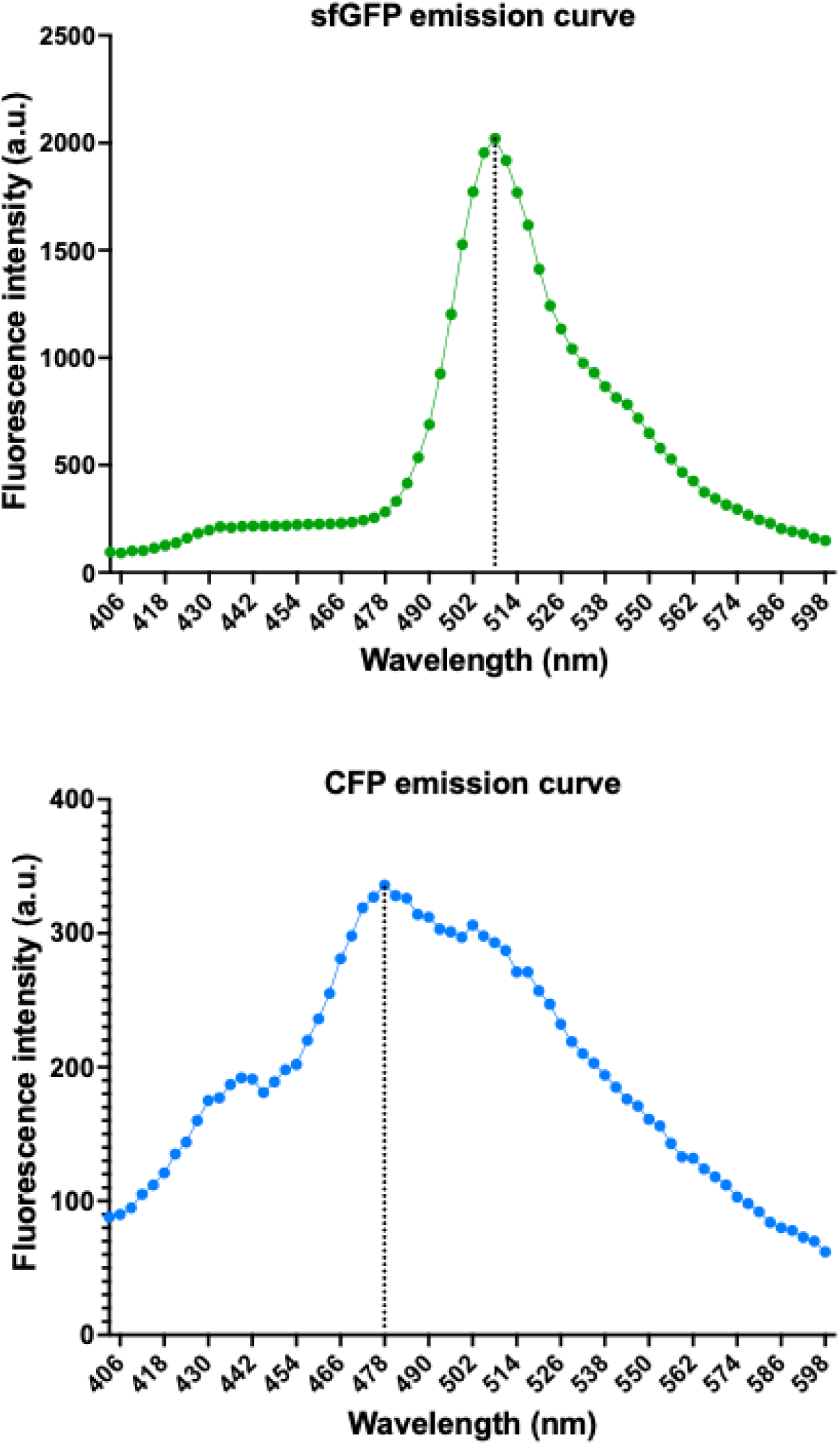
Emission spectra of sfGFP and CFP expressed in the CFPS system. Fluorescence emission spectra of sfGFP and CFP produced in CFPS reactions were measured to determine the optimal detection wavelengths for fluorescence quantification. Emission scans were performed using a Tecan plate reader across the indicated wavelength range. sfGFP exhibited a maximal fluorescence emission peak at 508 nm, while CFP showed maximal emission at 478 nm. Dotted lines mark the emission maxima.

**Supplementary Figure S5.**
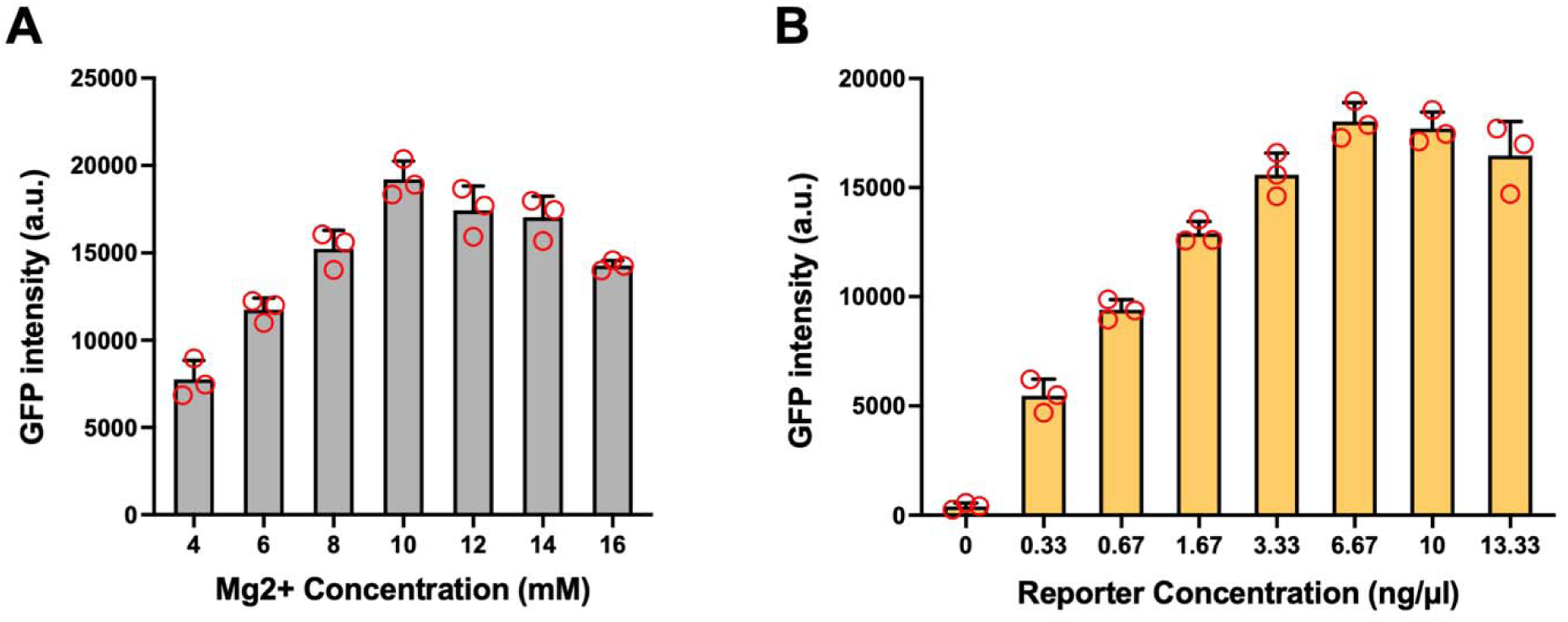
Optimization of CFPS reaction conditions for sfGFP reporter expression. GFP fluorescence was measured to optimize key reaction parameters in the *E. coli* cell-free protein synthesis system. (**A**) Effect of Mg^2+^ concentration on sfGFP reporter expression. GFP intensity increased with Mg^2+^ concentration and reached maximal signal at approximately 10 mM, with reduced output at higher concentrations. (**B**) Effect of reporter plasmid concentration on GFP expression. GFP intensity increased in a dose-dependent manner and plateaued at 6.67 ng/µL, with no further increase at the highest concentration tested. Bars represent mean ± SD; red circles indicate individual replicate reactions.

**Supplementary Figure S6.**
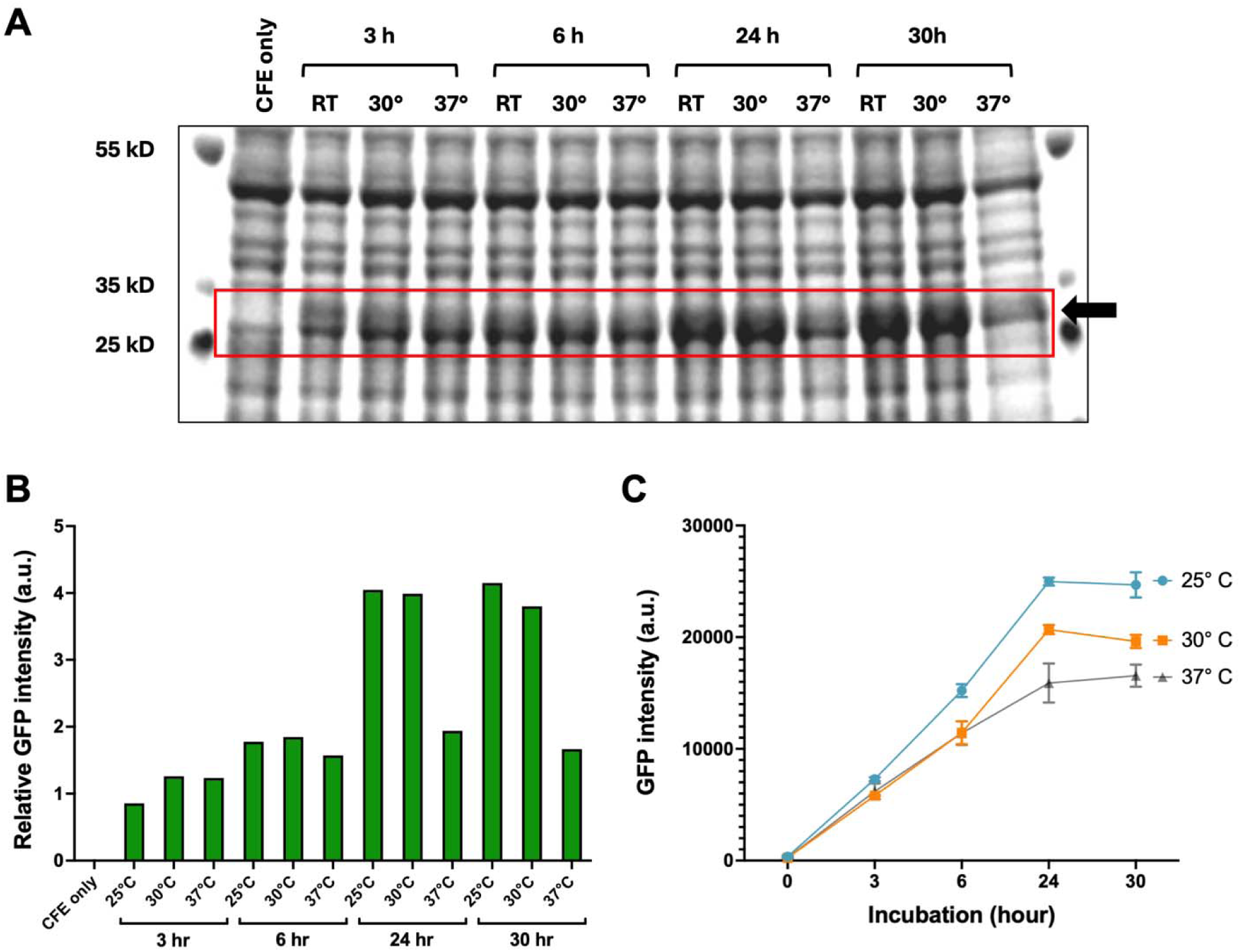
Optimization of incubation temperature and reaction time for sfGFP expression in cell-free protein synthesis reactions. (A) SDS–PAGE analysis of sfGFP expressed in CFPS reactions incubated at the indicated temperatures and time points. Samples were collected at each condition and analyzed to assess sfGFP production level. (B) Quantification of sfGFP band intensity from the SDS–PAGE gels shown in panel A. Band intensities were normalized to total lane protein and compared across incubation conditions to assess relative reporter protein yield. (C) GFP fluorescence measurements from CFPS reactions monitored at the indicated time points using a plate reader. Fluorescence intensity was used as an independent measure of sfGFP production and maturation under each reaction condition. Error bars represent standard deviation from three replicates.

**Supplementary Figure S7.**
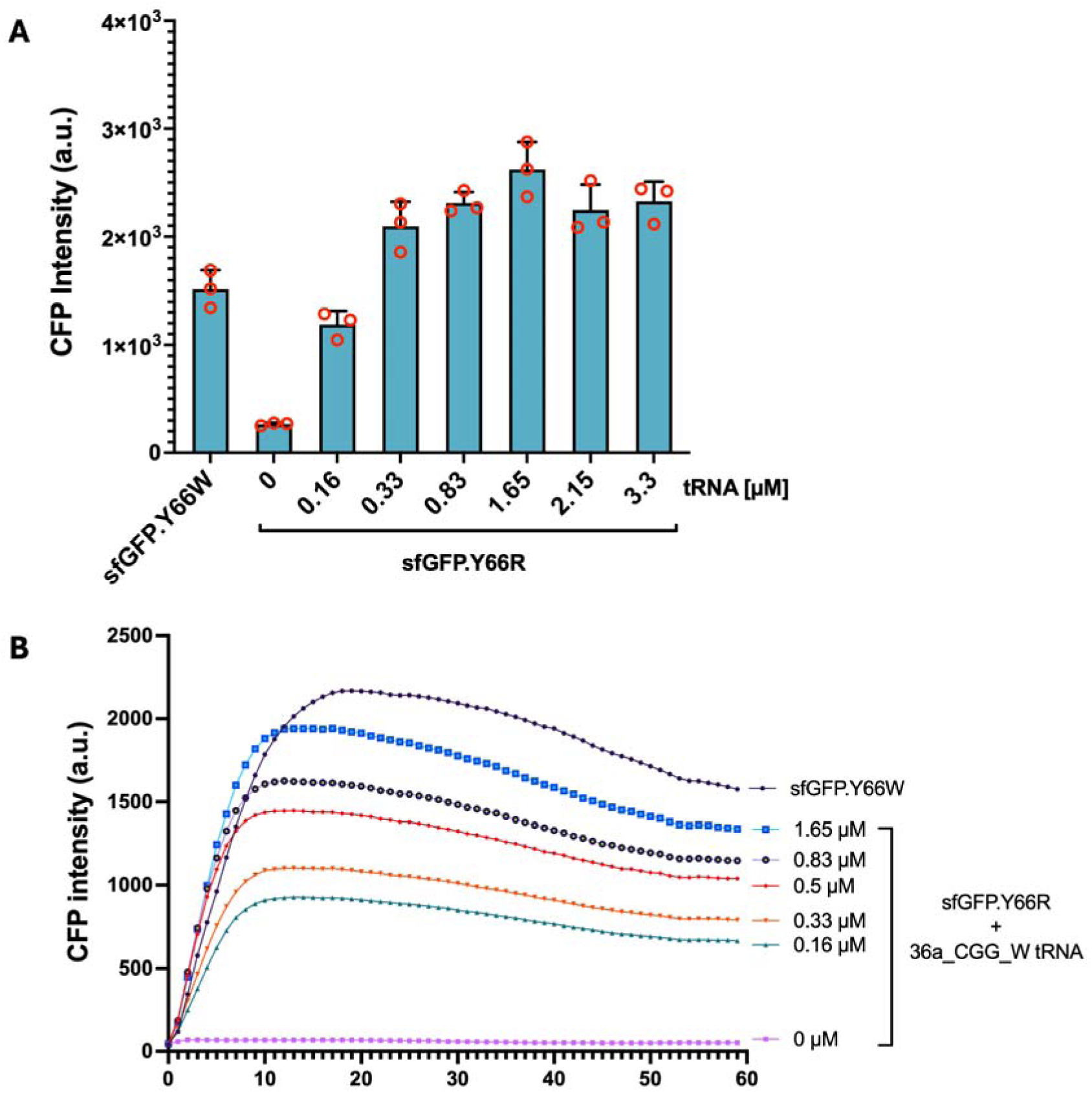
Optimization of alternative tRNA concentration in CFPS. (**A**) CFPS reactions expressing CFP were supplemented with increasing concentrations of 36a_CGG_W tRNA to determine the optimal amount required for efficient amino acid substitution and protein expression. CFP fluorescence intensity was measured following 20-hour incubation at 25 °C. The addition of alternative tRNA enhanced CFP production in a concentration-dependent manner, with 1.65 µM yielding the highest CFP signal. Red circles represent individual replicates, and error bars represent standard deviation. (**B**) Kinetic analysis of CFP was performed to evaluate the activity of the alternative tRNA over time in CFPS reactions. CFP intensity was measured every hour for 60 h with various concentrations of code 36a_CGG_W tRNA. CFP signal increased progressively over time and reached maximal intensity around 15 h of incubation before gradually decreasing. The sfGFP Y66W construct served as a positive CFP control, while reactions lacking alternative tRNA, 0 µM, showed minimal CFP signal.

**Supplementary Figure S8.**
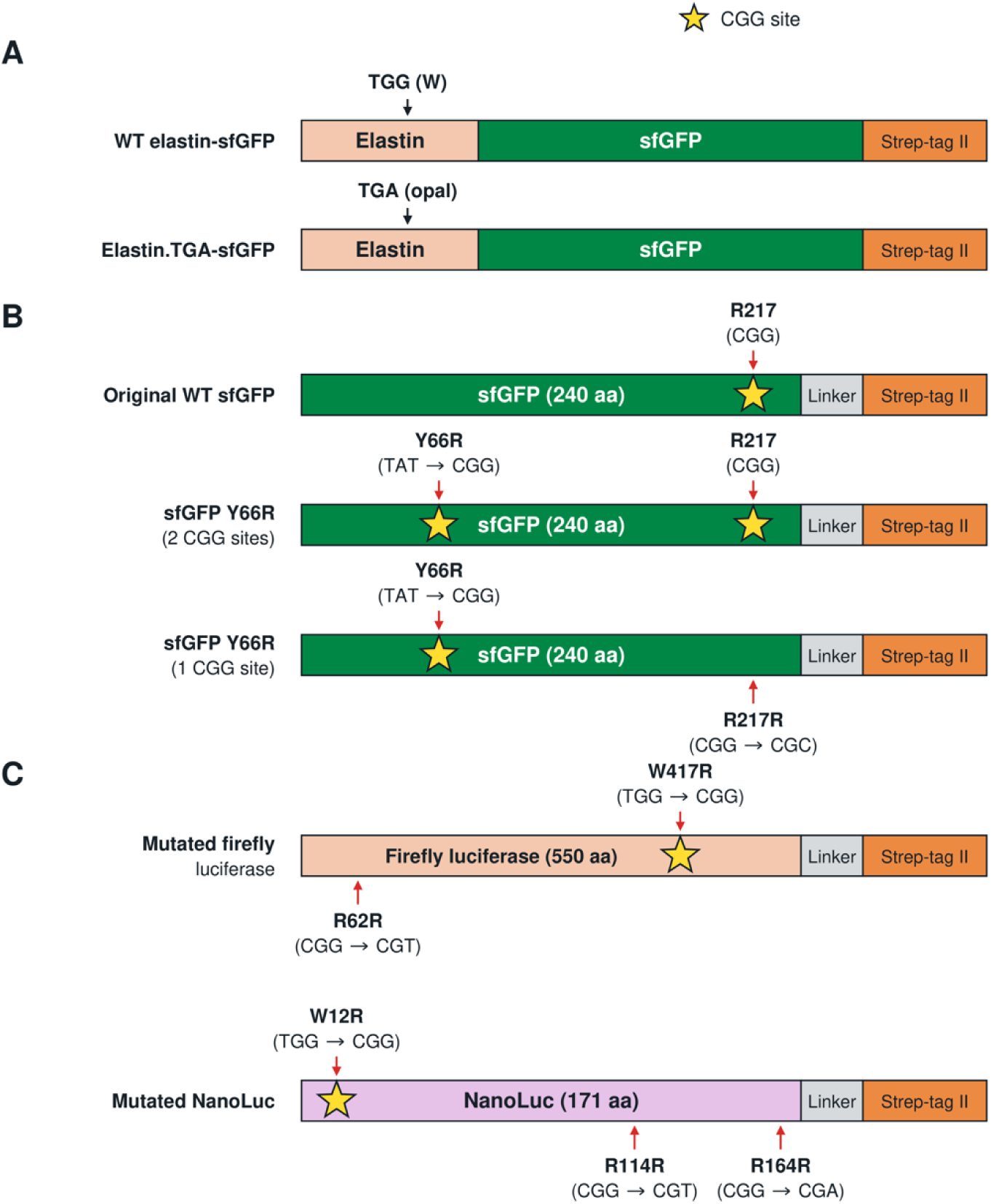
Schematic design of reporter constructs used to evaluate codon reassignment. (**A**) Elastin–sfGFP fusion reporters containing either the native tryptophan codon (TGG) or an opal stop codon (TGA) in the middle of elastin coding region. Both constructs contain a C-terminal Strep-tag for affinity purification. (**B**) sfGFP reporter variants used to evaluate CGG-mediated amino acid substitution. In the sfGFP Y66R construct, Y66 was changed from TAT to CGG, whereas the R217 codon was synonymously changed from CGG to CGC, leaving Y66 as the only CGG reassignment site. The sfGFP-Y66R (CGA) reporter used with the 37_CGA_W tRNA carries CGA instead of CGG at position 66 (not drawn). (**C**) Firefly luciferase and NanoLuc reporters are designed to contain a single CGG codon at a functionally essential tryptophan position. In firefly luciferase, W417 was changed from TGG to CGG, and the endogenous R62 codon was synonymously changed from CGG to CGT. In NanoLuc, W12 was changed from TGG to CGG, while endogenous CGG codons at R114 and R164 were synonymously changed to CGT and CGA, respectively.

**Supplementary Figure S9.**
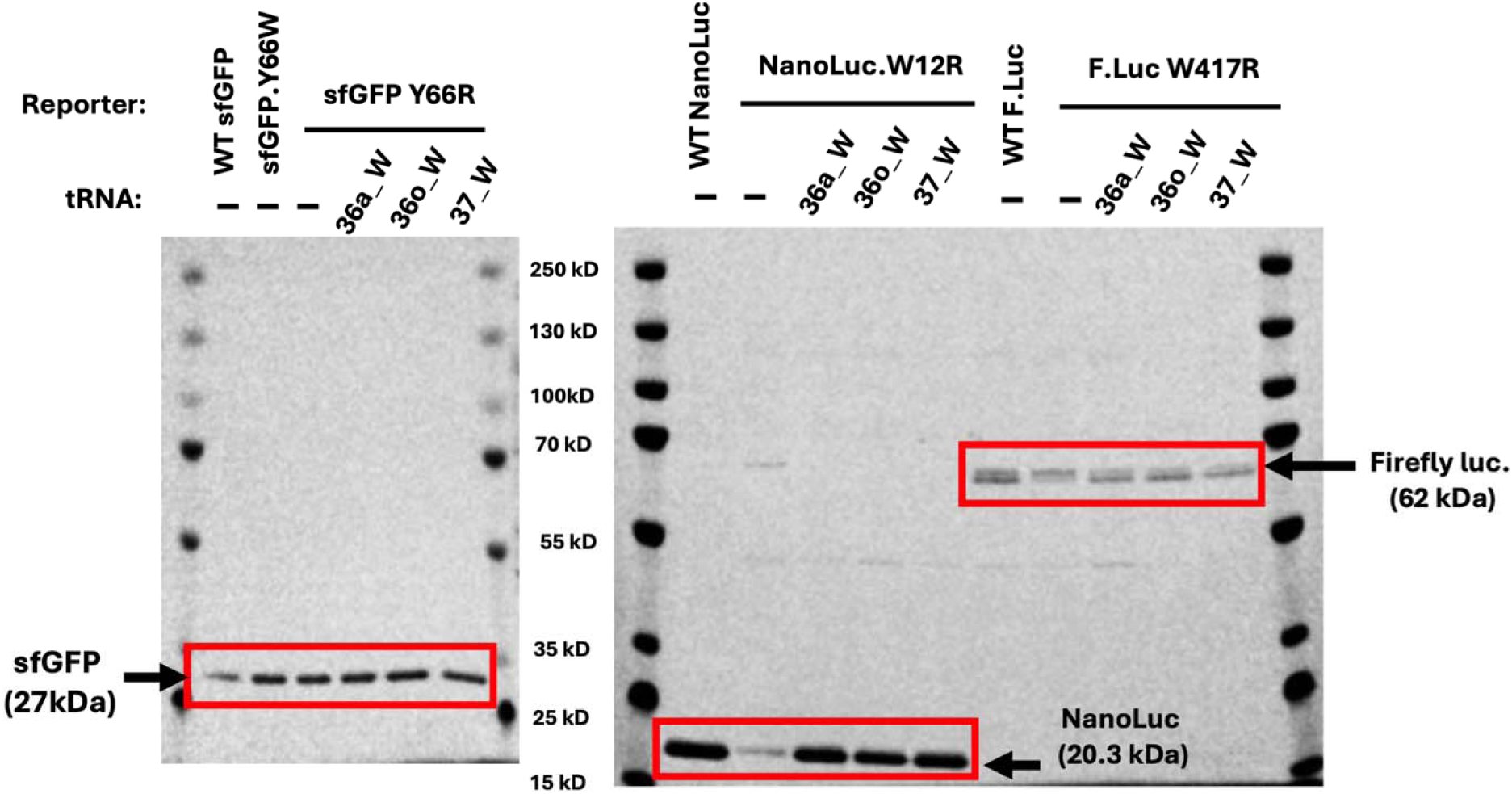
Preparation and purification of CFPS-generated proteins for mass spectrometry analysis. Proteins synthesized by cell-free protein synthesis reactions were purified using Strep-Tactin affinity purification methods prior to mass spectrometry analysis. SDS-PAGE analysis demonstrates prominent bands observed at the expected molecular weight marked by red boxes. These purified samples were subsequently used for downstream mass spectrometry to verify Arg→Trp amino acid reassignments.

**Supplementary Figure S10.**
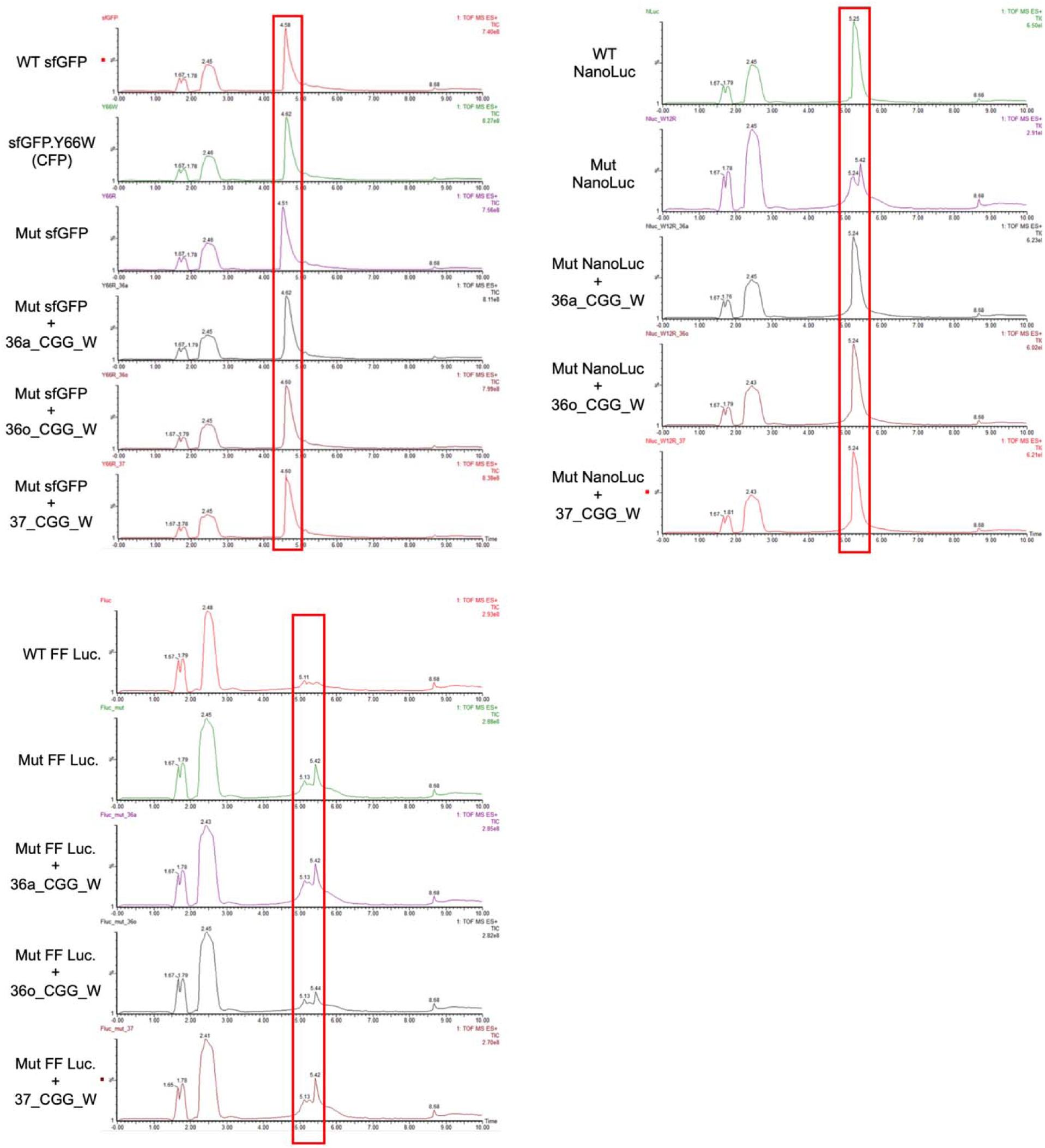
LC-MS retention profiles of purified reporter proteins used for intact-mass analysis. Total ion chromatograms acquired by positive-mode electrospray ionization time-of-flight mass spectrometry (ESI-TOF-MS) are shown for sfGFP, NanoLuc, and firefly luciferase (FF Luc.) reporter variants. Red boxes indicate the retention-time regions containing the reporter protein signals used for subsequent intact-mass spectral extraction and deconvolution. Chromatograms are displayed as relative ion intensity versus retention time.

**Supplementary Figure S11.**
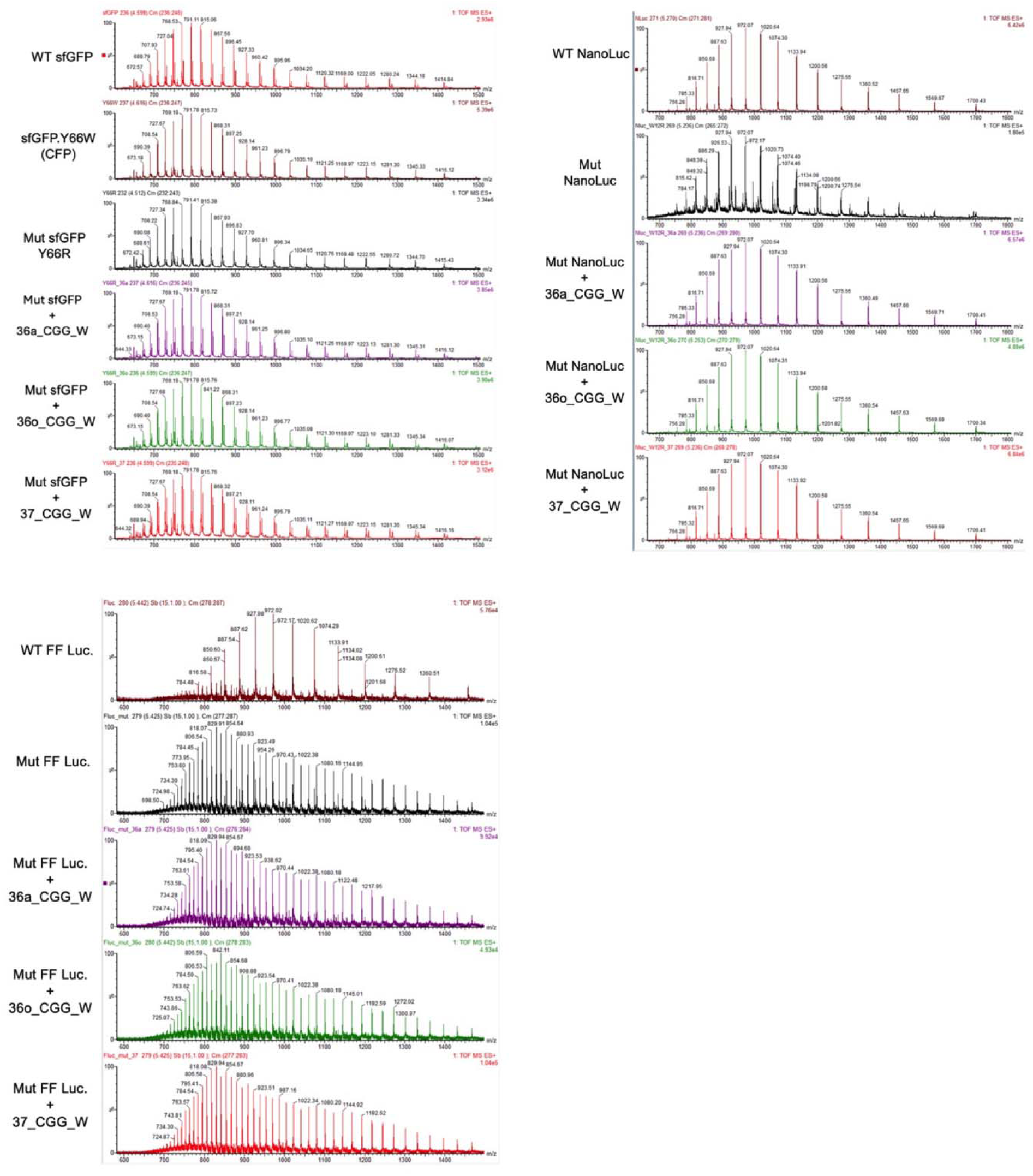
Electrospray ionization charge-state envelopes of purified reporter proteins used for intact-mass determination. Raw positive-ion ESI-TOF mass spectra are shown for sfGFP, NanoLuc, and firefly luciferase (FF Luc.) reporters. Spectra were extracted from the corresponding chromatographic peaks (Figure S10) and display the characteristic series of multiply protonated protein ions. The observed charge-state envelopes were used for subsequent spectral deconvolution and intact-mass determination. The x-axis indicates mass-to-charge ratio (m/z), and the y-axis indicates relative ion intensity.

**Supplementary Figure S12.**
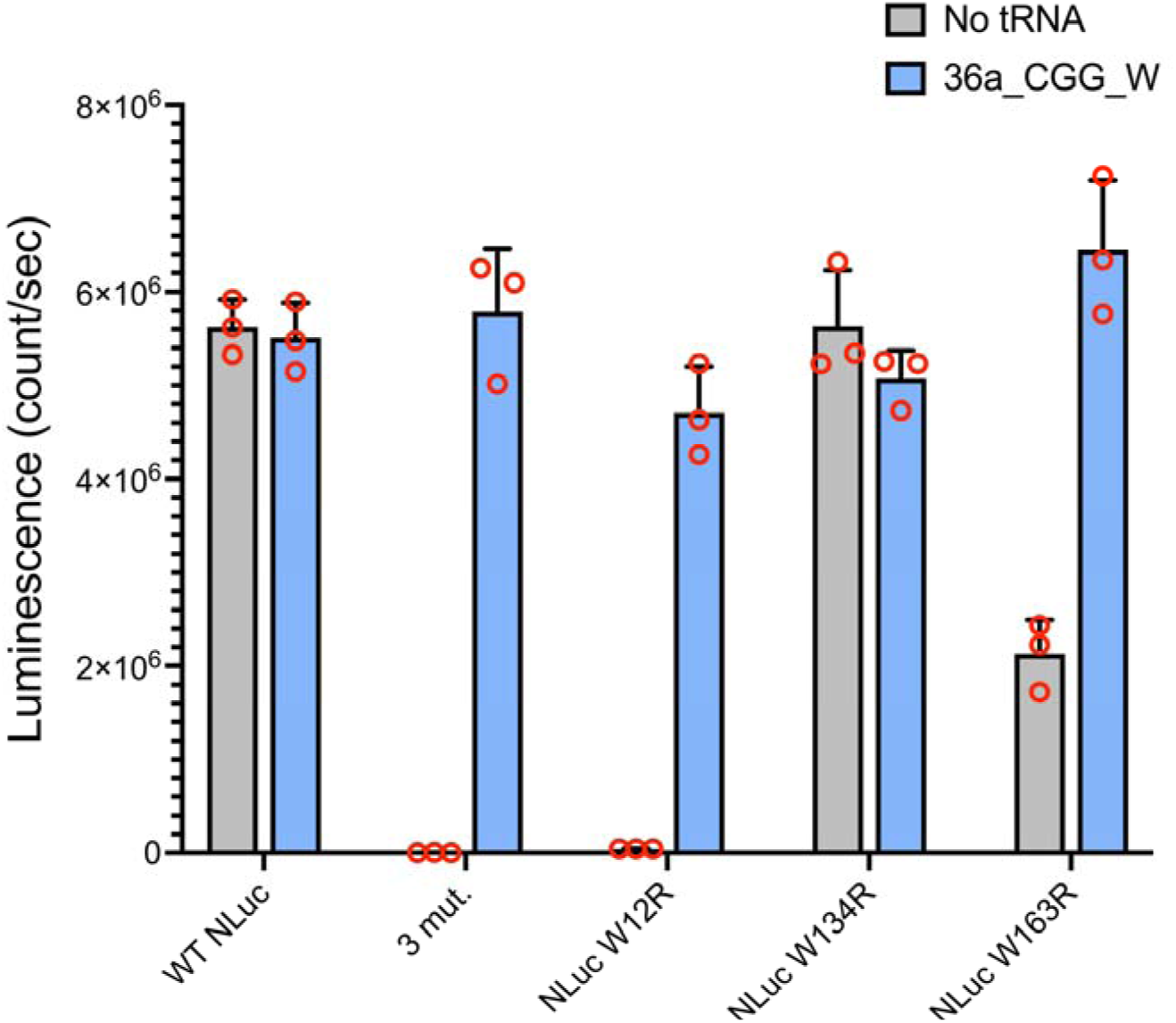
Effects of three NanoLuc mutation sites on activity and rescue by 36a_CGG_W tRNA. NanoLuc luminescence was measured for wild-type NanoLuc (WT Nluc), the three-site mutant (3 mut.), and individual W12R, W134R, and W163R variants in the absence of added tRNA (No tRNA, gray) or in the presence of 36a_CGG_W tRNA (blue). Supplementation with 36a_CGG_W restored luminescence of the three-site mutant and W12R and W163R variants, whereas the effect varied substantially among individual mutation sites. W134R retained relatively high luminescence even without added tRNA. Bars represent mean ± SD of three independent reactions (n = 3), and red circles indicate individual measurements.

**Supplementary Figure S13.**
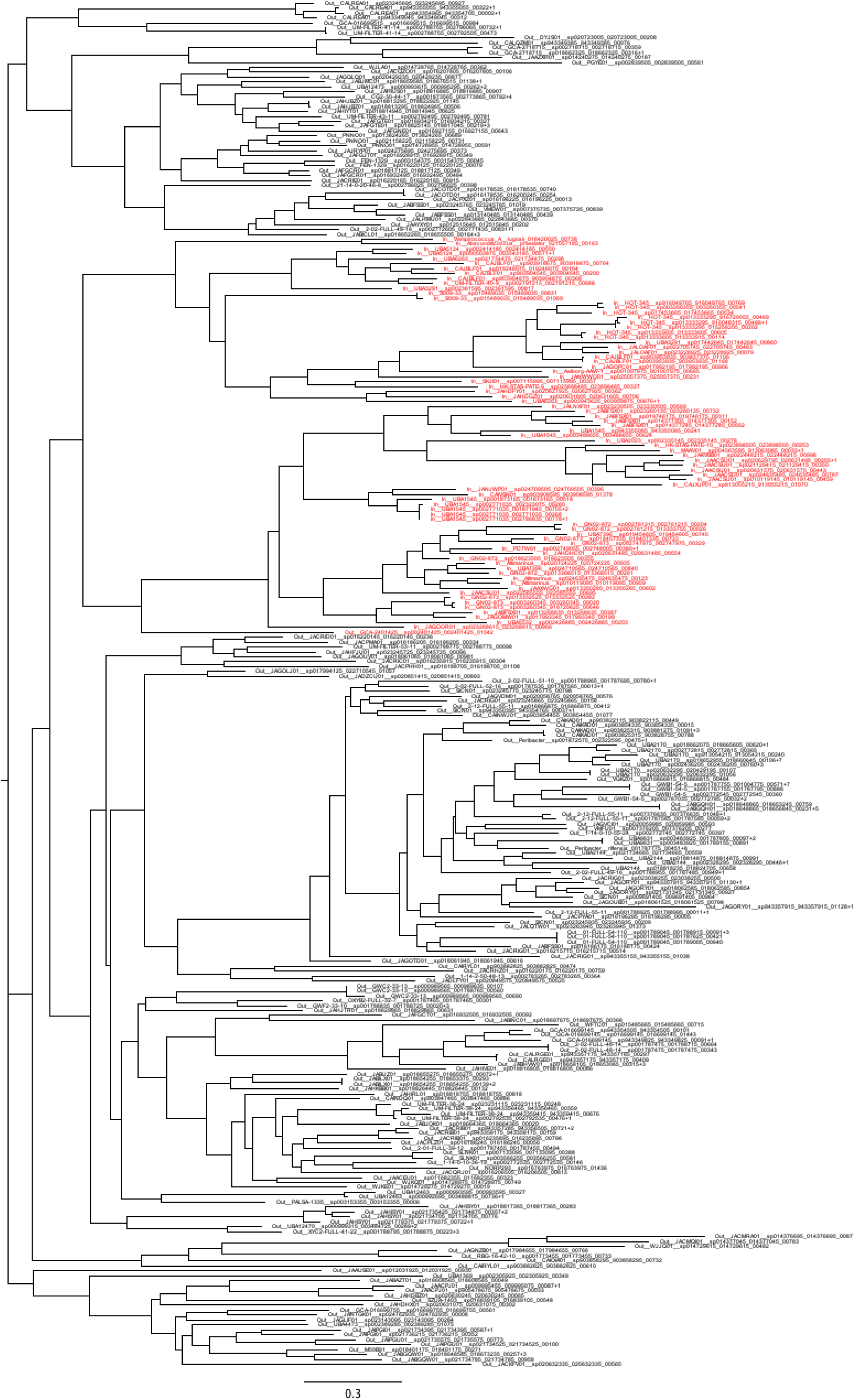
Phylogeny of the GlyRS from genomes of the sister clades using code 25 and 37 (red) and of outgroup genomes using code 11 (black). Trees in Figures S13–S18 are maximum-likelihood phylogenies inferred with RAxML-NG (LG+G8+F, 200 bootstrap replicates) from MAFFT alignments (Materials and methods); scale bars show substitutions per site. “In” and “Out” denote ingroup (code-altered) and outgroup genomes, and Ref_E_coli is the *E. coli* reference sequence.

**Supplementary Figure S14.**
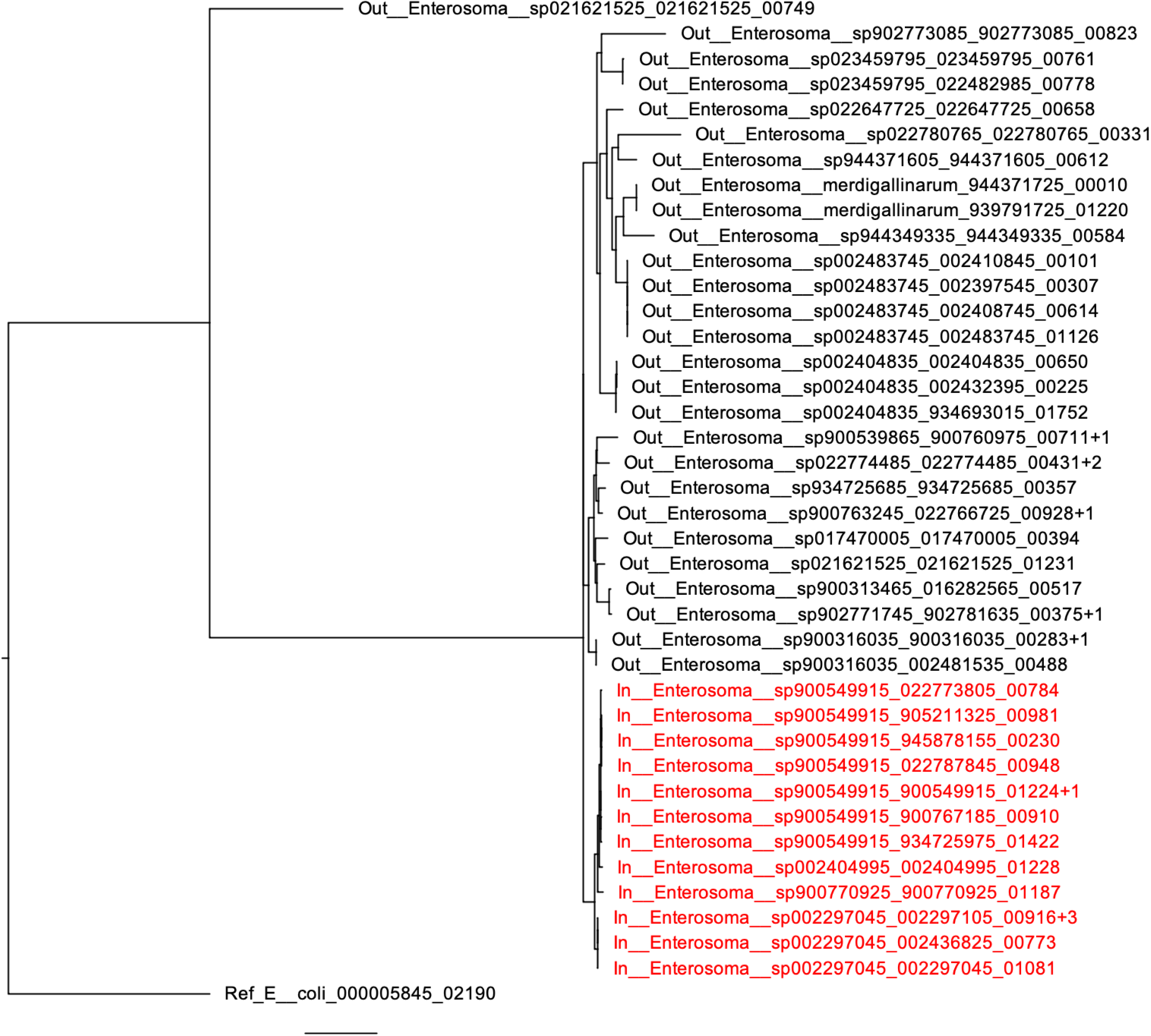
Phylogeny of the MetRS from genomes of the clade using code 34 (red) and of closely related outgroup genomes using code 11 (black), rooting with the reference *E. coli* sequence.

**Supplementary Figure S15.**
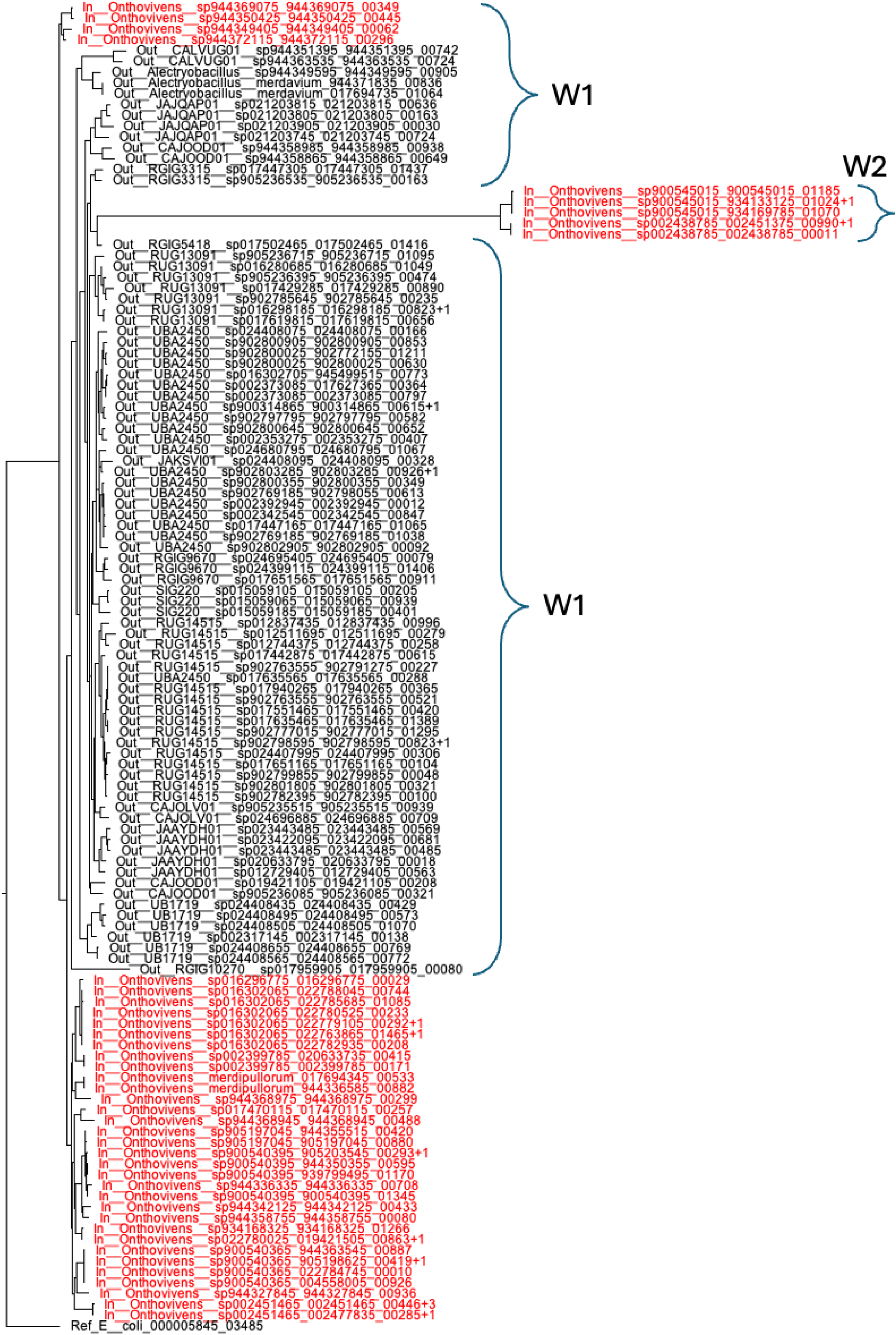
Phylogeny of the TrpRS from genomes of the Onthovivens clade using code 36 (red) and of closely related outgroup genomes using code 11 and reference *E. coli* (black), rooting with the reference *E. coli* sequence.

**Supplementary Figure S16.**
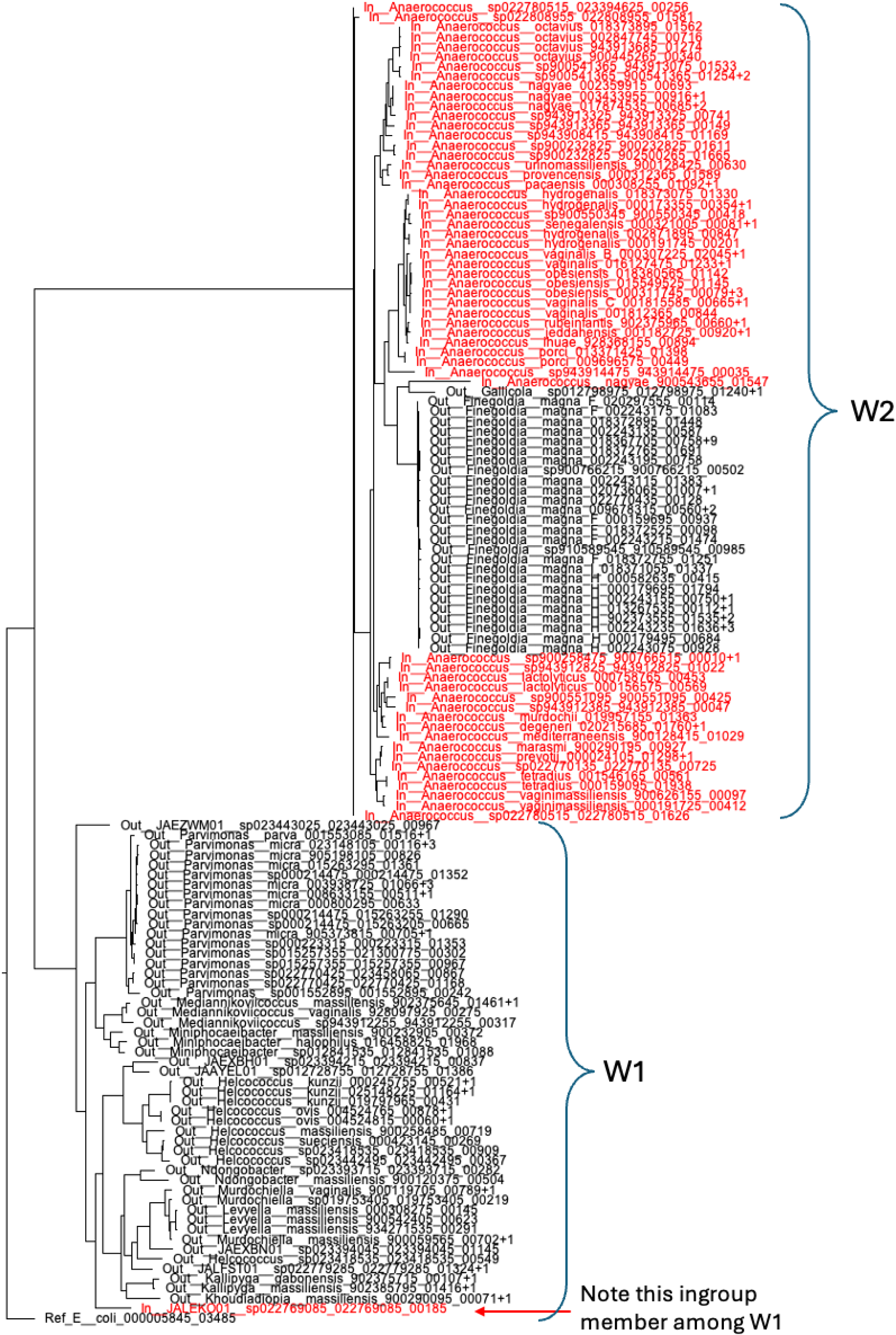
Phylogeny of the TrpRS from genomes of the *Anaerococcus* clade using code 36 (red) and of closely related outgroup genomes u reference *E. coli* sequence. code 11 and reference *E. coli* (black), rooting with the

**Supplementary Figure S17.**
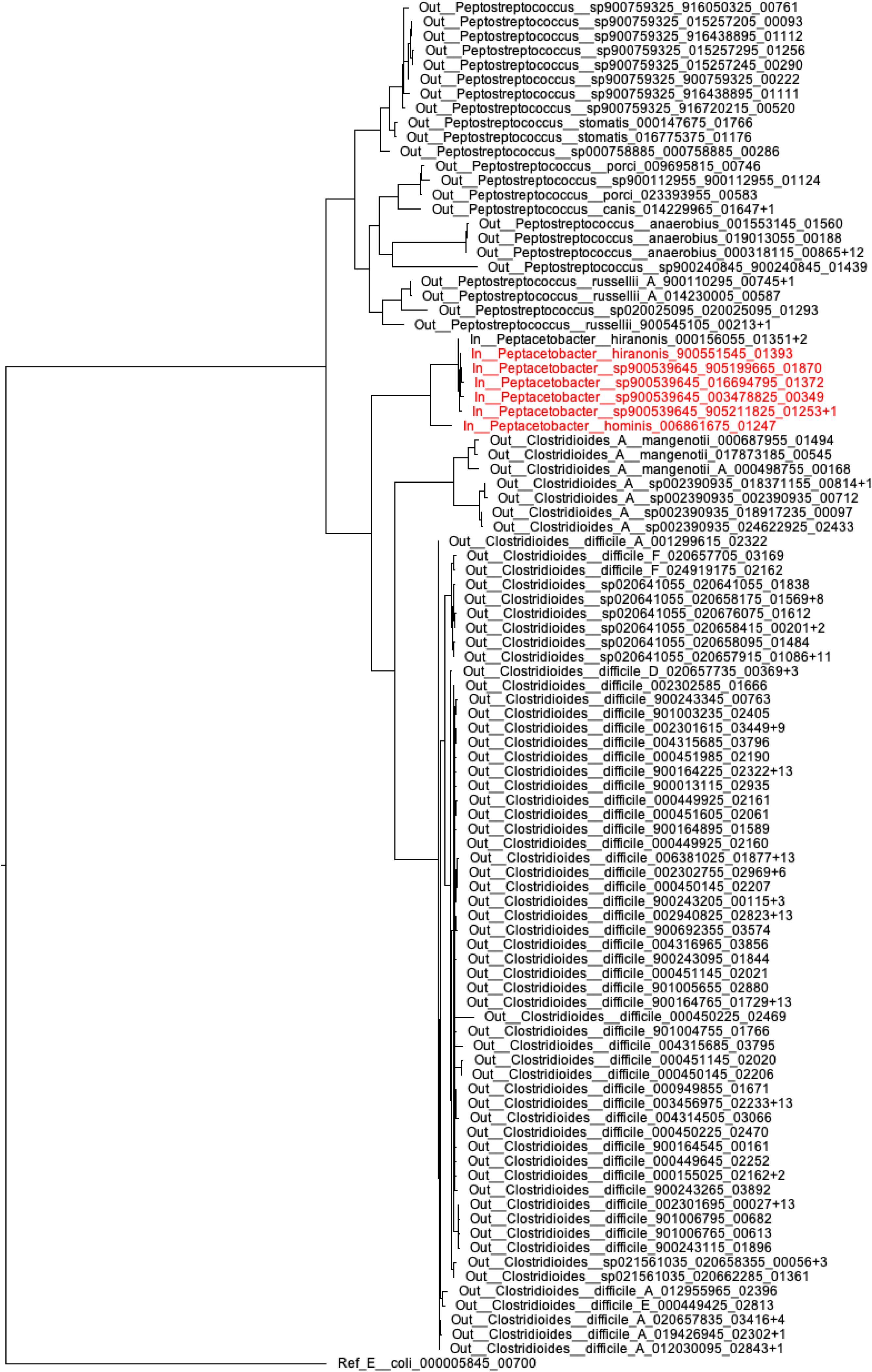
Phylogeny of the GlnRS from genomes of the Peptacetobacter clade using code 35 (red) and of closely related outgroup genomes u reference *E. coli* sequence. code 11 and reference *E. coli* (black), rooting with the

**Supplementary Figure S18.**
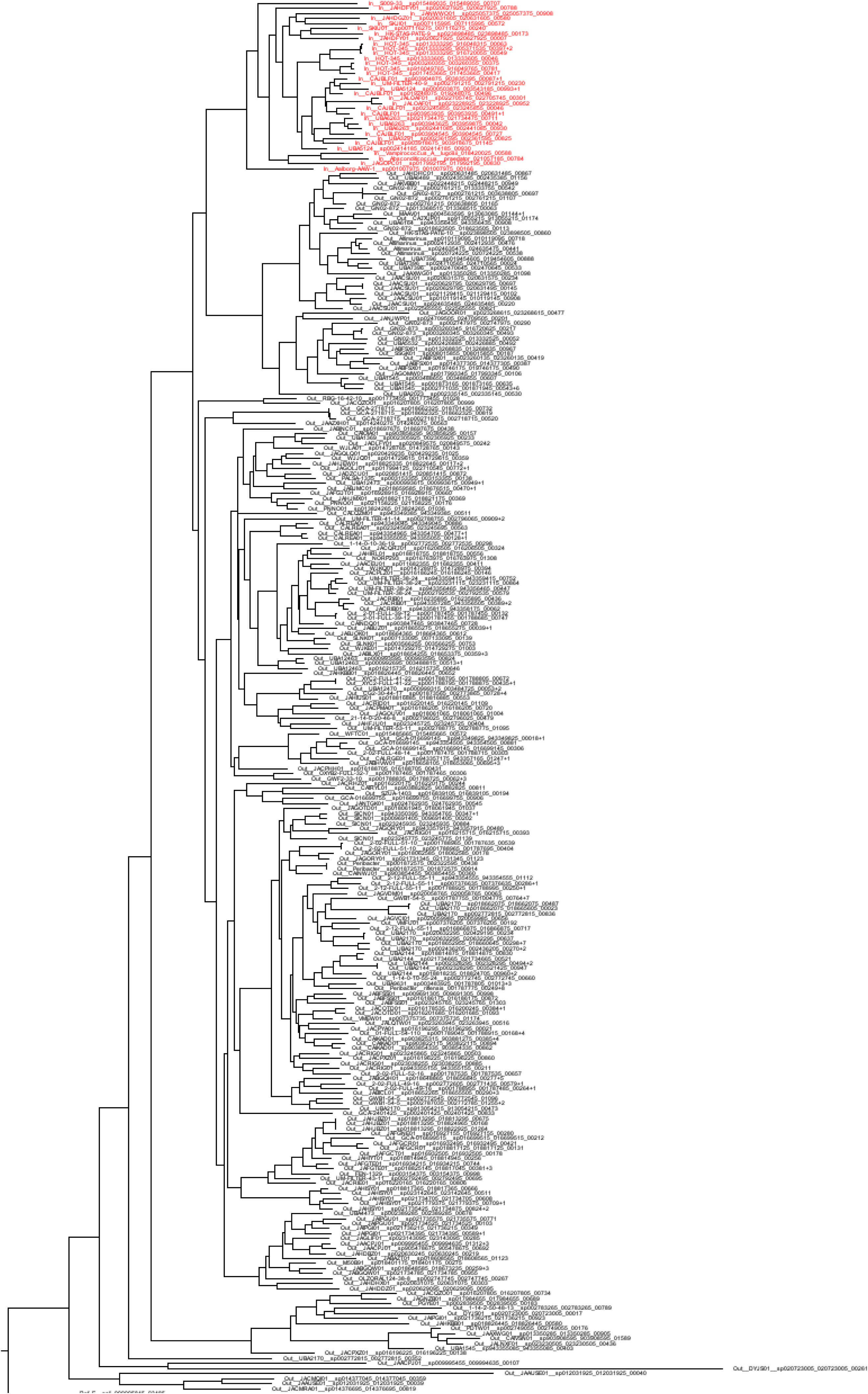
Phylogeny of the TrpRS from genomes of the clade using code 37 (red) and of closely related outgroup genomes using code 11 (black), rooting with the reference *E. coli* sequence.

**Supplementary Figure S19.**
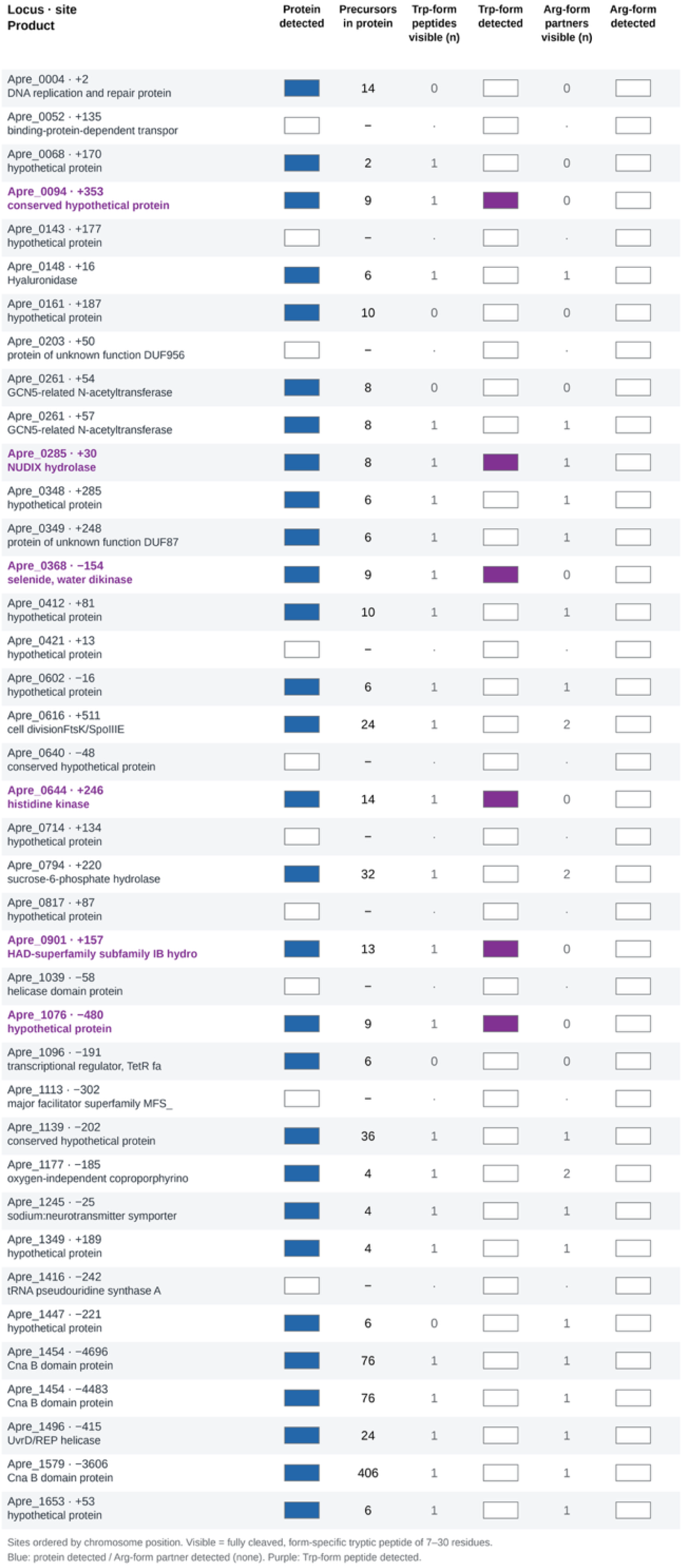
Status of all 39 CGG codons of *A. prevotii*. Sites are ordered by position on chromosome CP001708.1 (+ or − gives the strand of the coding sequence; the number is the amino-acid position). Columns show whether the protein was detected (blue) and its number of precursors; the number of visible Trp-form peptides (fully cleaved tryptic peptides of 7–30 residues that span the site in the CGG→Trp protein); whether a Trp-form peptide was detected (purple); the number of visible Arg-form partner peptides (fully cleaved peptides of 7–30 residues that end at, or begin immediately after, the Arg residue); and whether any of them was detected (none was). Purple loci are the six sites with a Trp-containing peptide.

**Supplementary Figure S20.**
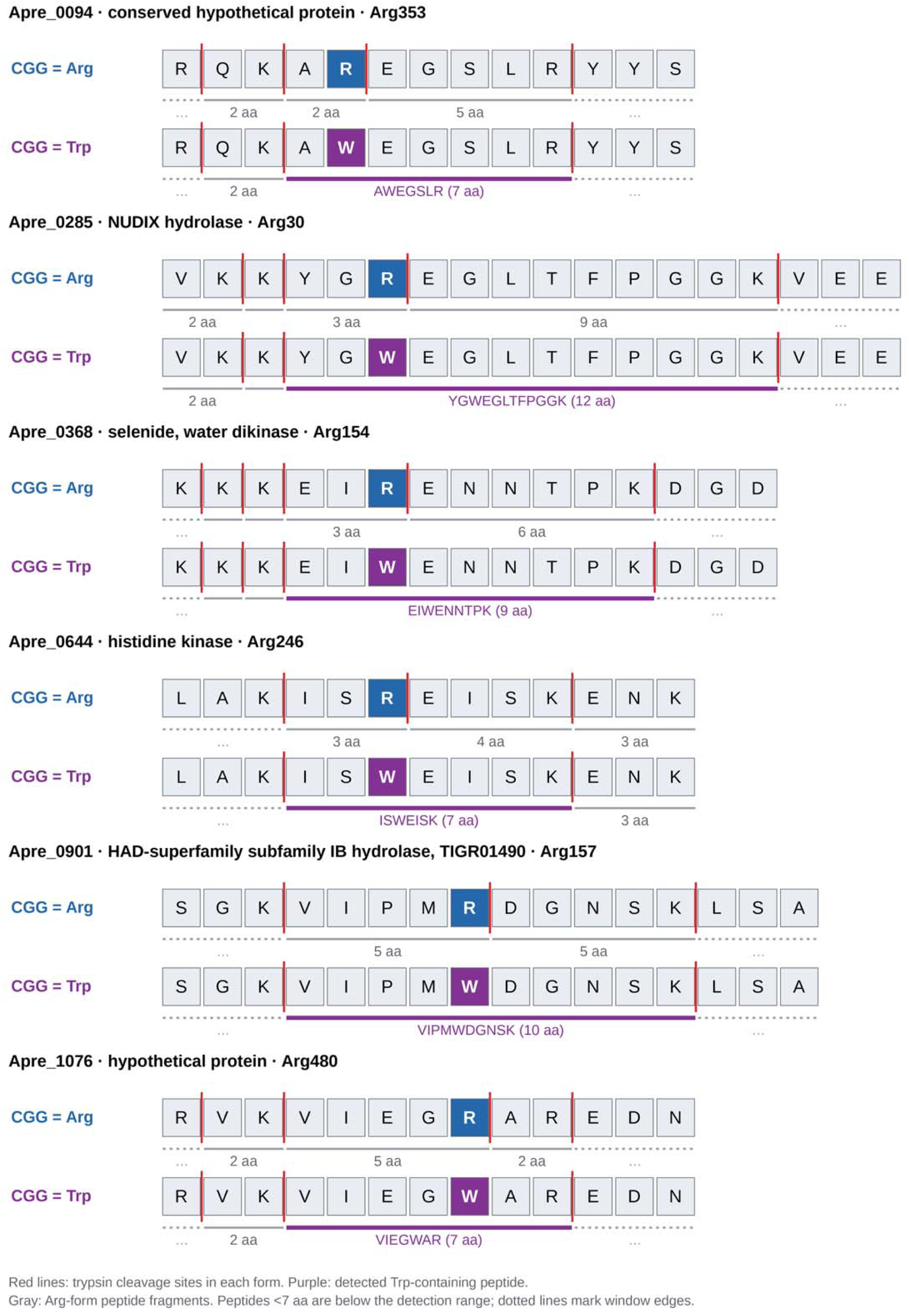
Tryptic peptide maps for the six sites with a detected Trp-containing peptide. For each site, the reference (Arg) and CGG→Trp protein windows are shown with trypsin cleavage sites (red lines) and the resulting peptides (gray bars; the detected Trp-containing peptide is purple). Peptides shorter than 7 residues are below the range of identified peptides (Figure S24A). Only Apre_0285 has a visible Arg-form partner peptide (EGLTFPGGK, 9 residues), and it was not detected. Dotted bars mark the window edge.

**Supplementary Figure S21.**
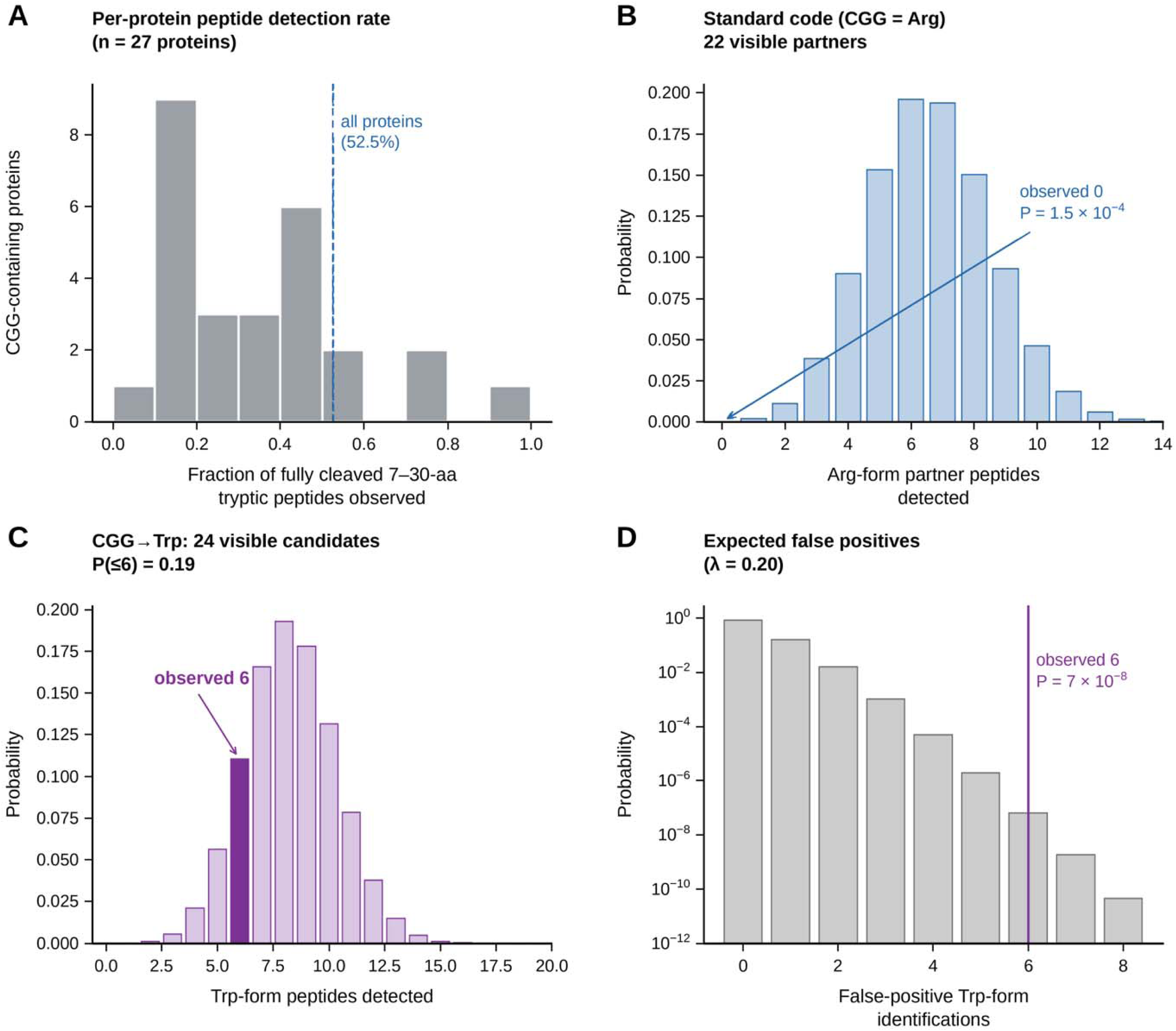
Peptide observability and false-positive estimates. (A) Distribution over the 27 detected CGG-containing proteins of the fraction of fully cleaved 7–30-residue tryptic peptides that were identified (dashed line: 52.5% for all other detected proteins). (B) Probability distribution of the number of Arg-form partner peptides detected if CGG were decoded as Arg (22 visible peptides; Poisson-binomial with per-protein detection probabilities); none was observed (P = 1.5 × 10^−4^). (C) Same for Trp-form peptides if CGG is decoded as Trp (24 visible peptides); six were observed (P(≤6) = 0.19). (D) Poisson distribution of false-positive Trp-form identifications expected at 1% precursor FDR (λ = 0.20, from 151 variant-only peptides × 3 charge states among 125, 492 × 3 library precursors); six were observed (P ≈ 7 × 10^−8^).

**Supplementary Figure S22.**
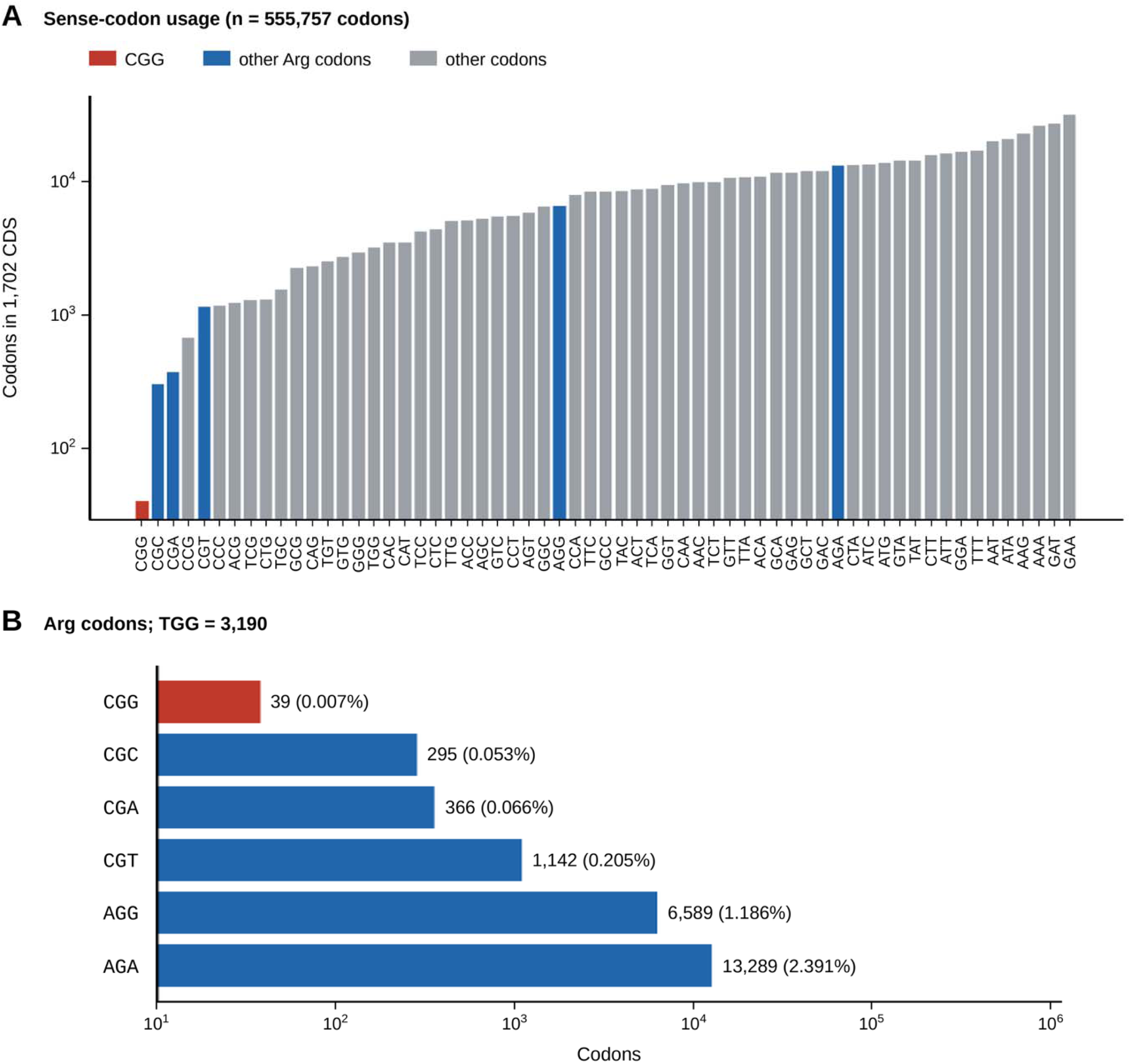
Codon usage in *A. prevotii*. (A) Counts of the 61 sense codons in the 1, 702 annotated coding sequences (555, 757 codons; all CDSs reproduce their annotated translations). CGG (red) is the rarest codon. (B) Arginine codons and TGG (Trp) with counts and percentages of all codons.

**Supplementary Figure S23.**
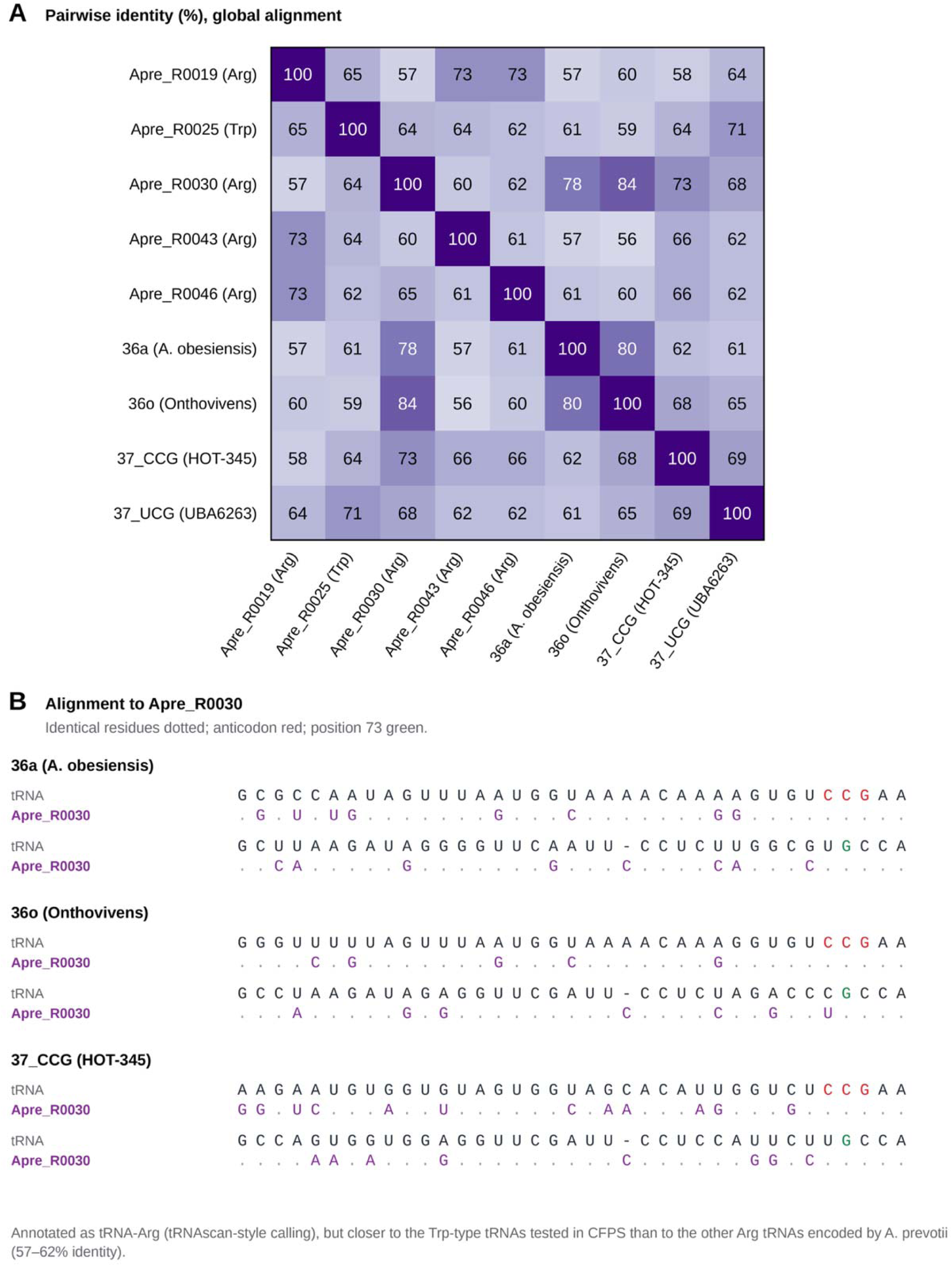
Comparison of *A. prevotii* Arg and Trp tRNAs with the tRNAs tested in CFPS. (A) Pairwise identities (%) from global alignments of the five Arg/Trp tRNA genes of *A. prevotii* and four tested tRNAs (Table S1; 36a, 36o, 37_CCG [37_CGG_W] and 37_UCG [37_CGA_W]). (B) Alignments of 36a, 36o and 37_CCG to Apre_R0030 (the only CCG-anticodon tRNA of *A. prevotii*, annotated as tRNA-Arg); identical positions are dotted, the anticodon is red and position 73 is green.

**Supplementary Figure S24.**
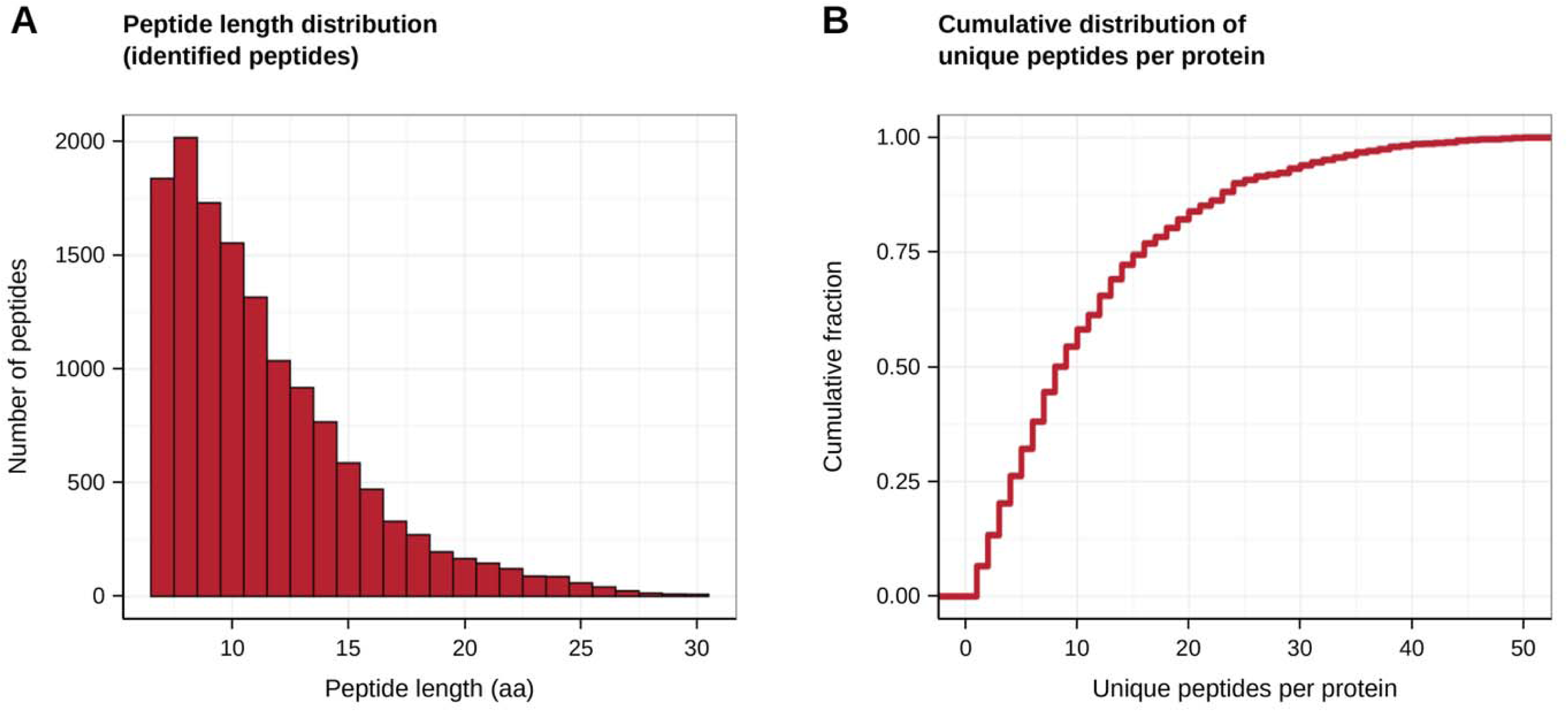
Run-level quality control of the DIA experiment (Novogene). (A) Length distribution of identified peptides (7–30 residues), which defines the visible-peptide range used in the analysis. (B) Cumulative distribution of the number of unique peptides per protein. The single sample (n = 1) allows no replicate-based assessment (CV, PCA). Totals: 13, 800 peptides and 1, 347 protein groups.

**Supplementary Figure S25.**
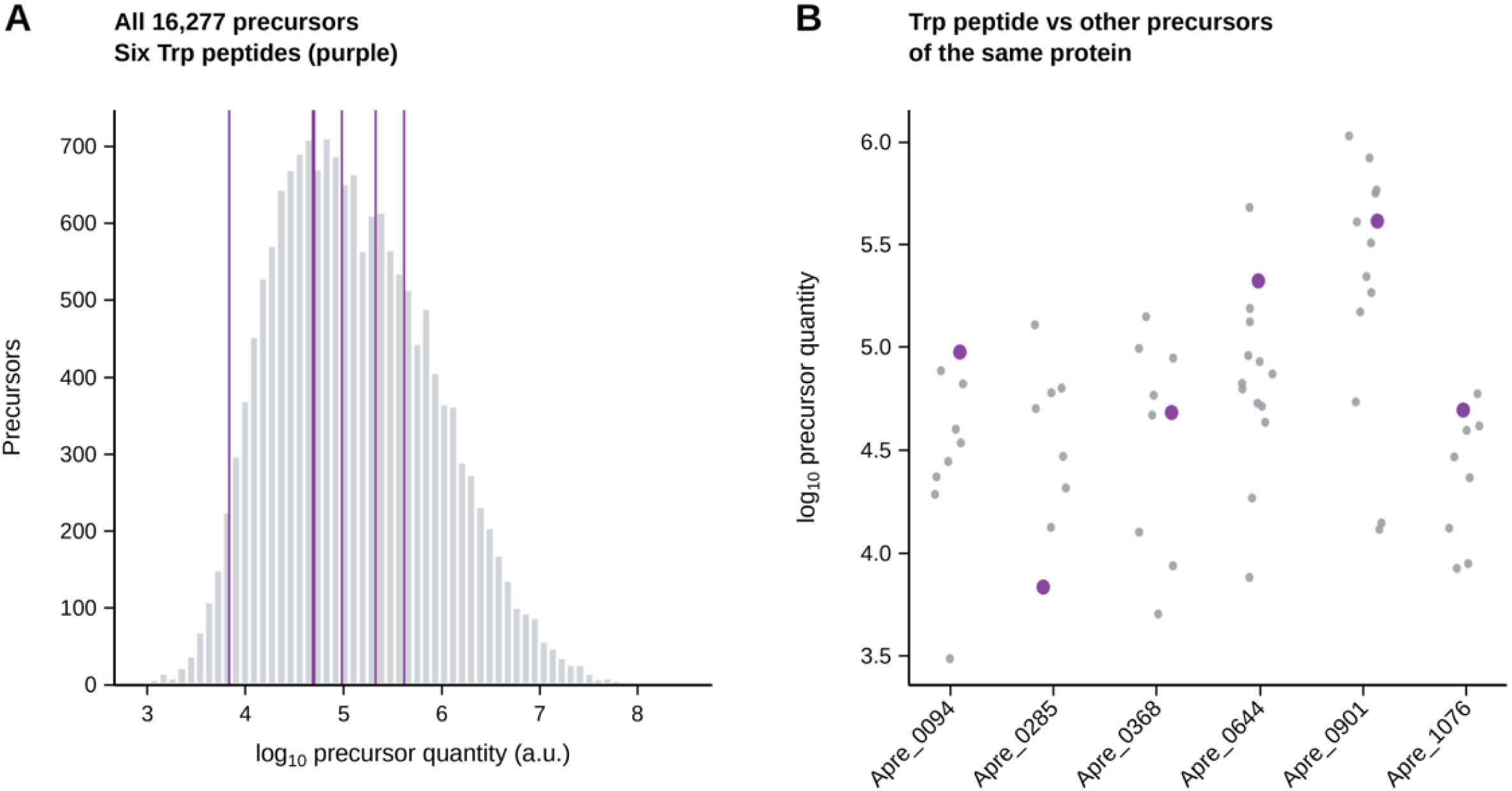
Intensity context of the Trp-containing peptides. (A) Distribution of log_10_ precursor quantities of all 16, 277 precursors in the run; the six Trp-containing precursors are marked in purple. (B) Precursor quantities of each of the six proteins (gray) with the Trp-containing precursor highlighted (purple). Quantities are DIA-NN values (arbitrary units).

**Supplementary Figure S26.**
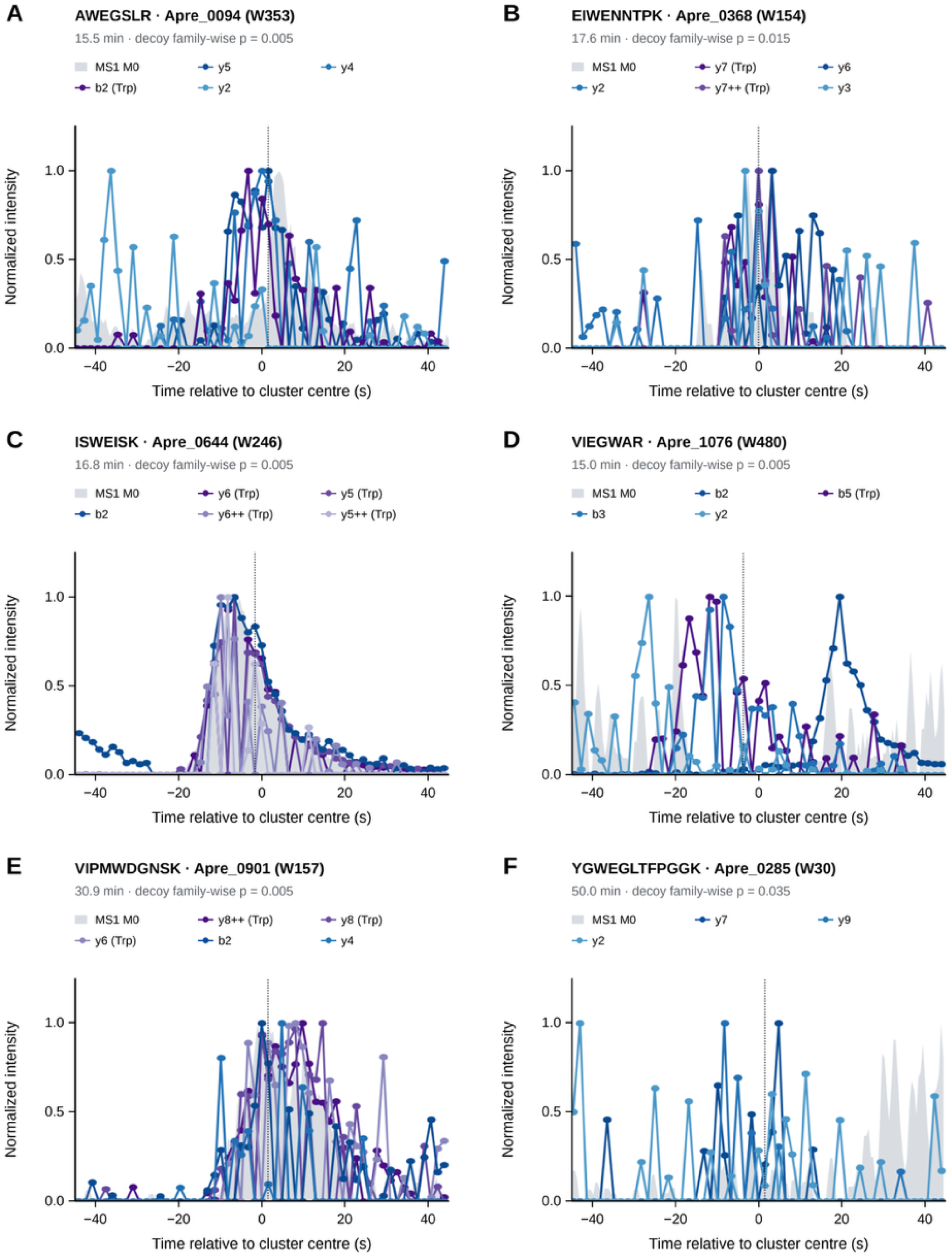
Fragment-ion chromatograms of the six Trp-form peptides, extracted from the raw DIA file. Retention times from the DIA-NN report were not available, so for each peptide the window of five consecutive MS2 scans with the highest fragment-ion score was located from the raw data (Materials and methods); the dotted line marks the scan shown in Figure S27. Lines show up to six b and y ions that were matched within 5 ppm in at least three of the seven scans nearest the cluster center (purple: ions that contain the Trp residue; blue: other ions), each normalized to its maximum within the window. Gray area: MS1 intensity at the monoisotopic m/z (±10 ppm; three-point smoothing). A given isolation window is sampled once every three cycles (about every 1.6 s), so each MS2 trace has few points. P is the family-wise empirical probability that a decoy fragment list scores as high anywhere in the 60-min run (198 decoys). ISWEISK and VIPMWDGNSK show fragment ions that rise and fall with the MS1 signal; the traces of the other four peptides are sparser or less coherent.

**Supplementary Figure S27.**
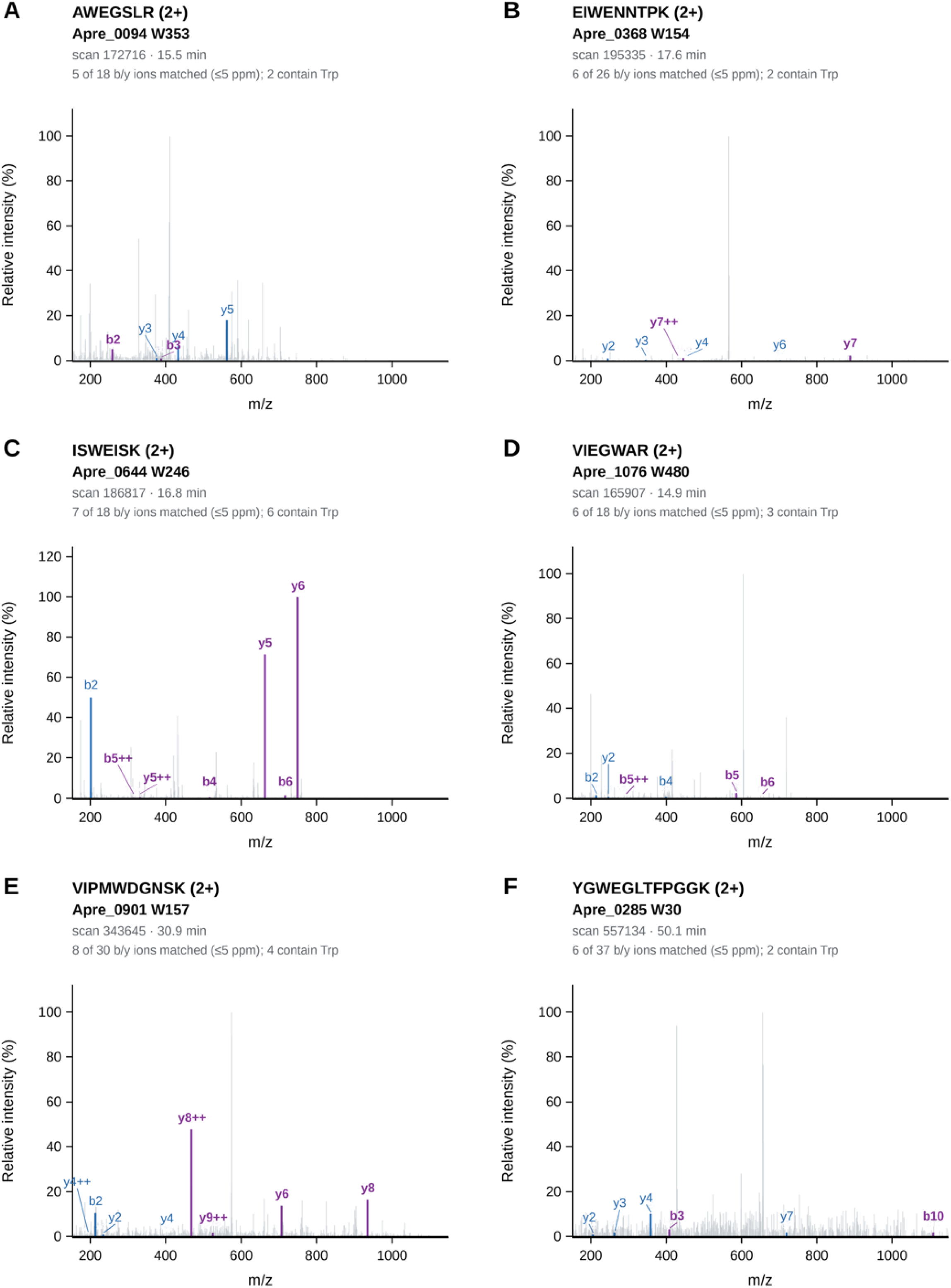
Annotated MS2 spectra of the six Trp-form peptides. Highest-scoring MS2 scan of the cluster shown in Figure S26 for each peptide (doubly charged precursor; isolation window 2 Th). Purple: matched b and y ions (≤5 ppm) that contain the Trp residue; blue: other matched ions; gray: unmatched peaks. The tall unlabeled peak near the center of the isolation window in several panels is the unfragmented precursor or a co-isolated ion. The number of matched ions excludes b1 and y1 and counts 1+ and 2+ ions separately. For ISWEISK, the three most intense peaks match y6, y5 and b2; for VIPMWDGNSK, y8 (2+ and 1+) and y6 are matched among the most intense peaks; in the other panels matched ions are of low intensity.

**Supplementary Figure S28.**
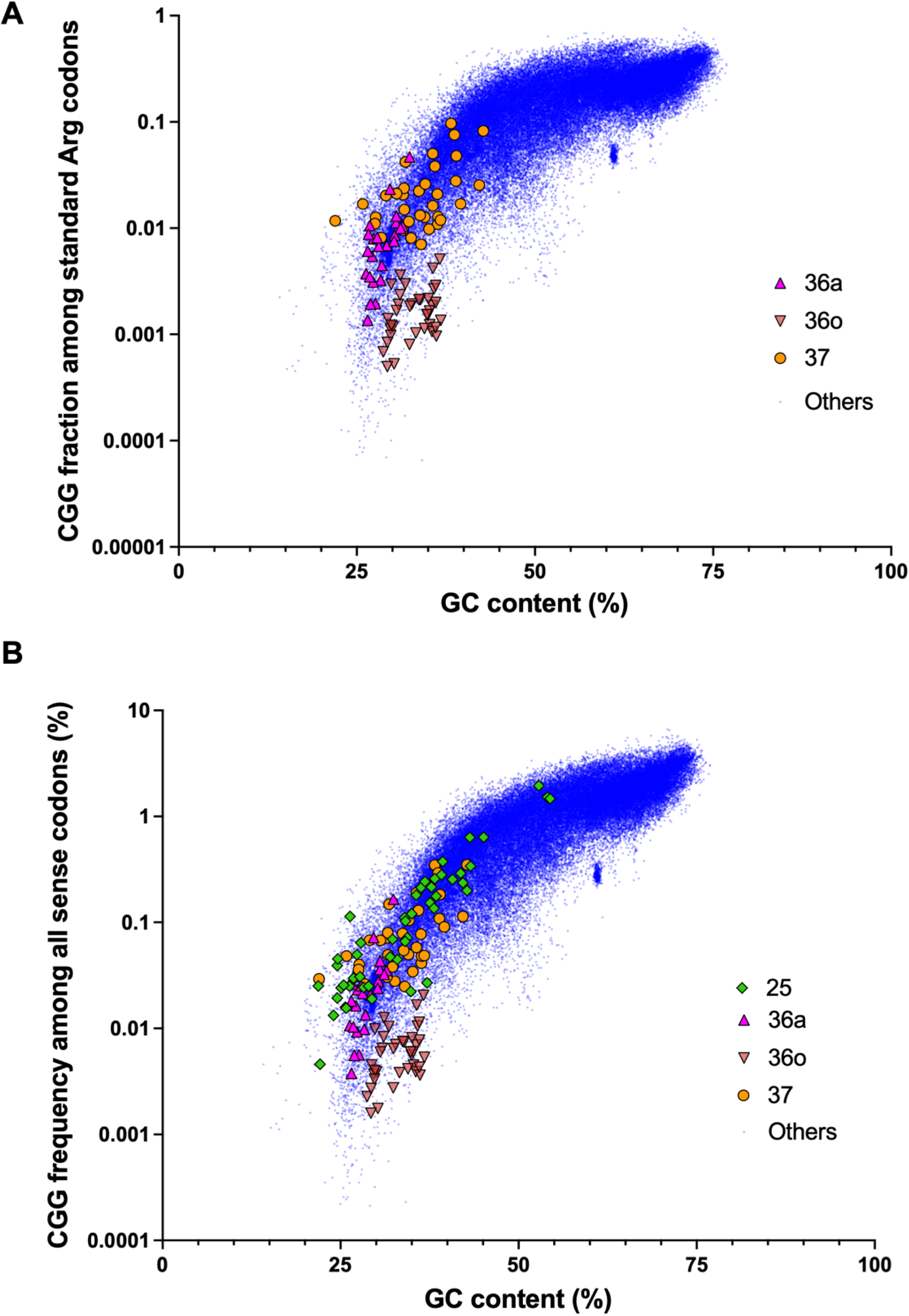
CGG codon usage as a function of genomic GC content. Each point is a genome. CGG abundance is relative to (A) standard Arg codons or (B) all standard codons. Symbols: 36a, magenta upward triangles; 36o, salmon downward triangles; 37, orange circles; 25, green diamonds (B only); others, blue points. Both y-axes are logarithmic (base 10).

**Supplementary Table S1.**
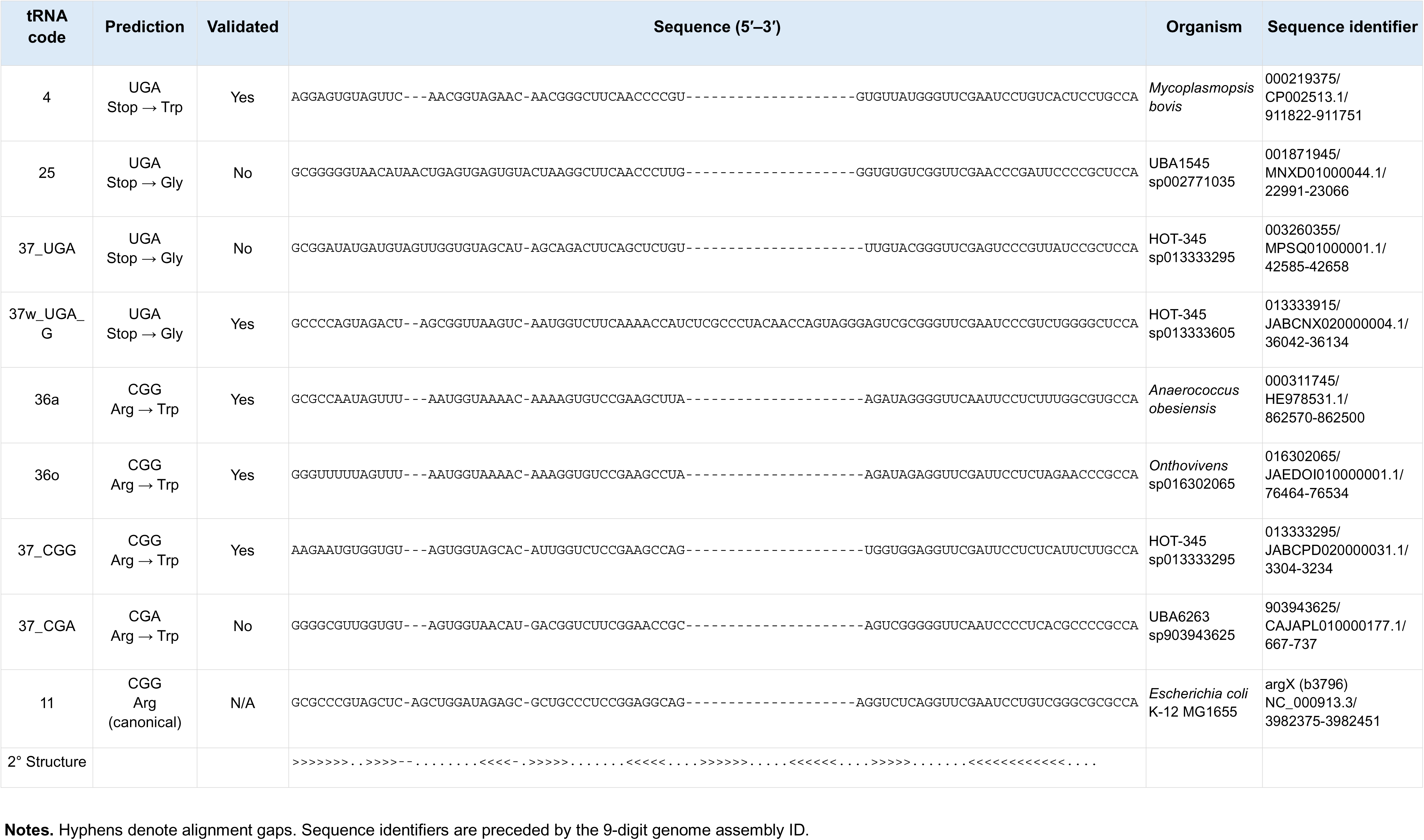
Alternative tRNAs used in this study.

**Supplementary Table S2.**
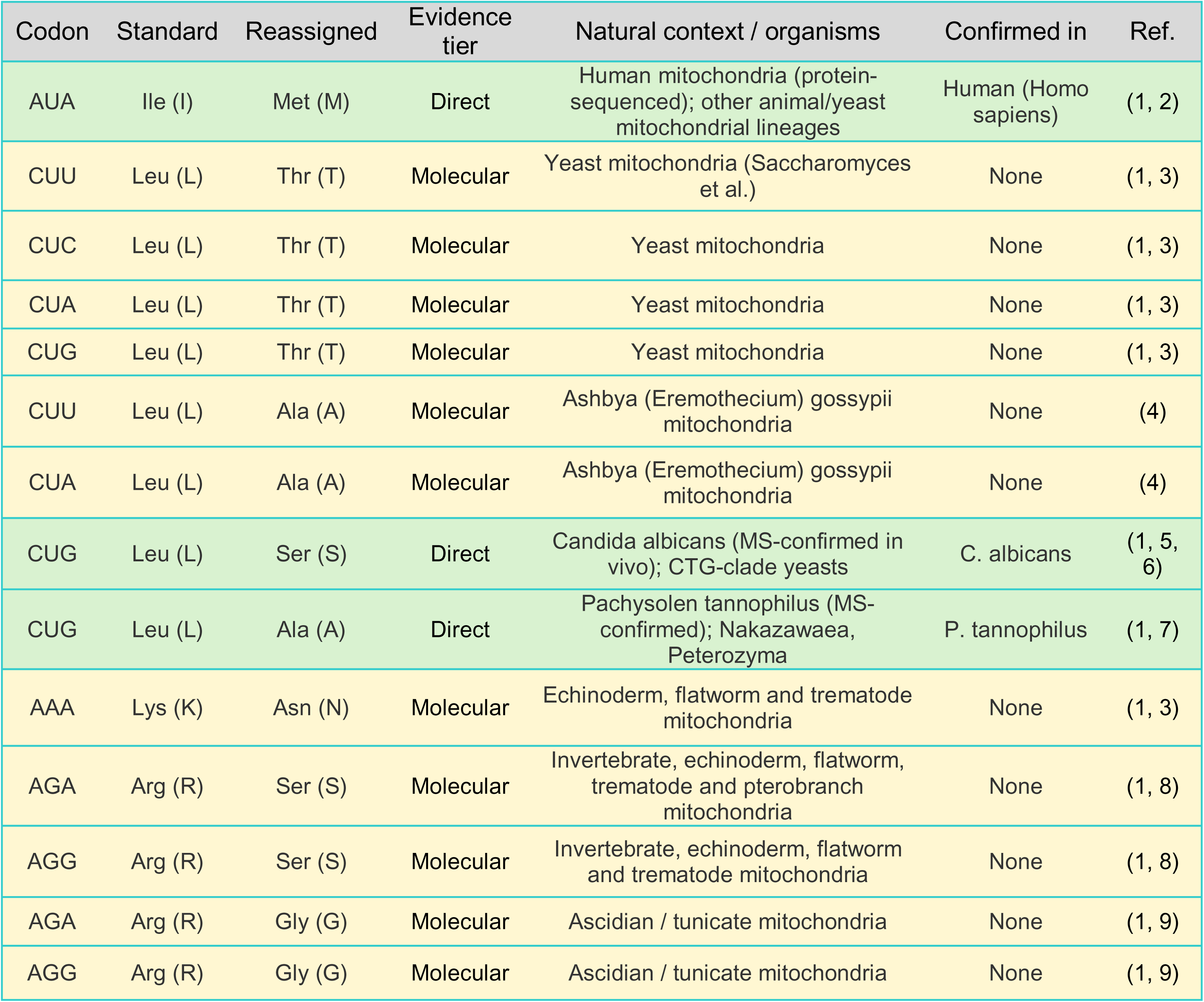
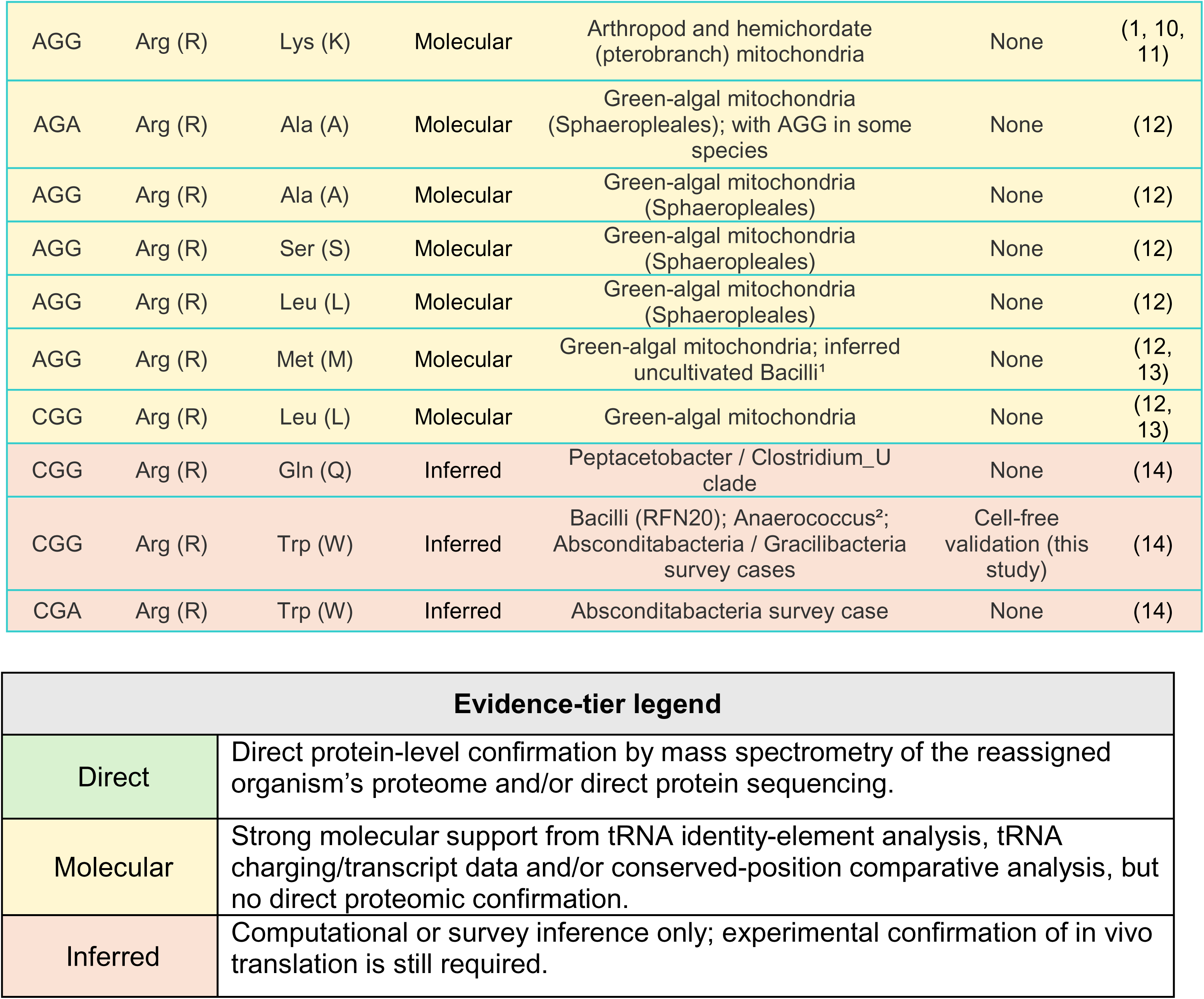
Summary of known and predicted sense-to-sense codon reassignments. Reported sense-to-sense codon reassignments are listed with the canonical amino acid assignment, proposed reassigned amino acid, evidence tier, natural context or taxonomic range, experimentally confirmed organism when available and supporting references. Evidence tiers indicate the strength of support for each reassignment; Direct: direct protein-level confirmation by mass spectrometry or protein sequencing; Molecular: strong molecular support such as tRNA identity-element analysis, tRNA charging or transcript evidence, and/or conserved-position comparative analysis without direct proteomic confirmation; Inferred: computational or survey-based inference only. The natural context/organism column indicates the biological systems in which each reassignment has been reported or predicted. The bacterial CGG Arg→Trp assignments (codes 36 and 37) were tested experimentally in this study by cell-free translation (main text).

| Codon | Standard | Reassigned | Evidence tier | Natural context / organisms | Confirmed in | Ref. |
|---|---|---|---|---|---|---|
| AUA | Ile (I) | Met (M) | Direct | Human mitochondria (protein-sequenced); other animal/yeast mitochondrial lineages | Human (Homo sapiens) | (1, 2) |
| CUU | Leu (L) | Thr (T) | Molecular | Yeast mitochondria (Saccharomyces et al.) | None | (1, 3) |
| CUC | Leu (L) | Thr (T) | Molecular | Yeast mitochondria | None | (1, 3) |
| CUA | Leu (L) | Thr (T) | Molecular | Yeast mitochondria | None | (1, 3) |
| CUG | Leu (L) | Thr (T) | Molecular | Yeast mitochondria | None | (1, 3) |
| CUU | Leu (L) | Ala (A) | Molecular | Ashbya (Eremothecium) gossypii mitochondria | None | (4) |
| CUA | Leu (L) | Ala (A) | Molecular | Ashbya (Eremothecium) gossypii mitochondria | None | (4) |
| CUG | Leu (L) | Ser (S) | Direct | Candida albicans (MS-confirmed in vivo); CTG-clade yeasts | C. albicans | (1, 5, 6) |
| CUG | Leu (L) | Ala (A) | Direct | Pachysolen tannophilus (MS-confirmed); Nakazawaea, Peterozyma | P. tannophilus | (1, 7) |
| AAA | Lys (K) | Asn (N) | Molecular | Echinoderm, flatworm and trematode mitochondria | None | (1, 3) |
| AGA | Arg (R) | Ser (S) | Molecular | Invertebrate, echinoderm, flatworm, trematode and pterobranch mitochondria | None | (1, 8) |
| AGG | Arg (R) | Ser (S) | Molecular | Invertebrate, echinoderm, flatworm and trematode mitochondria | None | (1, 8) |
| AGA | Arg (R) | Gly (G) | Molecular | Ascidian / tunicate mitochondria | None | (1, 9) |
| AGG | Arg (R) | Gly (G) | Molecular | Ascidian / tunicate mitochondria | None | (1, 9) |
| AGG | Arg (R) | Lys (K) | Molecular | Arthropod and hemichordate (pterobranch) mitochondria | None | (1, 10, 11) |
| AGA | Arg (R) | Ala (A) | Molecular | Green-algal mitochondria (Sphaeropleales); with AGG in some species | None | (12) |
| AGG | Arg (R) | Ala (A) | Molecular | Green-algal mitochondria (Sphaeropleales) | None | (12) |
| AGG | Arg (R) | Ser (S) | Molecular | Green-algal mitochondria (Sphaeropleales) | None | (12) |
| AGG | Arg (R) | Leu (L) | Molecular | Green-algal mitochondria (Sphaeropleales) | None | (12) |
| AGG | Arg (R) | Met (M) | Molecular | Green-algal mitochondria; inferred uncultivated Bacilli<sup>1</sup> | None | (12, 13) |
| CGG | Arg (R) | Leu (L) | Molecular | Green-algal mitochondria | None | (12, 13) |
| CGG | Arg (R) | Gln (Q) | Inferred | Peptacetobacter / Clostridium_U clade | None | (14) |
| CGG | Arg (R) | Trp (W) | Inferred | Bacilli (RFN20); Anaerococcus<sup>2</sup>; Absconditabacteria / Gracilibacteria survey cases | Cell-free validation (this study) | (14) |
| CGA | Arg (R) | Trp (W) | Inferred | Absconditabacteria survey case | None | (14) |

**Evidence-tier legend**
Direct: Direct protein-level confirmation by mass spectrometry of the reassigned organism’s proteome and/or direct protein sequencing.
Molecular: Strong molecular support from tRNA identity-element analysis, tRNA charging/transcript data and/or conserved-position comparative analysis, but no direct proteomic confirmation.
Inferred: Computational or survey inference only; experimental confirmation of in vivo translation is still required.

**Supplementary Table S3.** A list of reagents used to prepare cell-free extract and CFPS reactions.

| Reagent/Item | Product # | Notes/Vendor |
|---|---|---|
| ATP disodium salt hydrate | A2383-1G | Sigma |
| GTP sodium salt hydrate | G8877-1G | Sigma |
| UTP trisodium salt hydrate | U6625-1G | Sigma |
| CTP disodium salt | C1506-1G | Sigma |
| Magnesium glutamate | 49605-250G | Sigma |
| Potassium glutamate | G1501-100G | Sigma |
| Ammonium Glutamate | n/a | Jewett Lab |
| Creatine phosphate | 27920-5G | Sigma |
| Putrescine | P5780-5G | Sigma |
| Spermidine | S0266-5G | Sigma |
| Creatine phosphokinase | C3755-3.5KU | Sigma |
| Folinic acid | 47612-250MG | Sigma |
| tRNA | n/a | Jewett Lab |
| Coenzyme A | C4282-100MG | Sigma |
| NAD | N8535-15VL | Sigma |
| Potassium oxalate | P0963-100G | Sigma |
| PEP | 10108294001 | Sigma |
| TCEP | 77720 | Thermo |
| Potassium phosphate dibasic | P3786-100G | Sigma |
| potassium phosphate monobasic | P5655-100G | Sigma |
| Magnesium acetate tetrahydrate | 228648-100G | Sigma |
| Tris base | 252859-100G | Sigma |
| L-Isoleucine | I2752-25G | Sigma |
| L-Histidine | H8000-25G | Sigma |
| L-Serine | S4500-100G | Sigma |
| L-Proline | P0380-100G | Sigma |
| L-Glutamine | G3126-100G | Sigma |
| L-Leucine | L8000-50G | Sigma |
| L-Aspartic Acid | A9256-100G | Sigma |
| L-arginine | A0884-25G | Sigma |
| L-glycine | G7126-100G | Sigma |
| L-lysine | L5501-25G | Sigma |
| L-methionine | M9625-25G | Sigma |
| L-phenylalanine | P2126-100G | Sigma |
| L-threonine | T8625-25G | Sigma |
| L-tyrosine | T3754-50G | Sigma |
| L-Tryptophan | T8941-25G | Sigma |
| L-Valine | V0513-25G | Sigma |
| L-Alanine | A7469-25G | Sigma |
| L-Cysteine | C7352-25G | Sigma |
| L-Asparagine monohydrate | A7094-100G | Sigma |
| L-Glutamic Acid | G8415-100G | Sigma |
| 2x YT media | Y2377-250G | Sigma |
| Glucose | G7021-1KG | Sigma |

**Supplementary Table S4.**
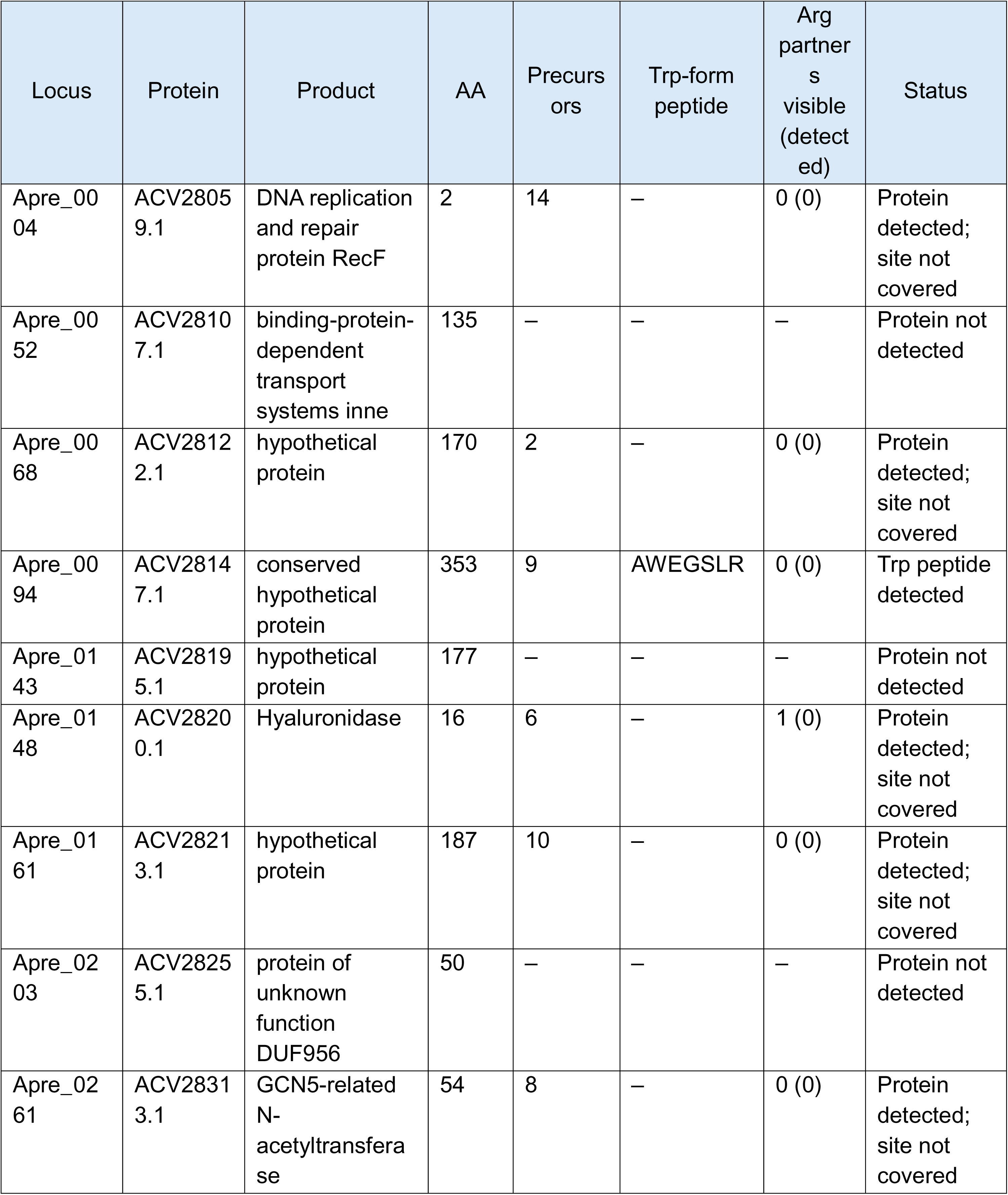
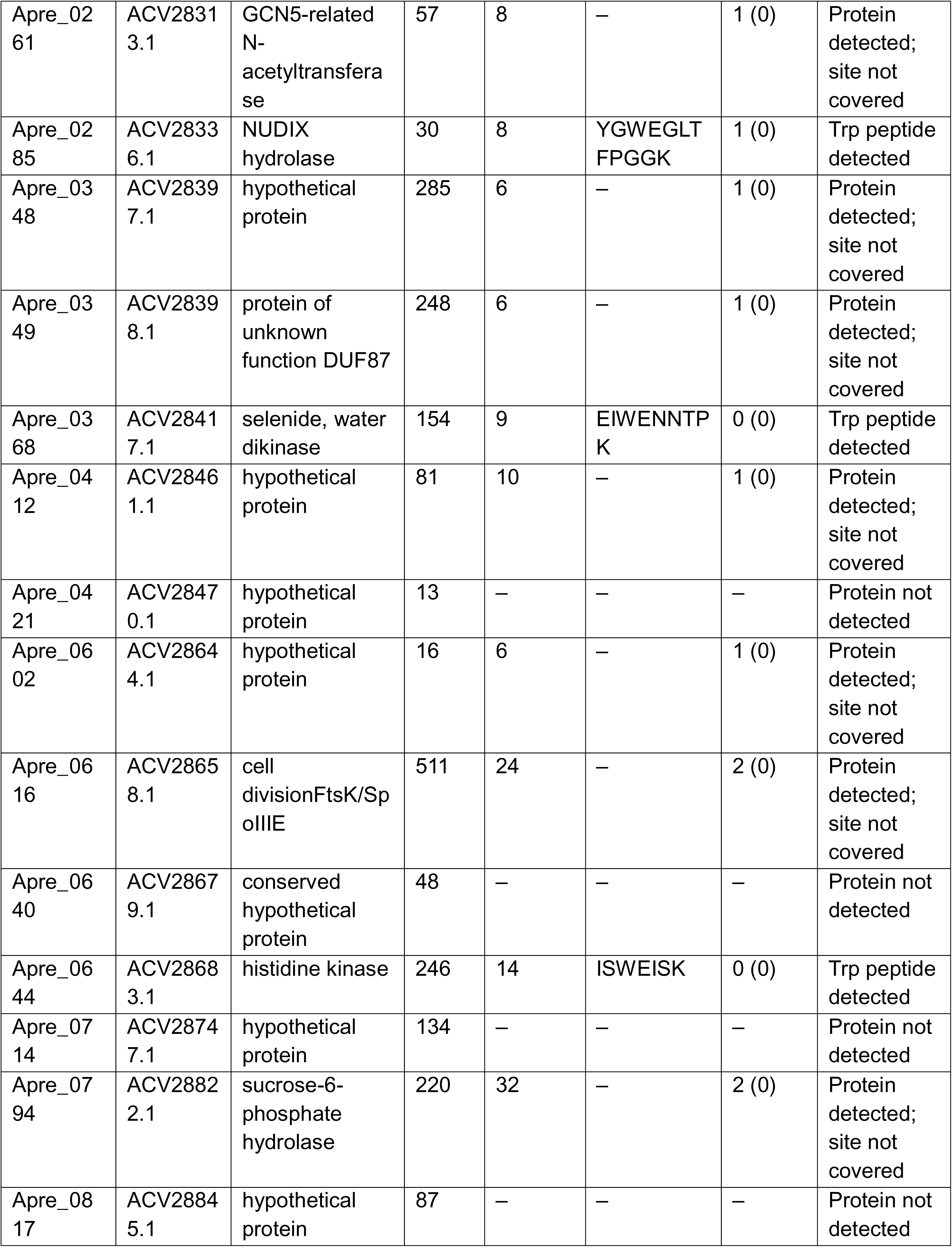
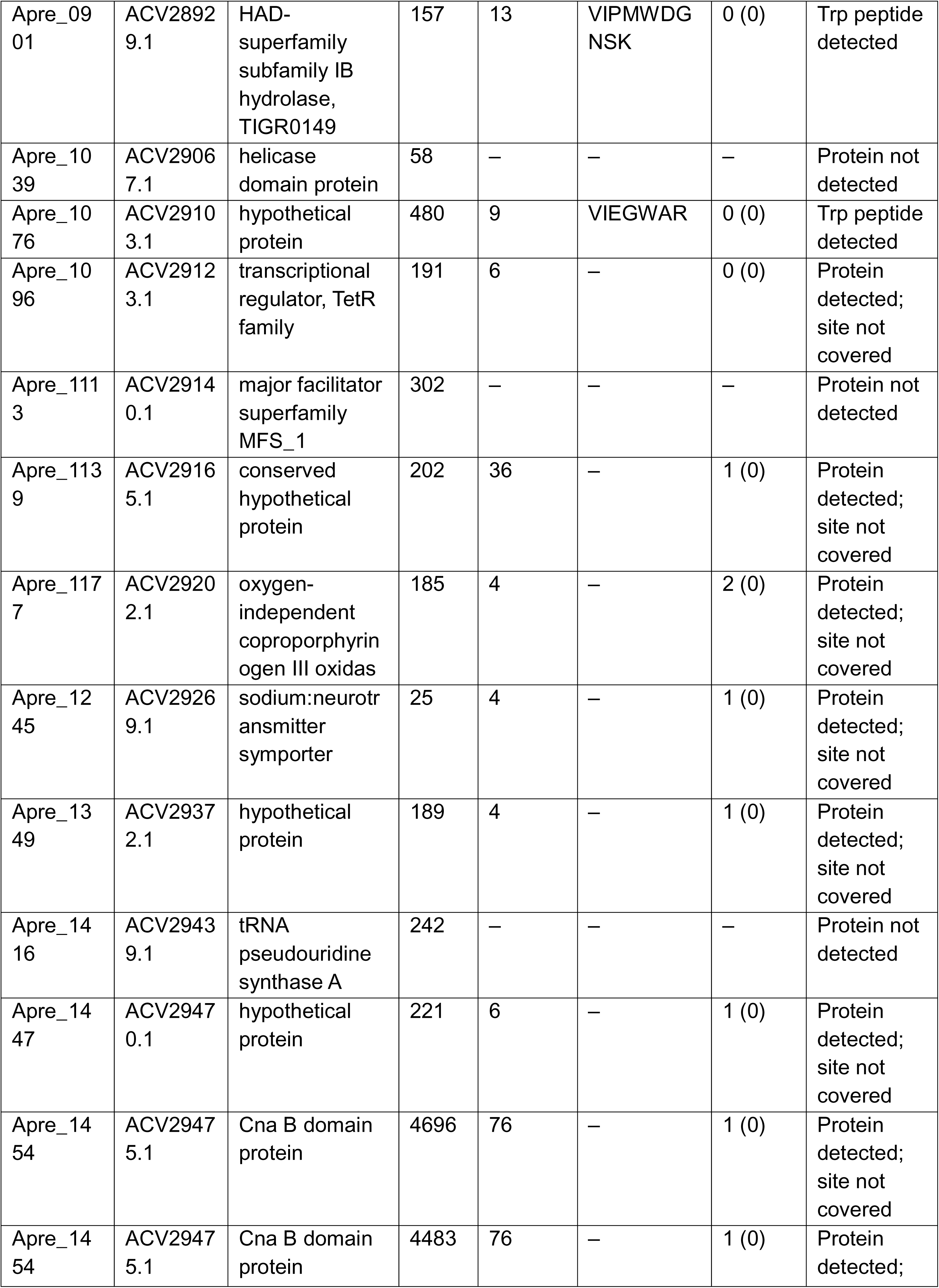
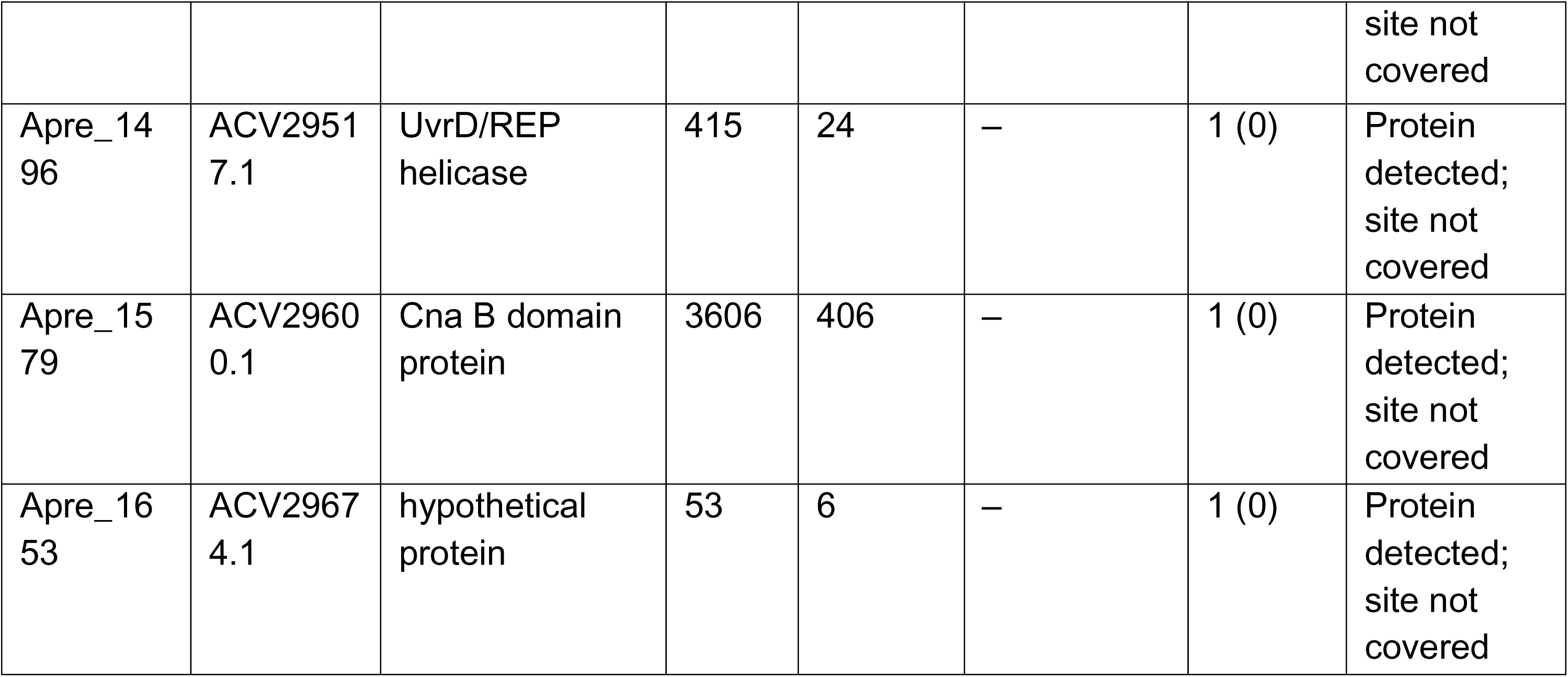
The 39 in-frame CGG codons of *A. prevotii* and their proteomic coverage. Product names are from the GenBank record (truncated). “Arg partners visible (detected)”: fully cleaved Arg-form peptides of 7–30 residues that end at, or begin after, the residue at the site; none was detected. Full table: Supplementary Data workbook.

| Locus | Protein | Product | AA | Precursors | Trp-form peptide | Arg partners visible (detected) | Status |
|---|---|---|---|---|---|---|---|
| Apre_0004 | ACV28059.1 | DNA replication and repair protein RecF | 2 | 14 | – | 0 (0) | Protein detected; site not covered |
| Apre_0052 | ACV28107.1 | binding-protein-dependent transport systems inne | 135 | – | – | – | Protein not detected |
| Apre_0068 | ACV28122.1 | hypothetical protein | 170 | 2 | – | 0 (0) | Protein detected; site not covered |
| Apre_0094 | ACV28147.1 | conserved hypothetical protein | 353 | 9 | AWEGSLR | 0 (0) | Trp peptide detected |
| Apre_0143 | ACV28195.1 | hypothetical protein | 177 | – | – | – | Protein not detected |
| Apre_0148 | ACV28200.1 | Hyaluronidase | 16 | 6 | – | 1 (0) | Protein detected; site not covered |
| Apre_0161 | ACV28213.1 | hypothetical protein | 187 | 10 | – | 0 (0) | Protein detected; site not covered |
| Apre_0203 | ACV28255.1 | protein of unknown function DUF956 | 50 | – | – | – | Protein not detected |
| Apre_0261 | ACV28313.1 | GCN5-related N-acetyltransferase | 54 | 8 | – | 0 (0) | Protein detected; site not covered |
| Apre_0261 | ACV28313.1 | GCN5-related N-acetyltransferase | 57 | 8 | – | 1 (0) | Protein detected; site not covered |
| Apre_0285 | ACV28336.1 | NUDIX hydrolase | 30 | 8 | YGWEGLTFPGGK | 1 (0) | Trp peptide detected |
| Apre_0348 | ACV28397.1 | hypothetical protein | 285 | 6 | – | 1 (0) | Protein detected; site not covered |
| Apre_0349 | ACV28398.1 | protein of unknown function DUF87 | 248 | 6 | – | 1 (0) | Protein detected; site not covered |
| Apre_0368 | ACV28417.1 | selenide, water dikinase | 154 | 9 | EIWENNTPK | 0 (0) | Trp peptide detected |
| Apre_0412 | ACV28461.1 | hypothetical protein | 81 | 10 | – | 1 (0) | Protein detected; site not covered |
| Apre_0421 | ACV28470.1 | hypothetical protein | 13 | – | – | – | Protein not detected |
| Apre_0602 | ACV28644.1 | hypothetical protein | 16 | 6 | – | 1 (0) | Protein detected; site not covered |
| Apre_0616 | ACV28658.1 | cell divisionFtsK/SpoIIIE | 511 | 24 | – | 2 (0) | Protein detected; site not covered |
| Apre_0640 | ACV28679.1 | conserved hypothetical protein | 48 | – | – | – | Protein not detected |
| Apre_0644 | ACV28683.1 | histidine kinase | 246 | 14 | ISWEISK | 0 (0) | Trp peptide detected |
| Apre_0714 | ACV28747.1 | hypothetical protein | 134 | – | – | – | Protein not detected |
| Apre_0794 | ACV28822.1 | sucrose-6-phosphate hydrolase | 220 | 32 | – | 2 (0) | Protein detected; site not covered |
| Apre_0817 | ACV28845.1 | hypothetical protein | 87 | – | – | – | Protein not detected |
| Apre_0901 | ACV28929.1 | HAD-superfamily subfamily IB hydrolase, TIGR0149 | 157 | 13 | VIPMWDGNSK | 0 (0) | Trp peptide detected |
| Apre_1039 | ACV29067.1 | helicase domain protein | 58 | – | – | – | Protein not detected |
| Apre_1076 | ACV29103.1 | hypothetical protein | 480 | 9 | VIEGWAR | 0 (0) | Trp peptide detected |
| Apre_1096 | ACV29123.1 | transcriptional regulator, TetR family | 191 | 6 | – | 0 (0) | Protein detected; site not covered |
| Apre_1113 | ACV29140.1 | major facilitator superfamily MFS_1 | 302 | – | – | – | Protein not detected |
| Apre_1139 | ACV29165.1 | conserved hypothetical protein | 202 | 36 | – | 1 (0) | Protein detected; site not covered |
| Apre_1177 | ACV29202.1 | oxygen-independent coproporphyrinogen III oxidas | 185 | 4 | – | 2 (0) | Protein detected; site not covered |
| Apre_1245 | ACV29269.1 | sodium:neurotransmitter symporter | 25 | 4 | – | 1 (0) | Protein detected; site not covered |
| Apre_1349 | ACV29372.1 | hypothetical protein | 189 | 4 | – | 1 (0) | Protein detected; site not covered |
| Apre_1416 | ACV29439.1 | tRNA pseudouridine synthase A | 242 | – | – | – | Protein not detected |
| Apre_1447 | ACV29470.1 | hypothetical protein | 221 | 6 | – | 1 (0) | Protein detected; site not covered |
| Apre_1454 | ACV29475.1 | Cna B domain protein | 4696 | 76 | – | 1 (0) | Protein detected; site not covered |
| Apre_1454 | ACV29475.1 | Cna B domain protein | 4483 | 76 | – | 1 (0) | Protein detected; site not covered |
| Apre_1496 | ACV29517.1 | UvrD/REP helicase | 415 | 24 | – | 1 (0) | Protein detected; site not covered |
| Apre_1579 | ACV29600.1 | Cna B domain protein | 3606 | 406 | – | 1 (0) | Protein detected; site not covered |
| Apre_1653 | ACV29674.1 | hypothetical protein | 53 | 6 | – | 1 (0) | Protein detected; site not covered |

**Supplementary Table S5.** The six Trp-containing precursors and their protein-level support. Intensities are DIA-NN precursor quantities. “Isobaric tryptic peptides” counts tryptic peptides (≤2 missed cleavages) of the reference proteome whose 2+ m/z lies within 10 ppm of the Trp peptide; mass alone does not discriminate them, so identification rests on DIA-NN fragment scoring.

| Locus | Site | Peptide | z | m/z (theor.) | Intensity (a.u.) | Protein: precursors / peptides / coverage | Rank in protein | Run (%) | Isobaric tryptic peptides (≤10 ppm) |
|---|---|---|---|---|---|---|---|---|---|
| Apre_0094 | Arg353 | AWEGSLR | 2 | 409.7114 | 9.51e+04 | 9 / 9 / 21.7% | 1 of 9 | 48 | 4 |
| Apre_0285 | Arg30 | YGWEGLTFPGGK | 2 | 656.3220 | 6.82e+03 | 8 / 8 / 55.1% | 8 of 8 | 4 | 6 |
| Apre_0368 | Arg154 | EIWENNTPK | 2 | 565.7775 | 4.83e+04 | 9 / 9 / 31.5% | 5 of 9 | 34 | 10 |
| Apre_0644 | Arg246 | ISWEISK | 2 | 431.7371 | 2.11e+05 | 14 / 14 / 31.2% | 2 of 14 | 62 | 20 |
| Apre_0901 | Arg157 | VIPMWDGNSK | 2 | 573.7842 | 4.13e+05 | 13 / 12 / 40.9% | 5 of 13 | 73 | 8 |
| Apre_1076 | Arg480 | VIEGWAR | 2 | 415.7296 | 4.96e+04 | 9 / 9 / 10.6% | 2 of 9 | 35 | 6 |

**Supplementary Table S6.**
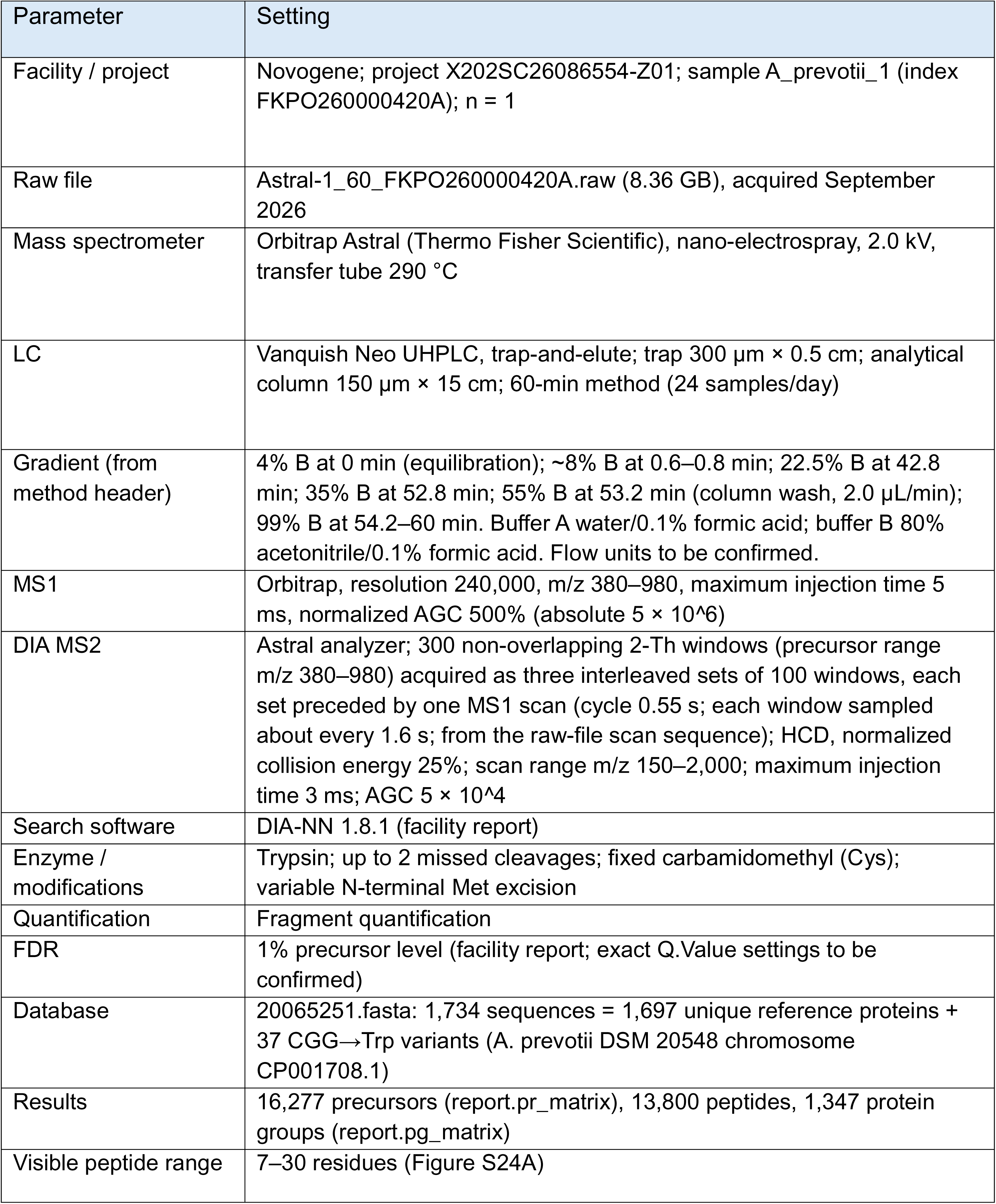
DIA acquisition and search parameters. Instrument and LC settings were read from the raw-file method header; search settings are from the facility report.

| Parameter | Setting |
|---|---|
| Facility / project | Novogene; project X202SC26086554-Z01; sample A_prevotii_1 (index FKPO260000420A); n = 1 |
| Raw file | Astral-1_60_FKPO260000420A.raw (8.36 GB), acquired September 2026 |
| Mass spectrometer | Orbitrap Astral (Thermo Fisher Scientific), nano-electrospray, 2.0 kV, transfer tube 290 °C |
| LC | Vanquish Neo UHPLC, trap-and-elute; trap 300 µm × 0.5 cm; analytical column 150 µm × 15 cm; 60-min method (24 samples/day) |
| Gradient (from method header) | 4% B at 0 min (equilibration); ~8% B at 0.6–0.8 min; 22.5% B at 42.8 min; 35% B at 52.8 min; 55% B at 53.2 min (column wash, 2.0 µL/min); 99% B at 54.2–60 min. Buffer A water/0.1% formic acid; buffer B 80% acetonitrile/0.1% formic acid. Flow units to be confirmed. |
| MS1 | Orbitrap, resolution 240,000, m/z 380–980, maximum injection time 5 ms, normalized AGC 500% (absolute 5 × 10^6) |
| DIA MS2 | Astral analyzer; 300 non-overlapping 2-Th windows (precursor range m/z 380–980) acquired as three interleaved sets of 100 windows, each set preceded by one MS1 scan (cycle 0.55 s; each window sampled about every 1.6 s; from the raw-file scan sequence); HCD, normalized collision energy 25%; scan range m/z 150–2,000; maximum injection time 3 ms; AGC 5 × 10^4 |
| Search software | DIA-NN 1.8.1 (facility report) |
| Enzyme / modifications | Trypsin; up to 2 missed cleavages; fixed carbamidomethyl (Cys); variable N-terminal Met excision |
| Quantification | Fragment quantification |
| FDR | 1% precursor level (facility report; exact Q.Value settings to be confirmed) |
| Database | 20065251.fasta: 1,734 sequences = 1,697 unique reference proteins + 37 CGG→Trp variants (A. prevotii DSM 20548 chromosome CP001708.1) |
| Results | 16,277 precursors (report.pr_matrix), 13,800 peptides, 1,347 protein groups (report.pg_matrix) |
| Visible peptide range | 7–30 residues (Figure S24A) |

**Supplementary Table S7.** Arg and Trp tRNA genes of *A. prevotii*. Anticodon and position 73 (the base preceding the 3′-terminal CCA) were read from global alignments to tRNA 36a; identity is to Apre_R0030. Sequences are in the Supplementary Data workbook.

| Locus | Annotation | Location | Length (nt) | Anticodon | Position 73 | Identity to Apre_R0030 (%) |
|---|---|---|---|---|---|---|
| Apre_R0019 | tRNA-Arg | 242212..242288 | 77 | UCU | G | 57 |
| Apre_R0025 | tRNA-Trp | 243030..243105 | 76 | CCA | G | 64 |
| Apre_R0030 | tRNA-Arg | 583059..583132 | 74 | CCG | G | 100 |
| Apre_R0043 | tRNA-Arg | Complement (1382505..1382581) | 77 | CCU | A | 60 |
| Apre_R0046 | tRNA-Arg | Complement (1425651..1425727) | 77 | ACG | G | 62 |

**Supplementary Table S8.** Raw-data inspection of the six Trp-form peptides. The cluster is the window of five consecutive MS2 scans with the highest fragment-ion score (Materials and methods); its time was inferred from the raw file because DIA-NN retention times were not available. “Ions matched” counts b and y ions (1+ and 2+; b1 and y1 excluded) within 5 ppm in the highest-scoring scan, of those that fall in m/z 150– 1, 500; the number in parentheses contains the Trp residue. Rank is the intensity rank of the ion among all peaks of the scan. Family-wise P: fraction of 198 decoy fragment lists whose best window anywhere in the run scored at least as high (minimum possible 0.005). MS1 M0 is the summed intensity of five MS1 scans at the monoisotopic m/z (±10 ppm); isotope ratios are observed (theoretical from the elemental composition). “Other clusters”: further windows with family-wise P ≤ 0.05; a similar fragment pattern at several times, as for ISWEISK, may reflect isomeric or co-isolated peptides that share fragment ions, which this analysis cannot resolve. DIA-NN percentile: rank of the precursor quantity in the run (Table S5).

| Peptide | Locus (site) | Cluster time (min) | Ions matched / possible (Trp-containing) | Most intense matched ion (rank) | Family-wise P | MS1 M0 (a.u.) | M+1/M0 obs. (exp.) | M+2/M0 obs. (exp.) | Other clusters (min) | DIA-NN run percentile |
|---|---|---|---|---|---|---|---|---|---|---|
| ISWEISK | Apre_0644 (W246) | 16.8 | 7 / 18 (6) | y6 (1) | 0.005 | 6.4e+07 | 0.39 (0.48) | 0.08 (0.14) | 27.8, 8.9, 10.8 | 62 |
| VIPMWDGNSK | Apre_0901 (W157) | 30.9 | 8 / 30 (4) | y8++ (3) | 0.005 | 1.3e+06 | 0.82 (0.62) | 0.35 (0.27) | 31.3, 52.5 | 73 |
| EIWENNTPK | Apre_0368 (W154) | 17.6 | 6 / 26 (2) | y7 (15) | 0.015 | 3.5e+05 | 3.24 (0.60) | 1.19 (0.21) | – | 34 |
| VIEGWAR | Apre_1076 (W480) | 15.0 | 6 / 18 (3) | b5 (37) | 0.005 | – | not detected | – | 10.1, 8.3, 18.4 | 35 |
| AWEGSLR | Apre_0094 (W353) | 15.5 | 5 / 18 (2) | y5 (15) | 0.005 | 2.8e+06 | 0.08 (0.44) | 0.08 (0.12) | 12.9 | 48 |
| YGWEGLTFPGGK | Apre_0285 (W30) | 50.0 | 6 / 37 (2) | y4 (41) | 0.035 | 2.5e+05 | 0.60 (0.75) | 0.19 (0.31) | 53.7 | 4 |

## Supplementary sequences of selected plasmids

### Complete plasmid sequence of pJL1-sfGFP

sfGFP under control of T7 promoter and RBS, and T7 terminator.

>pJL1-sfGFP (2480 bp)

gatgccacgattggcaaactgacgctgaaatttatctgcaccaccggcaaactgccggtgccgtggccgacgctggtgac caccctgacctatggcgttcagtgttttagtcgctatccggatcacatgaaacgtcacgatttctttaaatctgcaatgccgga aggctatgtgcaggaacgtacgattagctttaaagatgatggcaaatataaaacgcgcgccgttgtgaaatttgaaggcg ataccctggtgaaccgcattgaactgaaaggcacggattttaaagaagatggcaatatcctgggccataaactggaatac aactttaatagccataatgtttatattacggcggataaacagaaaaatggcatcaaagcgaattttaccgttcgccataacgt tgaagatggcagtgtgcagctggcagatcattatcagcagaataccccgattggtgatggtccggtgctgctgccggataa tcattatctgagcacgcagaccgttctgtctaaagatccgaacgaaaaaggcacgcgggaccacatggttctgcacgaat atgtgaatgcggcaggtattacgtggagccatccgcagttcgaaaaataaccggctgctaacaaagcccgaaaggaag ctgagttggctgctgccaccgctgagcaataactagcataaccccttggggcctctaaacgggtcttgaggggttttttgctg aaagccaattctgattagaaaaactcatcgagcatcaaatgaaactgcaatttattcatatcaggattatcaataccatattttt gaaaaagccgtttctgtaatgaaggagaaaactcaccgaggcagttccataggatggcaagatcctggtatcggtctgcg attccgactcgtccaacatcaatacaacctattaatttcccctcgtcaaaaataaggttatcaagtgagaaatcaccatgagt gacgactgaatccggtgagaatggcaaaagcttatgcatttctttccagacttgttcaacaggccagccattacgctcgtcat caaaatcactcgcatcaaccaaaccgttattcattcgtgattgcgcctgagcgagacgaaatacgcgatcgctgttaaaag gacaattacaaacaggaatcgaatgcaaccggcgcaggaacactgccagcgcatcaacaatattttcacctgaatcag gatattcttctaatacctggaatgctgttttcccggggatcgcagtggtgagtaaccatgcatcatcaggagtacggataaaa tgcttgatggtcggaagaggcataaattccgtcagccagtttagtctgaccatctcatctgtaacatcattggcaacgctacct ttgccatgtttcagaaacaactctggcgcatcgggcttcccatacaatcgatagattgtcgcacctgattgcccgacattatcg cgagcccatttatacccatataaatcagcatccatgttggaatttaatcgcggcttcgagcaagacgtttcccgttgaatatgg ctcataacaccccttgtattactgtttatgtaagcagacagttttattgttcatgatgatatatttttatcttgtgcaatgtaacatcag agattttgagacacaacgtgagatcaaaggatcttcttgagatcctttttttctgcgcgtaatctgctgcttgcaaacaaaaaa accaccgctaccagcggtggtttgtttgccggatcaagagctaccaactctttttccgaaggtaactggcttcagcagagcg cagataccaaatactgttcttctagtgtagccgtagttaggccaccacttcaagaactctgtagcaccgcctacatacctcgc tctgctaatcctgttaccagtggctgctgccagtggcgataagtcgtgtcttaccgggttggactcaagacgatagttaccgg ataaggcgcagcggtcgggctgaacggggggttcgtgcacacagcccagcttggagcgaacgacctacaccgaactg agatacctacagcgtgagctatgagaaagcgccacgcttcccgaagggagaaaggcggacaggtatccggtaagcg gcagggtcggaacaggagagcgcacgagggagcttccagggggaaacgcctggtatctttatagtcctgtcgggtttcg ccacctctgacttgagcgtcgatttttgtgatgctcgtcaggggggcggagcctatggaaaaacgccagcaacgcgatcc cgcgaaattaatacgactcactatagggagaccacaacggtttccctctagaaataattttgtttaactttaagaaggagata tacaatgagcaaaggtgaagaactgtttaccggcgttgtgccgattctggtggaactggatggcgatgtgaacggtcacaa attcagcgtgcgtggtgaaggtgaaggc

### Complete plasmid sequence of pJL1-Elastin-sfGFP

Elastin-sfGFP under control of T7 promoter, RBS, and T7 terminator.

>pJL1-Elastin-sfGFP (2566 bp)

gatgccacgattggcaaactgacgctgaaatttatctgcaccaccggcaaactgccggtgccgtggccgacgctggtgac caccctgacctatggcgttcagtgttttagtcgctatccggatcacatgaaacgtcacgatttctttaaatctgcaatgccgga aggctatgtgcaggaacgtacgattagctttaaagatgatggcaaatataaaacgcgcgccgttgtgaaatttgaaggcg ataccctggtgaaccgcattgaactgaaaggcacggattttaaagaagatggcaatatcctgggccataaactggaatac aactttaatagccataatgtttatattacggcggataaacagaaaaatggcatcaaagcgaattttaccgttcgccataacgt tgaagatggcagtgtgcagctggcagatcattatcagcagaataccccgattggtgatggtccggtgctgctgccggataa tcattatctgagcacgcagaccgttctgtctaaagatccgaacgaaaaaggcacgcgggaccacatggttctgcacgaat atgtgaatgcggcaggtattacgtggagccatccgcagttcgaaaaataagtcgaccggctgctaacaaacccgaaag gaagctgagttggctgctgccaccgctgagcaataactagcataaccccttggggcctctaaacgggtcttgaggggtttttt gctgaaagccaattctgattagaaaaactcatcgagcatcaaatgaaactgcaatttattcatatcaggattatcaataccat atttttgaaaaagccgtttctgtaatgaaggagaaaactcaccgaggcagttccataggatggcaagatcctggtatcggtc tgcgattccgactcgtccaacatcaatacaacctattaatttcccctcgtcaaaaataaggttatcaagtgagaaatcaccat gagtgacgactgaatccggtgagaatggcaaaagcttatgcatttctttccagacttgttcaacaggccagccattacgctc gtcatcaaaatcactcgcatcaaccaaaccgttattcattcgtgattgcgcctgagcgagacgaaatacgcgatcgctgtta aaaggacaattacaaacaggaatcgaatgcaaccggcgcaggaacactgccagcgcatcaacaatattttcacctgaa tcaggatattcttctaatacctggaatgctgttttcccggggatcgcagtggtgagtaaccatgcatcatcaggagtacggat aaaatgcttgatggtcggaagaggcataaattccgtcagccagtttagtctgaccatctcatctgtaacatcattggcaacgc tacctttgccatgtttcagaaacaactctggcgcatcgggcttcccatacaatcgatagattgtcgcacctgattgcccgacat tatcgcgagcccatttatacccatataaatcagcatccatgttggaatttaatcgcggcttcgagcaagacgtttcccgttgaa tatggctcataacaccccttgtattactgtttatgtaagcagacagttttattgttcatgatgatatatttttatcttgtgcaatgtaac atcagagattttgagacacaacgtgagatcaaaggatcttcttgagatcctttttttctgcgcgtaatctgctgcttgcaaacaa aaaaaccaccgctaccagcggtggtttgtttgccggatcaagagctaccaactctttttccgaaggtaactggcttcagcag agcgcagataccaaatactgttcttctagtgtagccgtagttaggccaccacttcaagaactctgtagcaccgcctacatac ctcgctctgctaatcctgttaccagtggctgctgccagtggcgataagtcgtgtcttaccgggttggactcaagacgatagtta ccggataaggcgcagcggtcgggctgaacggggggttcgtgcacacagcccagcttggagcgaacgacctacaccg aactgagatacctacagcgtgagctatgagaaagcgccacgcttcccgaagggagaaaggcggacaggtatccggta agcggcagggtcggaacaggagagcgcacgagggagcttccagggggaaacgcctggtatctttatagtcctgtcggg tttcgccacctctgacttgagcgtcgatttttgtgatgctcgtcaggggggcggagcctatggaaaaacgccagcaacgcg atcccgcgaaattaatacgactcactatagggagaccacaacggtttccctctagaaataattttgtttaactttaagaagga gatatacatatgagcaaaggcccaggcaaagttccgggcgccggtgttccggggtggggtgttccaggtgttggcaaag gtggtggcactatgagcaaaggtgaagaactgtttaccggcgttgtgccgattctggtggaactggatggcgatgtgaacg gtcacaaattcagcgtgcgtggtgaaggtgaaggc

### Complete plasmid sequence of pJL1-Firefly_luc-Strep tag

Firefly luc-Strep tag under control of T7 promoter, RBS, and T7 terminator.

>pJL1-Firefly luc-GGGS-Strep tag (3446 bp) ttgagatcctttttttctgcgcgtaatctgctgcttgcaaacaaaaaaaccaccgctaccagcggtggtttgtttgccggatcaa gagctaccaactctttttccgaaggtaactggcttcagcagagcgcagataccaaatactgttcttctagtgtagccgtagtta ggccaccacttcaagaactctgtagcaccgcctacatacctcgctctgctaatcctgttaccagtggctgctgccagtggcg ataagtcgtgtcttaccgggttggactcaagacgatagttaccggataaggcgcagcggtcgggctgaacggggggttcg tgcacacagcccagcttggagcgaacgacctacaccgaactgagatacctacagcgtgagctatgagaaagcgccac gcttcccgaagggagaaaggcggacaggtatccggtaagcggcagggtcggaacaggagagcgcacgagggagct tccagggggaaacgcctggtatctttatagtcctgtcgggtttcgccacctctgacttgagcgtcgatttttgtgatgctcgtcag gggggcggagcctatggaaaaacgccagcaacgcgatcccgcgaaattaatacgactcactatagggagaccacaa cggtttccctctagaaataattttgtttaactttaagaaggagatatacaatggaggatgctaaaaatattaaaaagggccca gccccattttatccccttgaagacggcactgcaggtgaacagctgcacaaagctatgaaacgttatgcgttagtcccgggt

acgatcgcgtttactgatgcgcatatcgaagtaaatatcacatacgcggaatattttgaaatgtcggttcgtctcgccgaggc aatgaaaagatatggtctgaacaccaaccacagaattgtggtatgctcagaaaactccctccaattctttatgcctgtgctgg gagcgctttttatcggtgtcgccgtcgctcctgcgaacgacatctataacgaacgtgaactcctgaactctatgaatatctccc agcccacggttgtctttgtgtctaaaaaaggtctgcaaaaaatattaaatgtacagaaaaagcttccgattattcaaaaaatc atcattatggattcgaagaccgactatcaggggttccaatccatgtatacattcgttacttcacatcttcctccgggttttaacga atatgacttcgttccggaaagcttcgatcgcgataaaacaatagccttaataatgaactcgagtgggagcaccggcttacct aaaggagttgcgctgccgcaccgtacagcgtgcgtccgtttttcacatgcccgtgaccccatttttggtaaccagattatccct gatacggccattctctcggttgttccattccatcatgggtttggcatgttcacgacattgggttatctgatttgcggatttcgtgtgg tgctgatgtatcgctttgaagaggaactttttttgaggtcgttgcaagactataagatacaatctgcgctcctggttccaaccttg ttttccttttttgcaaaatcgacactgatcgataaatacgatctgtcaaatctgcatgagatagctagcggtggtgcaccgttga gcaaagaagttggcgaggccgttgccaagagattccatctgcctggtattcgtcagggatacggtctcacggaaaccacg tcagctatcctgattactcccgaaggagatgataagcccggcgcagttgggaaagtcgttccgttttttgaagccaaagttgt tgatctggatacgggaaagacactgggggttaaccagcgtggcgaactttgcgttcgagggccaatgattatgtcgggtta cgtgaataatcctgaggcaaccaacgccttgatcgataaagacgggtggctgcattctggtgatattgcttattgggatgaa gatgagcattttttcatagttgaccgcctgaaatcgttaatcaaatataaaggatatcaggtagcaccagcggagctggagt ctattttacttcagcacccgaatatcttcgatgcaggcgtcgcgggtctccccgatgacgacgcgggggaactcccagcgg ccgttgtcgtcttggagcatggcaaaaccatgacagagaaagaaattgtagattatgtggcttcacaggtgaccacggca aagaaattacgagggggcgttgtctttgtggacgaagttcctaaaggactgacgggtaaactcgacgcccgtaagatccg tgagattttaattaaagcaaaaaagggcgggaagagcaaactgggtggtggtagctggagccatccgcagttcgaaaa ataaccggctgctaacaaagcccgaaaggaagctgagttggctgctgccaccgctgagcaataactagcataacccctt ggggcctctaaacgggtcttgaggggttttttgctgaaagccaattctgattagaaaaactcatcgagcatcaaatgaaact gcaatttattcatatcaggattatcaataccatatttttgaaaaagccgtttctgtaatgaaggagaaaactcaccgaggcagt tccataggatggcaagatcctggtatcggtctgcgattccgactcgtccaacatcaatacaacctattaatttcccctcgtcaa aaataaggttatcaagtgagaaatcaccatgagtgacgactgaatccggtgagaatggcaaaagcttatgcatttctttcca gacttgttcaacaggccagccattacgctcgtcatcaaaatcactcgcatcaaccaaaccgttattcattcgtgattgcgcct gagcgagacgaaatacgcgatcgctgttaaaaggacaattacaaacaggaatcgaatgcaaccggcgcaggaacac tgccagcgcatcaacaatattttcacctgaatcaggatattcttctaatacctggaatgctgttttcccggggatcgcagtggtg agtaaccatgcatcatcaggagtacggataaaatgcttgatggtcggaagaggcataaattccgtcagccagtttagtctg accatctcatctgtaacatcattggcaacgctacctttgccatgtttcagaaacaactctggcgcatcgggcttcccatacaat cgatagattgtcgcacctgattgcccgacattatcgcgagcccatttatacccatataaatcagcatccatgttggaatttaat cgcggcttcgagcaagacgtttcccgttgaatatggctcataacaccccttgtattactgtttatgtaagcagacagttttattgtt catgatgatatatttttatcttgtgcaatgtaacatcagagattttgagacacaacgtgagatcaaaggatcttc

### Complete plasmid sequence of pJL1-NanoLuc-Strep tag

NanoLuc-Strep tag under control of T7 promoter, RBS, and T7 terminator.

>pJL1-NanoLuc-GGGS-Strep tag (2309 bp) ccggctgctaacaaagcccgaaaggaagctgagttggctgctgccaccgctgagcaataactagcataaccccttggg gcctctaaacgggtcttgaggggttttttgctgaaagccaattctgattagaaaaactcatcgagcatcaaatgaaactgca atttattcatatcaggattatcaataccatatttttgaaaaagccgtttctgtaatgaaggagaaaactcaccgaggcagttcc ataggatggcaagatcctggtatcggtctgcgattccgactcgtccaacatcaatacaacctattaatttcccctcgtcaaaa ataaggttatcaagtgagaaatcaccatgagtgacgactgaatccggtgagaatggcaaaagcttatgcatttctttccaga cttgttcaacaggccagccattacgctcgtcatcaaaatcactcgcatcaaccaaaccgttattcattcgtgattgcgcctga gcgagacgaaatacgcgatcgctgttaaaaggacaattacaaacaggaatcgaatgcaaccggcgcaggaacactg ccagcgcatcaacaatattttcacctgaatcaggatattcttctaatacctggaatgctgttttcccggggatcgcagtggtga gtaaccatgcatcatcaggagtacggataaaatgcttgatggtcggaagaggcataaattccgtcagccagtttagtctga ccatctcatctgtaacatcattggcaacgctacctttgccatgtttcagaaacaactctggcgcatcgggcttcccatacaatc

gatagattgtcgcacctgattgcccgacattatcgcgagcccatttatacccatataaatcagcatccatgttggaatttaatc gcggcttcgagcaagacgtttcccgttgaatatggctcataacaccccttgtattactgtttatgtaagcagacagttttattgttc atgatgatatatttttatcttgtgcaatgtaacatcagagattttgagacacaacgtgagatcaaaggatcttcttgagatcctttt tttctgcgcgtaatctgctgcttgcaaacaaaaaaaccaccgctaccagcggtggtttgtttgccggatcaagagctaccaa ctctttttccgaaggtaactggcttcagcagagcgcagataccaaatactgttcttctagtgtagccgtagttaggccaccact tcaagaactctgtagcaccgcctacatacctcgctctgctaatcctgttaccagtggctgctgccagtggcgataagtcgtgt cttaccgggttggactcaagacgatagttaccggataaggcgcagcggtcgggctgaacggggggttcgtgcacacagc ccagcttggagcgaacgacctacaccgaactgagatacctacagcgtgagctatgagaaagcgccacgcttcccgaag ggagaaaggcggacaggtatccggtaagcggcagggtcggaacaggagagcgcacgagggagcttccaggggga aacgcctggtatctttatagtcctgtcgggtttcgccacctctgacttgagcgtcgatttttgtgatgctcgtcaggggggcgga gcctatggaaaaacgccagcaacgcgatcccgcgaaattaatacgactcactatagggagaccacaacggtttccctct agaaataattttgtttaactttaagaaggagatatacaatggtcttcacactcgaagatttcgttggggactggcgacagaca gccggctacaacctggaccaagtccttgaacagggaggtgtgtccagtttgtttcagaatctcggggtgtccgtaactccga tccaaaggattgtcctgagcggtgaaaatgggctgaagatcgacatccatgtcatcatcccgtatgaaggtctgagcggc gaccaaatgggccagatcgaaaaaatttttaaggtggtgtaccctgtggacgatcatcactttaaggtgatcctgcactatg gcacactggtaatcgacggggttacgccgaacatgatcgactatttcggacgtccgtatgaaggcatcgccgtgttcgacg gcaaaaagatcactgtaacagggaccctgtggaacggcaacaaaattatcgacgagcgcctgatcaaccccgacggc tccctgctgttccgagtaaccatcaacggagtgaccggctggcgactgtgcgaacgcattctggcgggtggtggtagctgg agccatccgcagttcgaaaaataa

## Notes

https://www.ebi.ac.uk/pride/archive/projects/PXD086557/private

### Competing Interest Statement

The authors have declared no competing interest.

### Summary of Updates

Abstract: minor revision Author affiliations updated

## REFERENCES

1. Nirenberg, M., Caskey, T., Marshall, R., et al. (1966) The RNA code and protein synthesis. Cold Spring Harb Symp Quant Biol, 31, 11–24.

2. Lukeš, J., Eliáš, M., Kachale, A., et al. (2025) Natural and artificial variations of the standard genetic code. Curr Biol, 35, R1104–R1126.

3. Schultz, D.W. and Yarus, M. (1994) Transfer RNA mutation and the malleability of the genetic code. J Mol Biol, 235, 1377–1380.

4. Elzanowski, A. and Ostell, J. (2024) The Genetic Codes. National Center for Biotechnology Information.

5. Yamao, F., Muto, A., Kawauchi, Y., et al. (1985) UGA is read as tryptophan in Mycoplasma capricolum. Proc Natl Acad Sci U S A, 82, 2306–2309.

6. Oba, T., Andachi, Y., Muto, A., et al. (1991) Translation in vitro of codon UGA as tryptophan in Mycoplasma capricolum. Biochimie, 73, 1109–1112.

7. Campbell, J.H., O’Donoghue, P., Campbell, A.G., et al. (2013) UGA is an additional glycine codon in uncultured SR1 bacteria from the human microbiota. Proc Natl Acad Sci U S A, 110, 5540–5545.

8. Borges, A.L., Lou, Y.C., Sachdeva, R., et al. (2022) Widespread stop-codon recoding in bacteriophages may regulate translation of lytic genes. Nat Microbiol, 7, 918– 927.

9. Ivanova, N.N., Schwientek, P., Tripp, H.J., et al. (2014) Stop codon reassignments in the wild. Science, 344, 909–913.

10. Shulgina, Y. and Eddy, S.R. (2021) A computational screen for alternative genetic codes in over 250, 000 genomes. Elife, 10, e71402.

11. Silverman, A.D., Karim, A.S. and Jewett, M.C. (2020) Cell-free gene expression: an expanded repertoire of applications. Nat Rev Genet, 21, 151–170.

12. Hunt, A.C., Rasor, B.J., Seki, K., et al. (2024) Cell-Free Gene Expression: Methods and Applications. Chem Rev, 125, 91–149.

13. Nawrocki, E.P., Petrov, A.I. and Williams, K.P. (2025) Expansion of the tmRNA sequence database and new tools for search and visualization. NAR Genom Bioinform, 7, lqaf019.

14. Chaumeil, P.-A., Mussig, A.J., Hugenholtz, P., et al. (2022) GTDB-Tk v2: memory friendly classification with the genome taxonomy database. Bioinformatics, 38, 5315–5316.

15. Parks, D.H., Chuvochina, M., Rinke, C., et al. (2022) GTDB: an ongoing census of bacterial and archaeal diversity through a phylogenetically consistent, rank normalized and complete genome-based taxonomy. Nucleic Acids Res, 50, D785–D794.

16. Seki, K., Nguyen, M.T.A., Penev, P.I., et al. (2026) Engineered Orthogonal Translation Systems from Metagenomic Libraries Expand the Genetic Code. ACS Synth Biol, 15, 4157–4171.

17. Des Soye, B.J., Gerbasi, V.R., Thomas, P.M., et al. (2019) A Highly Productive, One-Pot Cell-Free Protein Synthesis Platform Based on Genomically Recoded *Escherichia coli*. Cell Chemical Biology, 26, 1743–1754.e9.

18. Jewett, M.C. and Swartz, J.R. (2004) Mimicking the Escherichia coli cytoplasmic environment activates long-lived and efficient cell-free protein synthesis. Biotechnol Bioeng, 86, 19–26.

19. Jewett, M.C., Calhoun, K.A., Voloshin, A., et al. (2008) An integrated cell-free metabolic platform for protein production and synthetic biology. Mol Syst Biol, 4, MSB200857.

20. Schneider, C.A., Rasband, W.S. and Eliceiri, K.W. (2012) NIH Image to ImageJ: 25 years of image analysis. Nat Methods, 9, 671–675.

21. Demichev, V., Messner, C.B., Vernardis, S.I., et al. (2020) DIA-NN: neural networks and interference correction enable deep proteome coverage in high throughput. Nat Methods, 17, 41–44.

22. Hulstaert, N., Shofstahl, J., Sachsenberg, T., et al. (2020) ThermoRawFileParser: Modular, Scalable, and Cross-Platform RAW File Conversion. J Proteome Res, 19, 537–542.

23. Katoh, K. and Standley, D.M. (2013) MAFFT multiple sequence alignment software version 7: improvements in performance and usability. Mol Biol Evol, 30, 772– 780.

24. Kozlov, A.M., Darriba, D., Flouri, T., et al. (2019) RAxML-NG: a fast, scalable and user-friendly tool for maximum likelihood phylogenetic inference. Bioinformatics, 35, 4453–4455.

25. Le, S.Q. and Gascuel, O. (2008) An improved general amino acid replacement matrix. Mol Biol Evol, 25, 1307–1320.

26. Shulgina, Y. and Eddy, S.R. (2023) Codetta: predicting the genetic code from nucleotide sequence. Bioinformatics, 39, btac802.

27. Nyerges, A., Vinke, S., Flynn, R., et al. (2023) A swapped genetic code prevents viral infections and gene transfer. Nature, 615, 720–727.

28. Johnson, D.B.F., Wang, C., Xu, J., et al. (2012) Release factor one is nonessential in Escherichia coli. ACS Chem Biol, 7, 1337–1344.

29. Jørgensen, F., Adamski, F.M., Tate, W.P., et al. (1993) Release factor-dependent false stops are infrequent in Escherichia coli. J Mol Biol, 230, 41–50.

30. Heim, R., Prasher, D.C. and Tsien, R.Y. (1994) Wavelength mutations and posttranslational autoxidation of green fluorescent protein. Proc Natl Acad Sci U S A, 91, 12501–12504.

31. Heim, R. and Tsien, R.Y. (1996) Engineering green fluorescent protein for improved brightness, longer wavelengths and fluorescence resonance energy transfer. Curr Biol, 6, 178–182.

32. Pédelacq, J.-D., Cabantous, S., Tran, T., et al. (2006) Engineering and characterization of a superfolder green fluorescent protein. Nat Biotechnol, 24, 79–88.

33. Reid, B.G. and Flynn, G.C. (1997) Chromophore formation in green fluorescent protein. Biochemistry, 36, 6786–6791.

34. Hall, M.P., Unch, J., Binkowski, B.F., et al. (2012) Engineered luciferase reporter from a deep sea shrimp utilizing a novel imidazopyrazinone substrate. ACS Chem Biol, 7, 1848–1857.

35. Branchini, B.R., Ablamsky, D.M., Murtiashaw, M.H., et al. (2007) Thermostable red and green light-producing firefly luciferase mutants for bioluminescent reporter applications. Anal Biochem, 361, 253–262.

36. Conti, E., Franks, N.P. and Brick, P. (1996) Crystal structure of firefly luciferase throws light on a superfamily of adenylate-forming enzymes. Structure, 4, 287–298.

37. Tomabechi, Y., Hosoya, T., Ehara, H., et al. (2016) Crystal structure of nanoKAZ: The mutated 19 kDa component of Oplophorus luciferase catalyzing the bioluminescent reaction with coelenterazine. Biochem Biophys Res Commun, 470, 88–93.

38. Dementieva, E.I., Fedorchuk, E.A., Brovko, L.Y., et al. (2000) Fluorescent properties of firefly luciferases and their complexes with luciferin. Biosci Rep, 20, 21–30.

39. Dong, H., Nilsson, L. and Kurland, C.G. (1996) Co-variation of tRNA abundance and codon usage in Escherichia coli at different growth rates. J Mol Biol, 260, 649– 663.

40. Wei, Y., Silke, J.R. and Xia, X. (2019) An improved estimation of tRNA expression to better elucidate the coevolution between tRNA abundance and codon usage in bacteria. Sci Rep, 9, 3184.

41. Kramer, E.B. and Farabaugh, P.J. (2007) The frequency of translational misreading errors in E. coli is largely determined by tRNA competition. RNA, 13, 87–96.

42. Elf, J., Nilsson, D., Tenson, T., et al. (2003) Selective charging of tRNA isoacceptors explains patterns of codon usage. Science, 300, 1718–1722.

43. Chica, R.A., Moore, M.M., Allen, B.D., et al. (2010) Generation of longer emission wavelength red fluorescent proteins using computationally designed libraries. Proceedings of the National Academy of Sciences, 107, 20257–20262.

44. Himeno, H., Hasegawa, T., Asahara, H., et al. (1991) Identity determinants of E. coli tryptophan tRNA. Nucleic Acids Res, 19, 6379–6382.

45. Giegé, R., Sissler, M. and Florentz, C. (1998) Universal rules and idiosyncratic features in tRNA identity. Nucleic Acids Res, 26, 5017–5035.

46. Mukai, T., Yamaguchi, A., Ohtake, K., et al. (2015) Reassignment of a rare sense codon to a non-canonical amino acid in Escherichia coli. Nucleic Acids Res, 43, 8111–8122.

47. Schwark, D.G., Schmitt, M.A., Biddle, W., et al. (2020) The Influence of Competing tRNA Abundance on Translation: Quantifying the Efficiency of Sense Codon Reassignment at Rarely Used Codons. ChemBioChem, 21, 2274–2286.

48. Robertson, W.E., Funke, L.F.H., de la Torre, D., et al. (2021) Sense codon reassignment enables viral resistance and encoded polymer synthesis. Science, 372, 1057–1062.

49. Cui, Z., Wu, Y., Mureev, S., et al. (2018) Oligonucleotide-mediated tRNA sequestration enables one-pot sense codon reassignment in vitro. Nucleic Acids Res, 46, 6387–6400.

50. Krishnakumar, R., Prat, L., Aerni, H.-R., et al. (2013) Transfer RNA misidentification scrambles sense codon recoding. Chembiochem, 14, 1967–1972.

51. Shimizu, Y., Inoue, A., Tomari, Y., et al. (2001) Cell-free translation reconstituted with purified components. Nat Biotechnol, 19, 751–755.

52. Miyachi, R. and Ichihashi, N. (2026) Experimental verification of the error minimization theory using non-standard genetic codes constructed in vitro. Elife, 15, RP111164.

53. Grome, M.W., Nguyen, M.T.A., Moonan, D.W., et al. (2025) Engineering a genomically recoded organism with one stop codon. Nature, 639, 512–521.

54. Isaacs, F.J., Carr, P.A., Wang, H.H., et al. (2011) Precise manipulation of chromosomes in vivo enables genome-wide codon replacement. Science, 333, 348–353.

55. Fredens, J., Wang, K., de la Torre, D., et al. (2019) Total synthesis of Escherichia coli with a recoded genome. Nature, 569, 514–518.

56. Robertson, W.E., Rehm, F.B.H., Spinck, M., et al. (2025) Escherichia coli with a 57-codon genetic code. Science, 390, eady4368.

## Supplementary References

1. Elzanowski, A. and Ostell, J. The Genetic Codes.

2. Barrell, B.G., Bankier, A.T. and Drouin, J. (1979) A different genetic code in human mitochondria. Nature, 282, 189–194.

3. Osawa, S., Collins, D., Ohama, T., et al. (1990) Evolution of the mitochondrial genetic code. III. Reassignment of CUN codons from leucine to threonine during evolution of yeast mitochondria. J Mol Evol, 30, 322–328.

4. Ling, J., Daoud, R., Lajoie, M.J., et al. (2014) Natural reassignment of CUU and CUA sense codons to alanine in Ashbya mitochondria. Nucleic Acids Res, 42, 499– 508.

5. Kawaguchi, Y., Honda, H., Taniguchi-Morimura, J., et al. (1989) The codon CUG is read as serine in an asporogenic yeast Candida cylindracea. Nature, 341, 164– 166.

6. Santos, M.A., Keith, G. and Tuite, M.F. (1993) Non-standard translational events in Candida albicans mediated by an unusual seryl-tRNA with a 5’-CAG-3’ (leucine) anticodon. EMBO J, 12, 607–616.

7. Mühlhausen, S., Findeisen, P., Plessmann, U., et al. (2016) A novel nuclear genetic code alteration in yeasts and the evolution of codon reassignment in eukaryotes. Genome Res, 26, 945–955.

8. Osawa, S., Ohama, T., Jukes, T.H., et al. (1989) Evolution of the mitochondrial genetic code. I. Origin of AGR serine and stop codons in metazoan mitochondria. J Mol Evol, 29, 202–207.

9. Yokobori, S., Ueda, T. and Watanabe, K. (1993) Codons AGA and AGG are read as glycine in ascidian mitochondria. J Mol Evol, 36, 1–8.

10. Abascal, F., Posada, D., Knight, R.D., et al. (2006) Parallel evolution of the genetic code in arthropod mitochondrial genomes. PLoS Biol, 4, e127.

11. Perseke, M., Hetmank, J., Bernt, M., et al. (2011) The enigmatic mitochondrial genome of Rhabdopleura compacta (Pterobranchia) reveals insights into selection of an efficient tRNA system and supports monophyly of Ambulacraria. BMC Evol Biol, 11, 134.

12. Noutahi, E., Calderon, V., Blanchette, M., et al. (2019) Rapid Genetic Code Evolution in Green Algal Mitochondrial Genomes. Mol Biol Evol, 36, 766–783.

13. Žihala, D. and Eliáš, M. (2019) Evolution and Unprecedented Variants of the Mitochondrial Genetic Code in a Lineage of Green Algae. Genome Biol Evol, 11, 2992–3007.

14. Shulgina, Y. and Eddy, S.R. (2021) A computational screen for alternative genetic codes in over 250, 000 genomes. Elife, 10, e71402.

